# Enhancing the biocorrosion resistance and biocompatibility of Aluminium substrates using Graphene Oxide-PEDOT:PSS Hybrid Coating

**DOI:** 10.1101/2025.09.29.679258

**Authors:** Sourya Subhra Nasker, Ragnhild Elizabeth Aune, Amogh K. Ravi, Tharangattu N. Narayanan, Sasmita Nayak

**Affiliations:** School of Biotechnology, Kalinga Institute of Industrial Technology (KIIT), Bhubaneswar, Odisha 751024, India; Department of Materials Science and Engineering, Norwegian University of Science and Technology (NTNU), 7491 Trondheim, Norway; Materials and Interface Engineering Laboratory, Tata Institute of Fundamental Research (TIFR), Hyderabad, Telangana 500046, India; Department of Microbiology, Kalinga Institute of Medical Sciences, Kalinga Institute of Industrial Technology Deemed to be University, Bhubaneswar, Odisha 751024, India

**Keywords:** Microbiologically Influenced Corrosion (MIC), Antifouling properties, GO-PEDOT:PSS Coatings, Aluminium Alloy 1050, Biocompatibility

## Abstract

Biofilm-associated infections on medical devices remain a major clinical challenge due to antimicrobial resistance, biofouling, and biocorrosion, compromising implant longevity and biocompatibility. Conventional biomedical devices utilizing stainless steel and titanium exhibit limited resistance to biofilm-prone physiological environments, necessitating development of next-generation implant materials. While alumina (Al_2_O_3_) is favored for its bio-inertness, it raises concerns like leaching and systemic toxicity. Aluminum alloy 1050 (AA1050) offers intrinsic corrosion resistance via a passive oxide layer under dry conditions but remains prone to microbiologically influenced corrosion (MIC) under humid conditions. In the recent years, graphene family of materials have emerged as promising surface coating for metals and alloys due to their strong barrier properties and antimicrobial efficacy. Most graphene derivatives like GO require relatively higher concentrations to achieve antimicrobial activity, however compromising biocompatibility, limiting *in vivo* human uses. This study addresses these caveats by exploring GO activity at lower concentrations (50–500 µg/ml) on AA1050 to achieve a balance between antimicrobial efficiency and cytocompatibility. Additionally, poly(3,4-ethylenedioxythiophene):poly(styrenesulfonate) (PEDOT:PSS, 1 µg/ml) was integrated with GO to form a hybrid coating on Al (Al_GO/P), to improve GO adhesion, and corrosion resistance. Biological evaluations against *Escherichia coli* (*E. coli*), *Staphylococcus aureus* (*S. aureus*), and *Candida albicans* (*C. albicans*) demonstrated superior antifouling and antimicrobial efficacy of Al_GO/P substrates compared to GO-coated Al (Al_GO) and bare Al surfaces. Corrosion rate, FESEM, ICP-MS, and cytotoxicity analyses further confirmed reduced biocorrosion, minimal ion leaching, along with enhanced biocompatibility of the hybrid coated surfaces. Al_GO/P containing GO at 100 and 250 µg/ml concentration achieved the optimal balance between antimicrobial activity and biocompatibility and can be used as *in vivo* implant materials. Hence, this technology can be implemented towards surface modification of biomedical devices to mitigate periprosthetic infections as well as support *ex vivo* applications requiring durable antimicrobial performance.

## 1. Introduction

Biofilms, structured microbial communities embedded within a self-produced extracellular matrix (ECM), pose a persistent clinical challenge due to their resilience to conventional therapies and impact on human health^1^. The ECM, also referred to as extracellular polymeric substances (EPS), not only supports nutrient absorption but also enhances microbial defense mechanisms against both host immune system responses and antimicrobial drugs^2^. Biofilm formation on medical devices such as implants, catheters, and ventilators often leads to chronic infections, which reduce the therapeutic effectiveness of interventions against medical device-associated infections (MDAIs). These infections significantly contribute to healthcare-associated infections (HAIs), a critical public health concern that affects approximately one in 31 hospitalized patients each day^3^. HAIs extend hospitalization periods, increase healthcare costs and are predominantly linked to invasive medical devices^3,4^. According to the European Center for Disease Prevention and Control (ECDC), the European Union and European Economic Area (EU/EEA) report over 4.3 million HAI cases annually, resulting in more than 90,000 deaths, exceeding the mortality burden of both influenza and tuberculosis in the region^5–7^. HAIs are also prevalent in the United States, where multidrug-resistant (MDR) infections contribute substantially to high morbidity and mortality rates^8–10^. In India, the burden is particularly severe, with HAI incidence ranging between 11% and 83% depending on the infection type, underscoring systemic vulnerabilities in the healthcare infrastructure^11–15^.

Previous research demonstrates that MDAIs are typically caused by biofilms comprising mixed microbial communities, including Gram-negative bacteria (e.g., *Escherichia coli* and *Pseudomonas aeruginosa)*, Gram-positive bacteria (e.g., *Staphylococcus aureus*, *Staphylococcus epidermidis*, and *Enterococcus faecalis*), and fungi, predominantly the *Candida* species^16–18^. Microbial adhesion leads to biofouling, which subsequently initiates biocorrosion through the formation of differential aeration cells. These cells accelerate electrochemical degradation and pitting corrosion^19^. Unlike uniform corrosion, pitting results in severe, localized damage that compromises structural integrity and releases cytotoxic metal ions into the surrounding tissues^20^. The complex interplay between biofouling and biocorrosion highlights a key limitation of current implant materials, emphasizing the urgent need for surfaces engineered to resist both microbial colonization and corrosion-induced degradation^21,22^.

These challenges reflect broader limitations to traditional implant materials, including stainless steel, aluminum, and synthetic polymers^23^. Their clinical use is often limited by cytotoxic effects, inflammatory host responses, and rapid degradation through either chemical or microbiologically influenced corrosion (MIC), often worsened by biofilm activity^24,25^. Metals and alloys such as 316L stainless steel, cobalt-chromium alloys, and titanium-based materials suffer from long-term performance issues arising from limited biocompatibility and corrosion-induced failure^26^. In chloride-rich physiological environments, mechanical wear and corrosion can release harmful ions such as cobalt (Co^2+^), nickel (Ni^2+^), aluminum (Al^3+^), and vanadium (V^3+^), raising significant concerns over systemic toxicity and neurotoxic effects^27,28^. Therefore, there is a clear need for advanced surface engineering strategies to develop materials that effectively mitigate biocorrosion and cytotoxicity without compromising performance.

Aluminum alloy 1050 (AA1050) is widely used in multiple sectors due to its excellent thermal and electrical conductivity^29,30^. As a commercially pure alloy with a minimum aluminum content of 99.5%, AA1050 is particularly attractive for various applications including packaging, transportation, and electronics^31^. Its relatively lower density further enhances its suitability for aerospace and automotive components^29^. While alumina (Al_2_O_3_) serves as a popular implant material due to its bio-inertness, other aluminum alloys like AA1050, AA1100, AA2024, and AA6061 have yet to find clinical use in implants^32–35^. However, Al_2_O_3_ has limitations, such as potential ion release from wear debris, which can trigger inflammation, bone resorption, and aseptic loosening at the implant-bone interface, potentially resulting in long-term Al^3+^ leaching^36–38^. Commonly used aluminum alloys such as AA1100, AA2024, and AA6061 are unsuitable for biomedical implants due to their composition (high Si content) and corrosion tendencies that promote tissue damage through ion release^33,39,40^. However, AA1050 demonstrates intrinsic corrosion resistance via the formation of a passive oxide layer under dry conditions, this protective barrier becomes compromised in humid or chloride-rich environents^41^. Under such conditions, AA1050 is vulnerable to MIC, as biofilms significantly alter the electrochemical landscape and promote corrosion ^42^.

To address such issues, recent advances in nanomaterials have facilitated the development of hybrid coatings that limit microbial adhesion and reduce the infection rates associated with devices^43^. Among these, the graphene family of materials (Gfam), especially graphene oxide (GO), has gained attention due to its high surface area, excellent barrier properties, strong adhesion to metals, and broad-spectrum antimicrobial properties^44–48^. Moreover, several studies confirms GO coating on metal surfaces such as stainless steel, titanium, aluminium, and copper, effectively mitigates biocorrosion and microbial adhesion^49–53^. However, these coatings typically employ high GO concentrations (0.5-10 mg/ml) to achieve enhanced antimicrobial activity, which subsequently limits biocompatibility; restricting their applications for *in vivo* human uses. Even though GO at a concentration of 1 mg/ml on AA1050 has been shown to form impermeable barriers against corrosive agents (water, oxygen, ions), it poses risk of systemic toxicity due to inflammatory responses ^54–57^. This underscores the need for the use of lower concentration of GO coating (<1 mg/ml) to balance antimicrobial efficacy while preserving biocompatibility for safer human applications. Many recent works have reported the integration of polymers such as Poly(2,3-dihydrothieno-1,4-dioxin)-poly(styrenesulfonate) (PEDOT:PSS) with GO, with advantages like improved electrical conductivity^58^, enhanced conductive pathways^59^, superior electron mobility^60^, improving GO adhesion^61^, and enhancing corrosion resistance^62^. These observations led us to explore GO-PEDOT:PSS hybrid coating on Al (AA1050) containing lower concentrations of GO in order to achieve a balance between antimicrobial potential and biocompatibility while optimizing corrosion resistance.

However, several key research questions arise: (1) How does the lower concentrations of GO influence the antifouling efficacy of GO-PEDOT:PSS hybrid coated Al (Al_GO/P) substrates against broad-spectrum bacterial and fungal pathogens?; (2) Which GO concentration range provides the optimal balance between antibiofilm activity and biocompatibility?; (3) Whether the superior antimicrobial efficacy of Al_GO/P can be correlated to known mechanisms such as higher surface hydrophobicity and ROS production relative to bare Al?; (4) Whether Al_GO/P surfaces reduce the cytotoxicity by reducing the leaching compared to GO coating alone on Al (Al_GO)?; and finally (5) Do the Al_GO/P substrates exhibit higher biocorrosion resistance relative to uncoated aluminum and Al_GO substrates under similar physiological conditions? Addressing these questions is essential for advancing healthcare-related applications through the strategic optimization of GO-PEDOT:PSS hybrid coatings on AA1050 substrates^57^.

In the present study, a concentration dependent biocompatibility assessment highlighted the safety of PEDOT:PSS at 1 µg/ml. Next, GO at lower concentrations (50, 100, 250, and 500 µg/ml) was integrated with PEDOT:PSS (1 µg/ml), which was electrochemically deposited on Al (AA1050) substrates (Al_GO/P). The resulting substrates were designated as Al_GO50P, Al_GO100P, Al_GO250P, and Al_GO500P. For a comparative assessment, suitable controls such as glass, bare Al, and GO coated Al substrates (Al_GO) without PEDOT:PSS at respective GO concentrations (Al_GO50, Al_GO100, Al_GO250, and Al_GO500) were fabricated under similar deposition conditions. The structural identity and elemental composition of both Al_GO and Al_GO/P substrates were verified using Raman spectroscopy and X-ray photoelectron spectroscopy (XPS). Surface wettability and electrical conductivity were examined through static water contact angle (WCA) measurements and four-point probe analysis, respectively. Furthermore, FESEM analysis was performed on the Al_GO/P substrates to examine the uniformity of surface deposition, while porosity measurements were carried out to evaluate porosity percentage of the Al_GO/P substrates.

The biological performance of Al_GO/P and control substrates were evaluated against clinically relevant pathogens; Gram-negative; *Escherichia coli* (*E. coli*), Gram-positive; *Staphylococcus aureus* (*S. aureus*), and fungus; *Candida albicans* (*C. albicans*), commonly associated with healthcare-associated infections (HAIs). Initial studies included growth inhibition assay and viability assay for both control and test Al_GO/P substrates for a qualitative and quantitative evaluation of antimicrobial efficacy. Antifouling activity of substrates was further assessed by examining mixed species biofilm formation in presence of control and test Al_GO/P substrates via fluorescence microscopy and crystal-violet staining. Further mechanistic understanding was obtained by evaluating the generation of intracellular and surface reactive oxygen species (ROS) following exposure to various substrates.

A detailed corrosion analysis was conducted by measuring corrosion rates under abiotic and biotic conditions, along with FESEM images showing pre- and post-immersion condition. Further assessment of MIC potential and surface leaching for the test substrates were conducted via pH monitoring and inductively coupled plasma-mass spectroscopy (ICP-MS) analysis. Biocompatibility of the synthesized substrates was monitored via direct-contact and extract (indirect) methods. Overall, this systematic experimental framework underscores the enhanced antifouling, biocorrosion resistance, and biocompatibility characteristics of the Al_GO/P hybrid substrates relative to control surfaces. However, Al_GO100P and Al_GO250P are found to be the most suitable candidates as *in vivo* implant materials, balancing antimicrobial efficacy, and optimal biocompatibility. Al_GO500P may find promising *in vitro* applications due to its excellent biocidal activity. These observations collectively establish selected Al_GO/P substrates as favorable candidates for next-generation biomedical implant materials, water purification systems, and industrial applications, to name a few, where biofilm mitigation and long-term stability are essential.

## 2. Experimental Procedures

### 2.1. Substrates synthesis and characterization

#### 2.1.1. Substrate preparation^57,62^

Aluminium plates (AA1050) with 99.5% purity and a thickness of 1.5 mm were obtained from NALCO India Ltd. The pristine graphene oxide (GO, 0.4 wt%) was purchased from Graphenea Inc., and poly(3,4-ethylenedioxythiophene): polystyrene sulfonate (PEDOT:PSS) was obtained from Sigma-Aldrich (Cat. no. #655201). For ease of handling, aluminum sheets were sectioned into 2.5 cm × 2.5 cm and 1.5 cm x 1.5 cm squares using an electric discharge machine (EDM) wire cutter. The aluminum substrates were then polished using silicon carbide emery papers of grit sizes ranging from P320 to P2000 to improve surface suitability for electrochemical deposition.

Subsequently, the substrates underwent ultrasonic cleaning (100W, 40 kHz) in acetone [Cat. no. #31566, SRL] for 30 minutes to eliminate any remaining polishing residues and were dried at room temperature.

A facile electrochemical deposition (ECD) process, involving optimization of an aqueous electrolyte, was used to coat the sterile aluminium substrate with a GO-PEDOT:PSS hybrid layer (Al_GO/P). The GO slurry was prepared by dispersing GO in 0.1 M sodium sulfate (Na_2_SO_4_) [Cat. no. #59977, SRL] to form a Na_2_SO_4_-GO suspension. The mixture was probe-sonicated for 1 hour to ensure uniform dispersion.

GO suspensions of 50 µg/ml, 100 µg/ml, 250 µg/ml, and 500 µg/ml were prepared. The pH of each suspension was adjusted to a neutral value of 7.0 by adding 0.1 M sodium hydroxide (NaOH) [Cat. no. #68151, SRL] dropwise under continuous stirring.

At this stage, prior to deposition, PEDOT:PSS was added to the GO suspension at a concentration of 1 µg/ml (25 µL).

Electrochemical deposition was carried out in a glass beaker using the cleaned aluminum sheet as the working electrode and a pure copper sheet as the counter electrode. A constant positive DC voltage of 30 V was applied using a Keithley 2601B source meter for 90 seconds to achieve a homogenous, uniform deposition. GO coated AA1050 (Al_GO) substrates were also prepared without PEDOT:PSS, at different GO concentrations, for a comparative analysis.

Before proceeding to biological experiments, all bare and coated substrates were sterilized by autoclaving at 121°C under 15 lb pressure for 15 minutes, followed by ultraviolet light irradiation for 30 minutes to ensure sterility. Finally, all samples were washed twice with phosphate-buffered saline (PBS) and stored under sterile conditions until use.

#### 2.1.2. Spectroscopic analyses: Raman and X-ray photoelectron spectroscopy ^57,63^

The structural identity of the GO layer in the Al_GO and Al_GO/P samples was characterized using a Renishaw inVia^TM^ confocal Raman microscope. The Raman spectra were recorded at 25°C using a 532 nm laser as the excitation source. Spectral data were collected in the range of 101 cm^-1^ to 3202 cm^-1^ with a resolution of 0.8 cm^-1^.

XPS measurements were performed using a ULVAC-PHI (PHI 5000 Versa Probe III) system with nonmonochromatic Al Kα X-rays (1486.70 eV). XPS measurements were carried out in an ultra-high vacuum (UHV) chamber with a base pressure of < (6.75 x 10−10 Torr). The high-resolution spectra were recorded at a pass energy of 55 eV and the survey spectra were recorded at a pass energy of 280 eV. XPS peak fitting was performed using Casa XPS software.

#### 2.1.3. Surface morphological analysis: Field emission scanning electron microscopy (FESEM)^57^

FESEM analysis was performed to assess the surface morphology and microstructural variations between the bare aluminum and Al_GO/P-coated substrates. Micrographs were obtained using a JSM-7200F Schottky FESEM (JEOL Ltd., TIFR Hyderabad Facility). The samples (2.5 cm × 2.5 cm) were placed in an ultrahigh vacuum chamber (10^-6^ mbar) and irradiated with high-energy electrons at an accelerating voltage of 5 kV.

To enhance surface conductivity and image resolution, samples were sputter-coated for 2 minutes with a thin layer of gold (1∼2 nm) prior to imaging. Observations were conducted using the secondary electron (SE) detection mode to obtain high-contrast micrographs of surface features.

#### 2.1.4. Water Contact Angle (WCA)^63,64^

The surface wettability of bare Al, Al_GO, and Al_GO/P substrates was evaluated by measuring the static water contact angle (WCA) using a VCA Optima contact angle meter (AST Products Inc.). The sessile drop technique was employed, in which a droplet of ultrapure water was placed on the surface of each specimen. Images of the droplet were captured and analyzed to determine the contact angle, thereby indicating the relative hydrophobicity or hydrophilicity of the surface. The measurements were repeated at three different points per sample to ensure reproducibility.

#### 2.1.5. Four-probe conductivity measurement ^65^

Bulk electrical conductivity of bare Al, Al_GO, and Al_GO/P substrates was measured using the Van der Pauw method. A four-probe station and Keithley 2450 source meter (delta mode) were used for this purpose. Electrical contacts were placed at the edges of the substrate, and a 10 mA current was applied to measure the sheet resistance (*R_s_*).

The bulk conductivity (*σ*) was calculated using the relation:

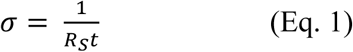

where *t* is the thickness of the aluminum substrate. For each sample, measurements were acquired at three different points, and the resulting data were expressed as mean ± standard deviation computed using GraphPad Prism version 9.0 for Windows. The experiment was conducted under 25 ± 1℃ with 40% relative humidity (non-condensing) conditions, and the conductivity values were compared across different GO concentrations.

#### 2.1.6. Porosity Evaluation ^57,66–68^

The porosity of Al_GO/P substrates were evaluated at standard laboratory temperature (25 ± 1°C) using electrochemical techniques in a 3.5 wt.% NaCl solution,^57,66–68^ simulating a corrosive environment. Due to the low intrinsic internal resistance of the electrolyte solution (<5 Ω), iR compensation was not necessary. A three-electrode setup was configured using a VSP-300 multichannel potentiostat/galvanostat (BioLogic, France). The working electrode was either a bare or coated aluminum substrate, the reference electrode was Ag/AgCl, and the counter electrode was a platinum mesh.

The active surface area was defined using an epoxy mask, exposing only 1 cm^2^ of the sample. The open circuit potential (OCP) was monitored for 30 minutes until stabilization. Tafel polarization was carried out at a scan rate of 1 mV/s, over a potential range of ±250 mV relative to OCP at room temperature. Tafel parameters, i.e., corrosion potential (*E_corr_*) and anodic Tafel slope (*β_a_*) for bare Al and Al_GO/P substrates were extracted using Tafel fit function of EC-Lab software. Polarization resistance (R_p_) for bare Al and Al_GO/P samples were determined from the same Tafel data using EC-Lab’s R_p_ fit function, as per the approach of Creus *et al.*^69^

Electrochemical impedance spectroscopy (EIS) was performed with an AC amplitude of 10 mV over a frequency range of 100 kHz to 100 mHz, but was not used for quantitative porosity measurements.

The porosity *P* (%) was calculated using the following equation^69^,

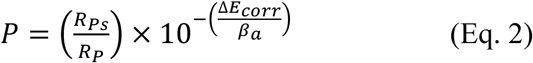

where *R_PS_* is the polarization resistance of bare aluminium, *R_p_* the polarization resistance of the coated substrates, *ΔE_corr_* the difference between the corrosion potential of the coated (*E_corr,coatings)_* and bare (*E_corr,Al_*) substrates (*ΔE_corr_=E_corr,coatings_ - E_corr,Al_*), and *β_a_* the anodic Tafel slope of the bare aluminum.

### 2.2. Antimicrobial and Antibiofilm Assay

#### 2.2.1. Microbial cell culture

*Candida albicans* (*C. albicans*) [fungus, ATCC 76485] cells were cultivated on Yeast-Peptone Dextrose (YPD) agar [Cat. no. #G038, HiMedia] plates at 30°C for 24 hours. A single colony was selected and inoculated into YPD broth [Cat. no. #M1363, HiMedia], which was then incubated overnight at 25°C with shaking at 200 rpm. From this overnight culture, a secondary culture was initiated at a 1:100 dilution and allowed to grow until it reached the mid-logarithmic phase. The final fungal cell suspension was prepared to achieve a concentration of 10^8^ CFU/ml.

*Escherichia coli* (*E. coli*) [Gram-negative bacterium, ATCC 29425] and *Staphylococcus aureus* (*S. aureus*) [Gram-positive bacterium, ATCC 25923] cells were grown on Luria Bertani (LB) agar [Cat. no. #G1151, HiMedia] plates at 37°C for 24 hours. A single colony was picked and inoculated into LB broth [Cat. no. #G1245, HiMedia], followed by overnight incubation at 37°C with shaking at 200 rpm. From this culture, a secondary inoculum was prepared using a 1:100 dilution, which was then grown to the mid-logarithmic phase. The final bacterial cell suspension was prepared to achieve concentrations of 10^7^ CFU/ml and 10^8^ CFU/ml, depending on the assay.

For mixed *Candida*-bacterial biofilms, a suspension was prepared by mixing 1 ml each of the *C. albicans*, *S. aureus*, and *E. coli* cultures in a 1:1:1 ratio ^70^.

#### 2.2.2. Microbial Growth Inhibition Assay and Cell Viability Assay^63,71^

For growth inhibition assay, Al_GO/P substrates (2.5 cm × 2.5 cm) were sterilized by autoclaving at 121°C for 15 minutes under 15 lb pressure, followed by ultraviolet light exposure for 30 minutes. 100 µL of bacterial suspension containing 10^7^ CFU/ml was pipetted onto different Al_GO/P and control substrates (Glass and Bare Al) inside sterile Petri plates. These were incubated at 37°C for 2 hours under aseptic conditions. After incubation, molten LB agar (45°C) was poured over the substrates and allowed to solidify at room temperature. The plates were then incubated overnight at 37°C, and bacterial colonies were analyzed the following day. Similarly, 100 µL of *C. albicans* suspension (10^7^ CFU/ml) was incubated on distinct Al_GO/P and control substrates in sterile Petri plates for 10 hours under aseptic conditions. Following incubation, molten YPD agar (45°C) was poured over the surfaces and allowed to solidify. The plates were incubated overnight at 30°C, and fungal colonies were analyzed after 24 hours.

For cell viability assay, bacterial and fungal cells at 10^7^ CFU were incubated on sterilized glass, bare Al, Al_GO, and Al_GO/P substrates (150 μl/cm^2^) in sterile Petri plates at 37°C under constant 95% relative humidity (maintained via autoclaved deionized water channels in closed containers to prevent desiccation). Cells were harvested at 0, 5, and 10 h, serially diluted up to 10^5^-fold in normal saline, spread-plated on LB agar (bacteria) or YPD agar (fungi), and incubated overnight at 37°C (bacteria) or 30°C (fungi). Colonies were counted the following day, and loss of cell viability percentage was calculated using the formula:

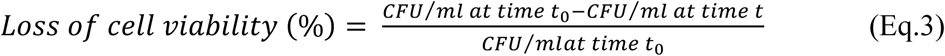

The results were plotted using GraphPad Prism version 9.0 for Windows, GraphPad Software, San Diego, CA, U.S.A. (www.graphpad.com).

#### 2.2.3. Antibiofilm assay^57^

A mixed microbial suspension was incubated on sterilized Al_GO/P substrates (150 μl/cm²) placed in sterile Petri dishes at room temperature. To maintain constant humidity (95% relative humidity), the plates were placed in closed containers containing autoclaved deionized water channels to prevent desiccation stress.

The mixed suspension was seeded onto both control and Al_GO/P-coated substrates and incubated for 10 hours. Two different inoculum concentrations (10^7^ CFU/ml and 10^8^ CFU/ml) were tested in parallel to assess biofilm formation. After incubation, 100 µL of each residual suspension was transferred to 35mm Petri plates containing a 1:1 mixture of YPD and LB broth, supplemented with 10% fetal bovine serum (FBS) [Cat. no. # A5256901, Gibco], and incubated at 37°C for an additional 10 hours.

#### 2.2.4. Fluorescence microscopy to visualize biofilm formation

After 10 hours of incubation under biofilm-stimulating conditions (37°C with 10% FBS), biofilm-coated substrates were gently washed with sterile distilled water to remove planktonic cells. The remaining biofilm biomass was sequentially stained using dark conditions with:

- Calcofluor white (4 µg/ml) [Cat. no. #18909, Sigma-Aldrich] for *C. albicans* (blue) ^72–74^
- Acridine orange (2% solution in H_2_O) [Cat. no. #A9231, Sigma-Aldrich] for *S. aureus* (red)^75^
- Sodium fluorescein (0.0001% solution in ethyl alcohol) [Cat. no. #46960, Sigma-Aldrich] for *E. coli* (green)^76^

The dyes were prepared according to established protocols^76^. Fluorescence imaging was performed using an EVOS Cell Imaging Station (Invitrogen) at 40× magnification. The fluorescence images were quantified to calculate mean fluorescence intensity using ImageJ software^77^.

#### 2.2.5. Biofilm biomass quantification^78^

Following 10-hour incubation, 100 µL of each suspension was seeded into wells of a 96-well plate containing YPD medium supplemented with 10% FBS and incubated at 37°C under static conditions for an additional 10 hours.

The biofilms were washed three times with Milli-Q water to remove non-adherent cells. Then, 200 μl of 0.1% (w/v) crystal violet solution [Cat. no. #TC510, HiMedia] was added to each well, and the plates were incubated at room temperature for 30 minutes on an orbital shaker at 70 rpm.

Excess stain was removed with three washes using Milli-Q water, and the wells were dried at room temperature for 30 minutes. The retained stain was solubilized with 200 μl of 70% (v/v) ethanol, and absorbance was measured at 595 nm.

- Growth control (GC): Glass substrate (100% biofilm)
- Sterility control (SC): Ultrapure water (0% biofilm)

Percentage biofilm growth was calculated using the following equation:

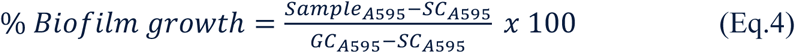

where *Sample_A595_* is the absorbance of the test substrate, *SC_A595_* the absorbance of the sterility control (0% biofilm), and *GC_A595_* the absorbance of the growth control (100% biofilm). The results were plotted using GraphPad Prism version 9.0 for Windows, GraphPad Software, San Diego, CA, U.S.A. (www.graphpad.com).

#### 2.2.6. Detection of ROS production^57,79^

Mixed *Candida-*bacterial suspension was seeded on sterile Al_GO/P substrates and incubated at room temperature for 10 hours under constant humidity (95% relative humidity). 10 µM of 2’,7’-dichlorodihydroflourescein diacetate (H_2_DCF-DA) [Cat. no. #D399, Invitrogen] was added to the suspension and incubated for 20 minutes in the dark. Imaging was performed using an in-house fluorescence microscope (Floid cell station, 40×, Invitrogen).

To detect surface-generated ROS, 10 µM of H_2_DCF-DA was applied directly onto washed substrates, and green fluorescence was recorded using a Nikon ECLIPSE Ci-E microscope at 40× magnification.

Mixed bacterial cell suspension was seeded on sterile Al_GO/P substrates and incubated at room temperature for 10 hours under constant humidity (95% relative humidity). A 10 µM solution of the cytosolic superoxide indicator dye dihydroethidium (DHE) [Cat. no. #D1168, Invitrogen] was used to stain the suspension for 20 minutes in the dark^80^. Following incubation, cell imaging was performed using an in-house fluorescence microscope (Floid Cell Station, 40×, Invitrogen).

To detect ROS generated at the material surface, a separate experiment was performed: mixed bacterial cell suspension was seeded on sterile Al_GO/P substrates and incubated at room temperature for 10 hours under constant humidity (95% relative humidity). After incubation, the substrates were rinsed and placed under a fluorescence microscope. A 10 µM solution of DHE was then applied directly to the surface of both the control and Al_GO/P substrates. The resulting red fluorescence signal was captured using a Nikon ECLIPSE Ci-E microscope at 40× magnification. All the fluorescence images were quantified to calculate average fluorescence intensity per ROS positive cell using ImageJ software^77^. The results were plotted using GraphPad Prism version 9.0 for Windows, GraphPad Software, San Diego, CA, U.S.A. (www.graphpad.com).

#### 2.2.7. Scanning electron microscopy (SEM)^57,63^

10^7^ CFU of the mixed *Candida-*bacterial suspension were incubated over sterilized Al_GO/P substrates (150 μl/cm^2^) placed in sterile Petri plates at room temperature for 10 hours under constant humidity (95% relative humidity). Following incubation, the cells were fixed with 2.5% glutaraldehyde and dehydrated through a graded ethanol series (30%, 60%, 70%, 80%, 90%), followed by treatment with 100% ethanol for 30 minutes.

The substrates were then sputter-coated with gold-palladium (1∼2 nm) for 2 minutes and imaged using a Gemini SEM 450 (Zeiss, KIIT Central Research Facility).

### 2.3. Evaluation of Corrosion Resistance

#### 2.3.1. Description of immersion systems ^42^

The experiments were carried out in two systems consisting of glass beakers with a usable volume of 300 ml, in which the substrates were fixed to the bottom after prior sterilization. The medium employed was a 1:1 mixture of LB and YPD broth, sterilized in an autoclave at 121°C and 15 lb pressure for 15 minutes. Both systems were assembled under aseptic conditions.

The abiotic system (control) contained only the sterile medium along with the following antibiotics: Fluconazole (antifungal, 15 µg/ml) [Cat. no. #22239, SRL], Chloramphenicol (antibacterial, 15 µg/ml) [Cat. no. #97686, SRL], and Vancomycin (antibacterial, 15 µg/ml) [Cat. no. #61078, SRL]. The biotic system included the same sterile medium, supplemented with 1% (v/v) of the mixed *Candida*-bacterial inoculum. Both the immersion systems were maintained at 25 ± 1℃ over 45 days.

#### 2.3.2. Weight loss measurement of the corroded substrates

After 45 days of exposure in both biotic and abiotic systems, weight loss was calculated following the chemical cleaning procedure described in ASTM G1-03 standard^81^. After removal of the biofilm, the samples were immersed in a 69.2 wt% nitric acid solution for approximately 5 minutes to remove corrosion products, rinsed with distilled water to eliminate residual acid, and then dried under flowing air.

The specimens were weighed before and after the experiment using an analytical balance, and the corrosion rate was calculated using the following equation:

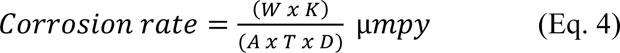

where *W* is the weight loss, *K* the unit conversion constant (8.76 x10^7^), *A* the exposed area, *T* the exposure time, *D* the material density, and *µmpy* the corrosion rate in micrometer per year.

The measured corrosion rates were classified according to the NACE Standard RP-07-75^82^, which provides validated guidelines for the preparation, analysis, and interpretation of corrosion measurements. All tests were performed in triplicates. Additionally, the pH of abiotic and biotic immersion systems were monitored over 45 days. pH values were measured daily, and the average value was determined from two measurements from the same day to minimize experimental error. The results were plotted using GraphPad Prism version 9.0 for Windows, GraphPad Software, San Diego, CA, U.S.A. (www.graphpad.com).

#### 2.3.3. Inductively Coupled Plasma-Mass Spectroscopy (ICP-MS) analysis of immersion medium^83^

After 45 days of incubation of the control and test substrates under abiotic and biotic immersion conditions, the spent media was collected for ICP-MS sample preparation. Samples were acid-digested with concentrated nitric acid (HNO_3_, 65%) and incubated overnight at 25 ± 1°C to ensure complete digestion of dissolved and particulate species. The digested solutions were then diluted to a final volume of 50 ml with ultrapure deionized water and filtered through 0.22 µm syringe filters to remove any remaining particulates. The filtrates were subsequently analyzed using an ICP–MS (NexION 5000, PerkinElmer, USA).

#### 2.3.4. FESEM analysis of immersed Al_GO/P substrates

FESEM was used to investigate microstructural variations on bare aluminum, CR steel (positive control), Al_GO and Al_GO/P substrates before and after 45-day immersion in both abiotic and biotic environments.

To observe the corrosion morphology, the substrates were cleaned in accordance with the ASTM G1-03 standard^81^. Micrographs were obtained using a JSM-7200F Schottky FESEM (JEOL Ltd., TIFR Hyderabad Facility), operated at an accelerating voltage of 30 kV. Samples of 2.5 cm × 2.5 cm were placed in an ultrahigh vacuum chamber (10^-6^ mbar). Prior to imaging, substrates were coated with a thin gold film (1∼2 nm) for 2 minutes to enhance surface conductivity and enable high-resolution image capture.

### 2.4. Cytotoxicity assay

#### 2.4.1. Mammalian Cell Culture ^84–87^

The Human Embryonic Kidney (HEK293T) cell line [ATCC CRL-11268] was used. HEK293T is a well-established model and is widely employed in cytotoxicity assays, including the evaluation of new biomaterials^88^. A frozen vial was retrieved from liquid nitrogen storage and thawed using a standard defrosting protocol. The cells were cultured in Dulbecco’s Modified Eagle Medium (DMEM) [Cat. no. #10564011, Gibco] supplemented with 10% fetal bovine serum (FBS) [Cat. no. # A5256901, Gibco] and 1X Antibiotic-Antimycotic solution [Cat. no. #A002, HiMedia]) for three passages prior to experimentation.

Cells were maintained at 37°C in a humidified incubator with 5% CO_2_ and 95% air. Upon reaching approximately 80% confluency, biocompatibility experiments were initiated. Cell counting was performed using a hemocytometer. Samples were prepared according to Sections 4.2 and 4.3 of ISO 10993-5: Biological Evaluation of Medical Devices - Part 5: Tests for In Vitro Cytotoxicity, and the experiments were performed in compliance with Section 8 of the same standard^89^.

#### 2.4.2. Indirect (extract) assay

HEK293T cells (passage 3) were seeded into 96-well plates at a density of 1×10^4^ cells per well and incubated for 24 h. In parallel, extracts of the test materials were prepared. After sterilization with 70% ethanol and ultraviolet light exposure for 30 minutes, the Al_GO/P substrates were placed in 35 mm Petri plates containing fresh growth medium. The extraction procedure was carried out at 37℃ for 24 hours to obtain 100% crude extracts.

Once the seeded cells reached approximately 70% confluency, the culture medium was replaced with the test extracts. Cells were incubated for an additional 24 and 48 hours. Each condition was tested in triplicate.

Cell viability was assessed by incubating the cells in medium containing 0.5 mg/ml of 3-[4,5-dimethylthiazol-2-yl]-2,5-diphenyltetrazolium bromide (MTT) [Cat. no. #CCK003, HiMedia] for 2 hours. The medium was then replaced with isopropanol to solubilize the purple formazan product, and absorbance was measured at 570 nm. Results are expressed as the percentage of MTT signal relative to untreated control cells, which were defined as 100% viable. Experiments were conducted according to Section 8.2 of ISO 10993-5: Biological Evaluation of Medical Devices - Part 5: Tests for In Vitro Cytotoxicity^89^. The results were plotted using GraphPad Prism version 9.0 for Windows, GraphPad Software, San Diego, CA, U.S.A. (www.graphpad.com).

#### 2.4.3. Direct contact assay

HEK293T cells (passage 3) were seeded at 1 × 10^4^ cells per 35 mm Petri dish containing pre-sterilized substrates (1.5 cm × 1.5 cm; 2.25 cm^2^ per 10^4^ cells) for direct contact assays. HEK293T cells were incubated for 24 and 48 hours in the presence of test materials, which were carefully placed in the center of each plate. Each test material was evaluated in triplicates.

After incubation, cell viability was measured directly by adding 0.5 mg/ml MTT reagent to the culture medium and incubating for 2 hours. The medium was then replaced with isopropanol to extract the formazan dye, and absorbance was measured at 570 nm. Results were reported as percentage MTT signal compared to the control, where untreated cells were set at 100% viability. All procedures were carried out in compliance with Section 8.3 of ISO 10993-5: Biological Evaluation of Medical Devices - Part 5: Tests for In Vitro Cytotoxicity^89^. The results were plotted using GraphPad Prism version 9.0 for Windows, GraphPad Software, San Diego, CA, U.S.A. (www.graphpad.com).

### 2.5. Statistical Analyses

All statistical analyses were performed by one-way analysis of variance (ANOVA) using Graph Pad Prism version 9.0 for Windows, GraphPad Software, San Diego, CA, U.S.A.(www.graphpad.com). The data are presented as mean ± standard deviation. The statistical significance (p-value) between the control and experimental groups is denoted by * (p < 0.05), or ** (p < 0.01), or *** (p < 0.001), or **** (p<0.0001).

## 3. Results and Discussion

### 3.1. Material property assessment

#### 3.1.1. Synthesis of Graphene oxide – PEDOT:PSS hybrid coated Aluminium (AA1050) substrates

The schematic diagram illustrating the facile two-step synthesis process for graphene oxide-coated Al substrates is presented in Figure 1. The sample preparation parameters for coating AA1050 substrates using GO (50, 100, 250, and 500 µg/ml) in presence and absence of PEDOT:PSS are detailed in Table 1. The inherent negative surface charge (zeta potential) of the aqueous GO dispersion causes anodic deposition at applied potentials of 10 volts or higher^57^. These electrostatic forces were sufficient to drive GO nanosheets toward the working electrode *–* the Al substrate^90^. PEDOT:PSS was used at a concentration of 1 µg/ml in the hybrid system to preserve biocompatibility and conductivity while enhancing the adhesion of GO to the underneath metal (Figure S23, Supporting Information). GO coated AA1050 (Al_GO) substrates were prepared without PEDOT:PSS, at different GO concentrations, for a comparative analysis.

**Figure 1.**
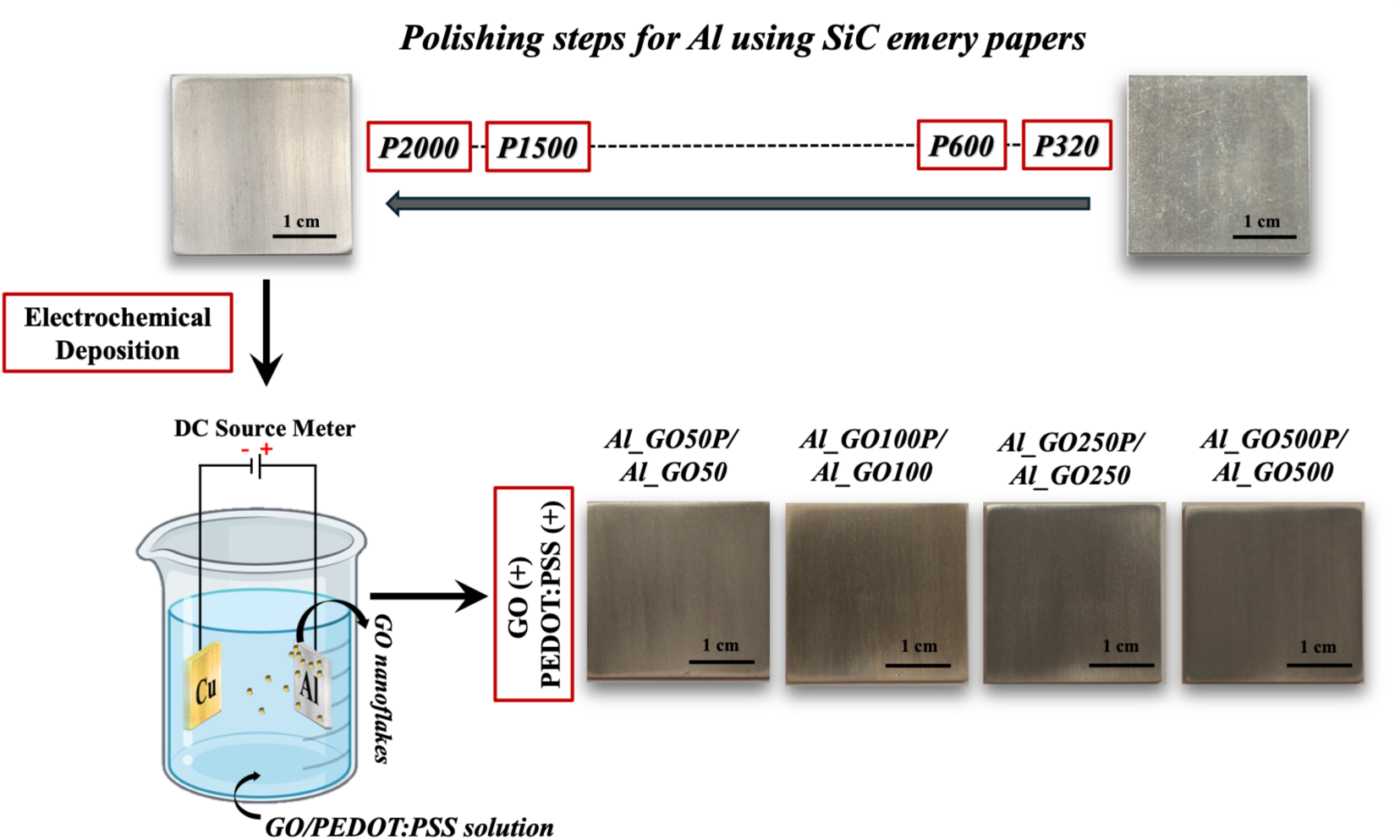
Facile two-step synthesis process of GO coated AA1050 substrates. Schematic diagram illustrating the preparation of Al_GO and Al_GO/P substrates, including sequential polishing of aluminum using SiC papers, followed by electrochemical deposition.

**Table 1.**
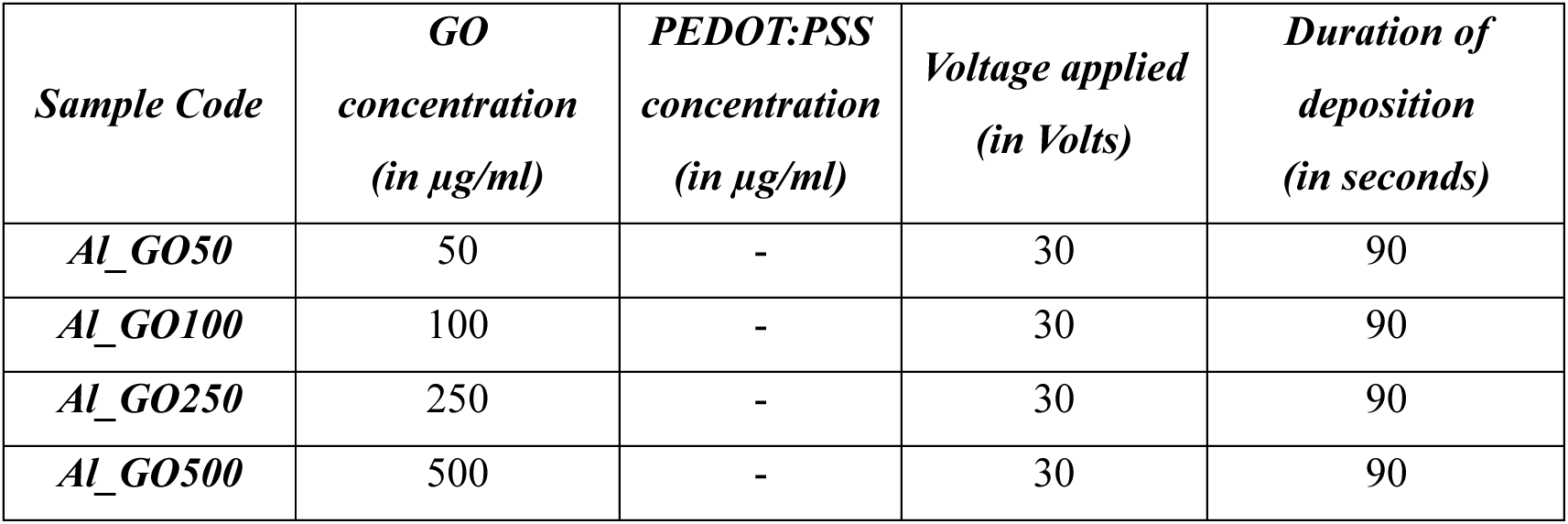
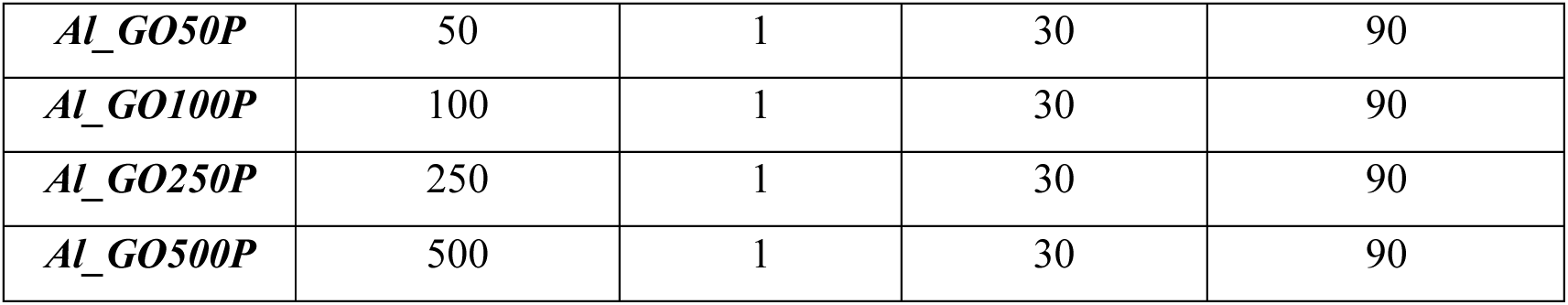
The details of sample preparation parameters used for coating Al substrates using varied concentration of graphene oxide sourced from Graphenea Inc.

| *Sample Code* | *GO concentration (in $\mu\text{g/ml}$)* | *PEDOT:PSS concentration (in $\mu\text{g/ml}$)* | *Voltage applied (in Volts)* | *Duration of deposition (in seconds)* |
| --- | --- | --- | --- | --- |
| *Al_GO50* | 50 | - | 30 | 90 |
| *Al_GO100* | 100 | - | 30 | 90 |
| *Al_GO250* | 250 | - | 30 | 90 |
| *Al_GO500* | 500 | - | 30 | 90 |
| *Al_GO50P* | 50 | 1 | 30 | 90 |
| *Al_GO100P* | 100 | 1 | 30 | 90 |
| *Al_GO250P* | 250 | 1 | 30 | 90 |
| *Al_GO500P* | 500 | 1 | 30 | 90 |

#### 3.1.2. Raman Spectrophotometry confirmed identity of GO on Al_GO/P and Al_GO substrates

Raman spectroscopy serves as a widely employed tool for acquiring structural insights into carbon-based materials. The identity of commercial PEDOT:PSS was confirmed via Raman spectroscopy (Figure S1, Supporting Information). The most prominent Raman band, located around 1437 cm^-1^, consistently exhibits a well-reproducible symmetric profile with a mixed pseudo-Voigt line shape (Figure S1, Supporting Information). Interestingly, this prominent band also displays the most substantial variation, encompassing a total shift of approximately 10 cm^-1^ within the PEDOT:PSS complex. Furthermore, our observations of the Raman signatures for the PEDOT:PSS utilized in this study are in strong concurrence with the Raman spectrum obtained from previous research^91,92^. This agreement underscores the reliability and consistency of the Raman spectroscopic characteristics of the PEDOT:PSS material used in the coating process.

The Raman spectra of graphitic carbon-based materials primarily manifest the G and D peaks, and their overtones. In Figure 2A, we present the Raman spectra of GO, Al_GO and Al_GO/P substrates. Notably for Al_GO and Al_GO/P substrates, we identified the prominent graphitic Raman peaks, namely G band at 1594 cm^-1^, and the D band at 1328 cm^-1^, correlating with prior literature^57,64^. The G peak primarily arises from the bond stretching of sp^2^ carbon atom pairs, encompassing both ring and chain structures^93^. In contrast, the D peak is indicative of structural defects within the graphene lattice. Moreover, the variation in intensity ratios of the D and G bands reveal the characteristics of GO deposited on Al_GO/P substrates as shown in Table S1 (Supporting Information). GO exhibited an I_D_/I_G_ ratio of 0.91, which shifted slightly to 0.93 and 0.91 for AlGO50 and AlGO50P, respectively. This ratio increased to 0.97 for AlGO100 but declined to 0.88 for AlGO100P. The I_D_/I_G_ ratio increased to 1.07 and 0.94 for AlGO250 and AlGO500, whereas both AlGO250P and AlGO500P exhibited 0.86 (Table S1, Supporting Information). These results confirm the successful deposition of GO on various Al_GO and Al_GO/P surfaces as shown in prior literature^57,94^.

**Figure 2.**
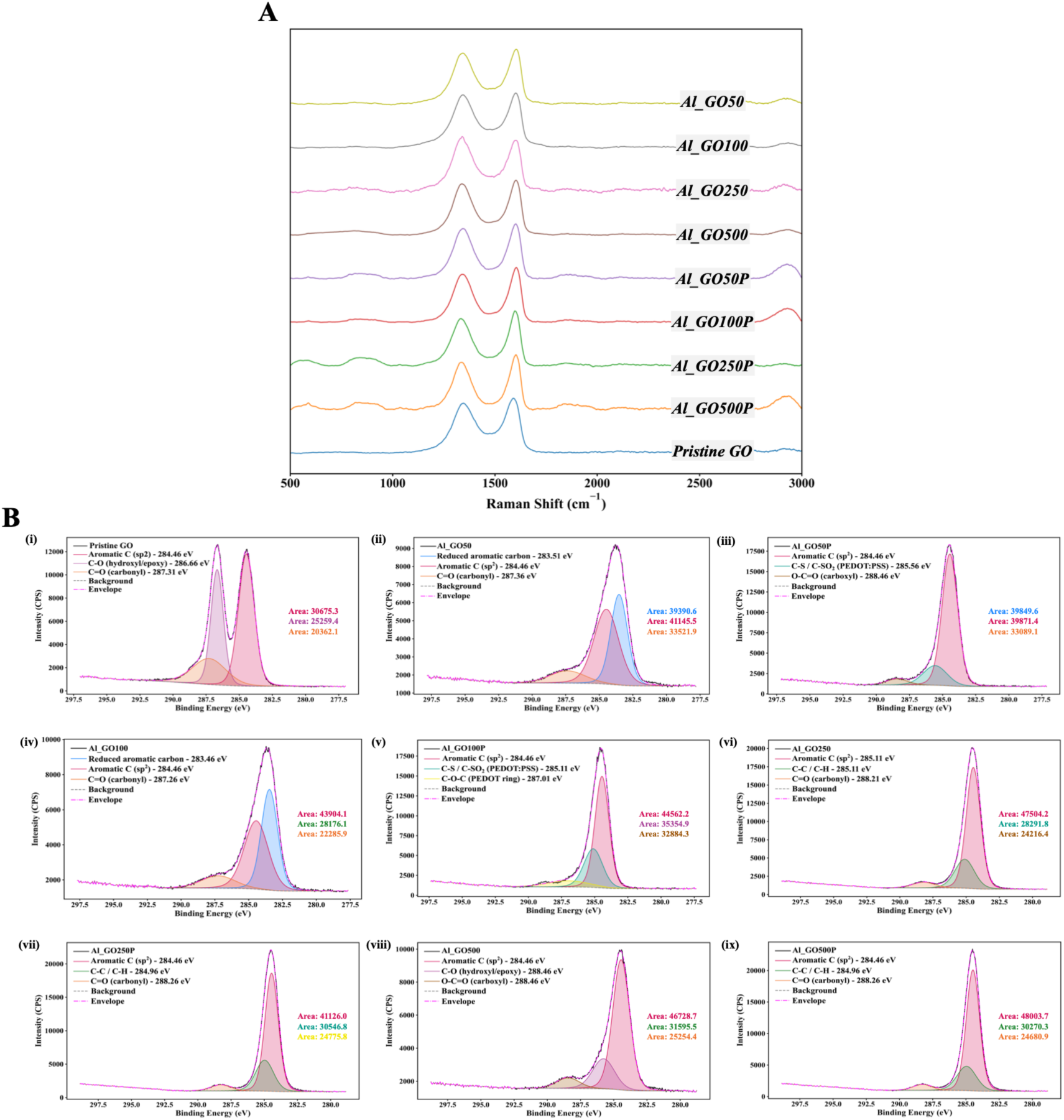
(A) Raman spectra of pristine GO, Al_GO, and Al_GO/P substrates. The figure shows the Raman signatures (D and G peaks) of GO and the fabricated Al_GO and Al_GO/P substrates. **(B) XPS analysis of pristine GO, Al_GO, and Al_GO/P substrates.** Deconvoluted peaks for the C_1s_ spectra attributing to different functional groups in (i) Pristine GO, (ii) Al_GO50, (iii) Al_GO50P, (iv) Al_GO100, (v) Al_GO100P, (vi) Al_GO250, (vii) Al_GO250P, (viii) Al_GO500, and (ix) Al_GO500P.

We have further calculated the full-width at half-maximum (FWHM) values presented in Table S1, Supporting Information. This clearly demonstrates that PEDOT:PSS modifies the structural environment of GO in a meaningful and systematic way. Smaller FWHM values for both D and G bands and consistently lower I_D_/I_G_ ratios in Al_GO/P substrates (Table S1, Supporting Information) point towards more ordered and less defective carbon framework^95^. This improvement can be attributed to the interactions between GO and PEDOT:PSS. Oxygen atoms from the sulfonate groups of PSS may participate in strong electrostatic and hydrogen bond interactions with GO’s hydroxyl, epoxy, carbonyl and carboxy functionalities^96^. Likewise, PEDOT provides positively charged sites that interact with the negatively charged functionalities of GO^97^. Such interactions may stabilize the GO sheets, reducing lattice distortions and defects^98^. Thus PEDOT:PSS not only coats the GO surface but also reorganizes the carbon lattice making it more structurally ordered and defect free.

Next, XPS was performed to understand the chemical modifications induced by PEDOT:PSS incorporation into GO across the test concentrations (50, 100, 250 and 500 µg/ml). Survey spectra confirmed the presence of carbon and oxygen as the dominant elements in the Al_GO and Al_GO/P substrates (Figure S2A, Supporting Information). High-resolution C_1s_ spectra were deconvoluted to resolve the contributions from aromatic sp^2^ carbon, C-O groups, carbonyl groups and C-S/C-S-O functionalities associated with PEDOT:PSS (Figure 2B).

A systematic distinction is observed between the Al_GO control and the Al_GO/P samples, particularly at lower GO loadings (50 and 100 µg/ml). In the Al_GO substrates (Al_GO50 and Al_GO100), the C_1s_ is dominated by aromatic sp^2^ carbon (∼ 284.4 eV), reduced aromatic carbon (∼ 283.4 – 283.5 eV) and a carbonyl component (∼ 287.3 to 287.5 eV)^99^. In contrast, the corresponding Al_GO/P samples (Al_GO50P and Al_GO100P) exhibited entirely different chemical signatures. First, the reduced graphitic carbon peak is absent, indicating PEDOT:PSS disrupts or masks the reduced sp^2^ domains present in the Al_GO samples^100^. Second, the Al_GO/P samples displayed new oxygenated and sulfur-containing groups (C-S/C-S-O peaks at ∼ 285.1 – 285.7 eV)^101^ and a distinct C-O (PEDOT ring) contribution (287.0 eV)^102^. These features are consistent with the chemical structure of PEDOT:PSS and confirm its successful incorporation into the GO matrix.

At higher GO concentrations (250 and 500 µg/ml), both Al_GO and Al_GO/P samples exhibited similar sets of functional groups (aromatic sp^2^ carbon, C-C/C-H and carbonyl species), reflecting the dominance of GO associated features at higher mass fractions^99^. Pristine GO (Figure 2Bi) exhibited the functionalities similar to Al_GO500 substrate. Nevertheless, even in these higher GO concentrations, the PEDOT:PSS containing Al substrates consistently shows higher C:O ratios (Table S2 and Figure S2B, Supporting Information), apart from the lower GO concentrated samples. All these collectively confirm the successful modification of the surface chemistry of GO by PEDOT:PSS.

#### 3.1.3. FESEM confirmed uniform deposition of GO on Al surfaces

Electrochemical deposition (ECD) of GO on Al surface is expected to produce a smooth and uniform coating, as the quality of the coating is determined by its homogeneity^62,103^. We employed an FESEM analysis of the bare Al surface and coated test surfaces (Al_GO/P) FESEM micrographs of the bare Al and Al_GO/P substrates are shown in Figure 3, providing visual evidence of GO nano-sheets overlaying on the Al substrate The polished Al surface exhibits a non-uniform and rough surface morphology characterized by the presence of potholes and cavities. Accordingly, the FESEM images of Al_GO/P substrates demonstrated smooth GO sheets with small wrinkles, folded at the edges as evidenced in existing literature^57,63^. FESEM micrographs, therefore, confirmed the uniform deposition of the GO coatings on Al_GO/P surface via ECD method.

**Figure 3.**
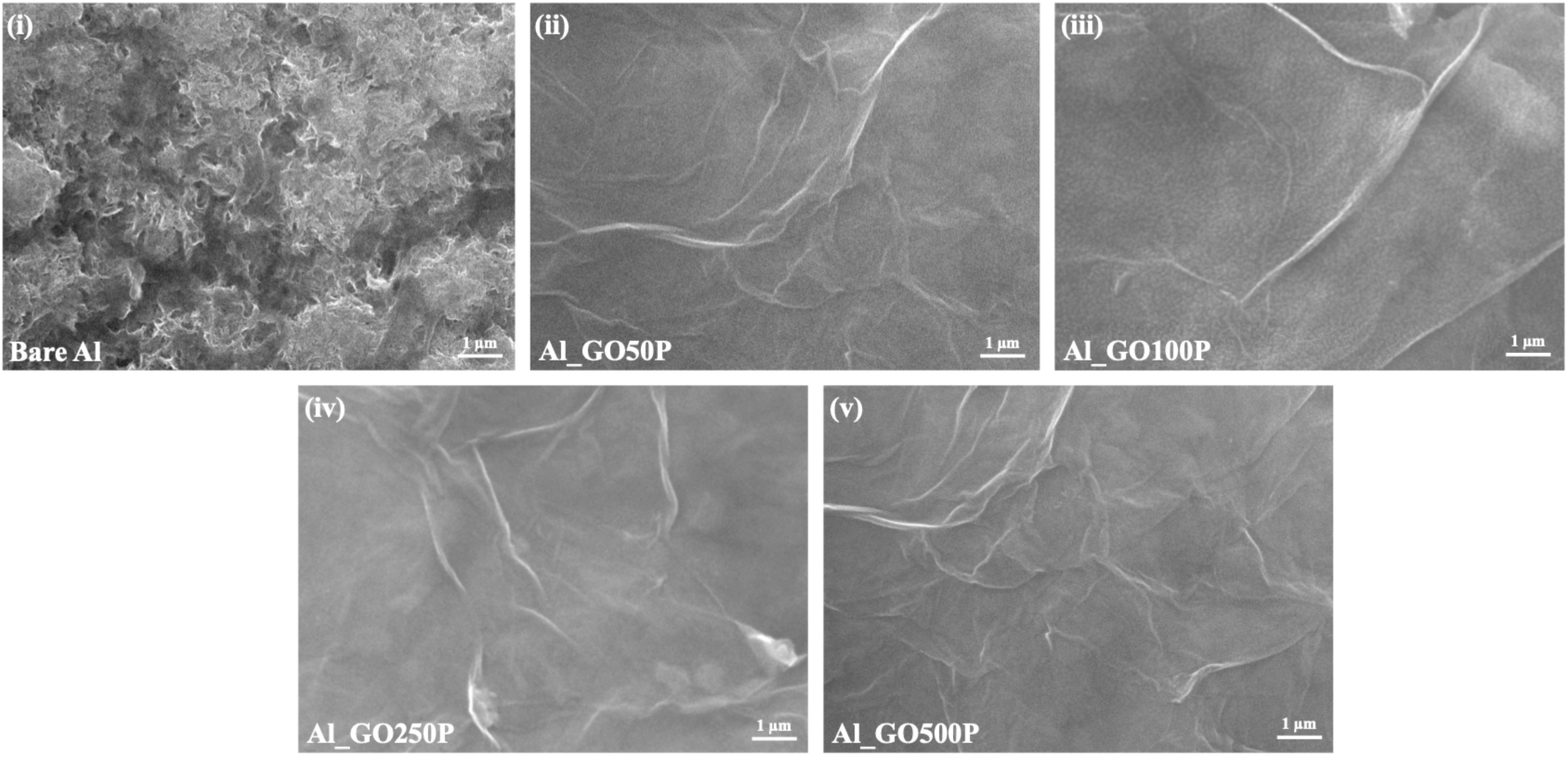
FE-SEM micrographs of bare Al vs Al_GO/P surfaces. FESEM images display (i) polished Al, and uniform deposition of GO on Al_GO/P substrates: (ii) Al_GO50P, (iii) Al_GO100P, (iv) Al_GO250P, and (v) Al_GO500P.

#### 3.1.4. Hydrophobic and Conductive nature of Al_GO/P surfaces confirmed by water contact angle (WCA) and four-probe conductivity measurements

The deposition of GO on Al substrates has been reported to enhance the surface hydrophobicity^57^. This increase in hydrophobicity is attributed to the uniform GO coating, which alters the surface energy and reduces wettability^104,105^. Current study evaluated the wettability of the bare Al, Al_GO, and Al_GO/P substrates, relying on the stationary WCA measurements (Table 2 and Figure S3, Supporting Information). The water contact angle of bare Al was 85.1°, but increased to over 90° for Al_GO and Al_GO/P surfaces, showing an increase in surface hydrophobicity (Table 2 and Figure S3, Supporting Information). Among the test samples, Al_GO500P had the highest WCA at 120.2°, followed by Al_GO250P (114.9°), Al_GO100P (106.8°), and Al_GO50P (101.6°). Following the similar trend, the control substrates, such as Al_GO500 had the highest WCA at 109.5°, followed by Al_GO250 (105.5°), Al_GO100 (103°), and Al_GO50 (94.2°). Taken together, these results indicate that increasing GO and GO-PEDOT:PSS loading changes the Al surface towards a hydrophobic state, which is expected to enhance the barrier properties against aqueous electrolyte penetration while simultaneously promoting closer microbial contact with the substrate surface^106^. In this work, a stable contact is advantageous, as it promotes a prolonged interaction of microbial cells to the GO-rich surface, thereby facilitating contact-mediated antimicrobial activity^57^.

**Table 2.** Stationary water contact angle (WCA) and electrical conductivity values of the synthesized Al_GO/P substrates and control surfaces; bare Al, Al_GO50, Al_GO100, Al_GO250, and Al_GO500.

| *Samples* | *WCA (in degrees)* | *Electrical Conductivity (S/m) x 10<sup>7</sup>* |
| --- | --- | --- |
| *Bare Al* | 85.1 ± 0.4 | 3.1 ± 0.7 |
| *Al_GO50* | 94.2 ± 4.3 | 1.8 ± 0.5 |
| *Al_GO100* | 103.0 ± 0.7 | 1.4 ± 0.2 |
| *Al_GO250* | 105.5 ± 1.2 | 1.4 ± 0.3 |
| *Al_GO500* | 109.5 ± 0.9 | 1.2 ± 0.2 |
| *Al_GO50P* | 101.6 ± 1.0 | 2.2 ± 0.5 |
| *Al_GO100P* | 106.8 ± 0.2 | 1.9 ± 0.2 |
| *Al_GO250P* | 114.9 ± 4.5 | 1.8 ± 0.2 |
| *Al_GO500P* | 120.2 ± 2.9 | 1.6 ± 0.4 |

However, the role of surface hydrophobicity in microbial attachment is more nuanced and strongly depends on the interplay between surface chemistry and physicochemical properties of the microorganisms themselves. Microorganisms generally exhibit stronger attachment on hydrophobic surfaces due to favorable hydrophobic–hydrophobic interactions and reduced interfacial free energy at the solid–liquid boundary^107,108^. On the other hand, hydrophilic surfaces tend to act as a physical and energetic barrier to microbial attachment, thereby suppressing early-stage colonization. In this context, the higher WCA values measured for the Al_GO/P substrates suggest that these surfaces may promote stronger microbial adhesion relative to bare Al, which is moderately hydrophilic.

Additionally, the electrical conductivity measurements for bare Al and Al_GO/P substrates are detailed in Table 2. It is worth noting that pure aluminum possesses inherently high electrical conductivity, registering at 3.43×10^7^ S/m (equivalent to 59.14% of the International Annealed Copper Standard, IACS), consistent with previous literature^109^. In the current work, conductivity of the aluminum alloy AA1050 was determined to be 3.11×10^7^ S/m, which is slightly lower than that of pure aluminum but falls within the same order of magnitude. It was observed that while the conductivity of the Al_GO and Al_GO/P substrates shows a relative reduction compared to the bare Al surface used in this study, the values were within the conductivity ranges for known materials ^110–112^. Moreover, this reduction in conductivity becomes more obvious with increasing GO concentration as detailed in Table 2. As the GO concentration increases, the conductivity of the Al_GO and Al_GO/P substrates decreases since GO disrupts the continuous sp^2^ carbon network needed for efficient electron transport, as in pristine graphene^113^. Higher GO loading introduces more oxygen-containing functional groups, which convert conductive sp^2^ regions into insulating sp^3^ regions and break up the π-electron pathways. This phenomenon may be attributed to the inherent insulating nature of graphene oxide, as previously documented^114^. It is plausible that the reduction in conductivity is a consequence of the expansion of interlayer spacing among the GO nanosheets due to the presence of functional groups within the basal planes^115^.

#### 3.1.5. Porosity evaluation of Al_GO/P substrates

The presence of porosity in coatings directly correlates to lower density and reduced compactness of the coating than the bulk metal^68^. Moreover, a porous coating decreases their protective ability as it allows foreign particles, microorganisms, or corrosive ions to reach and potentially damage the underlying metal, and cause corrosion^57,116^. The corrosion potential (E_corr_) and anodic potential (β_a_), and polarization resistance (R_p_) for bare aluminium and Al_GO/P substrates were determined through Tafel fit and R_p_ fit function of EC-Lab software utilizing the Tafel plot (Figure 4). The R_p_ fit and Tafel fit parameters are presented in Table S3 (Supporting Information), and the Nyquist impedance plots are displayed in Figure S4, Supporting Information.

**Figure 4.**
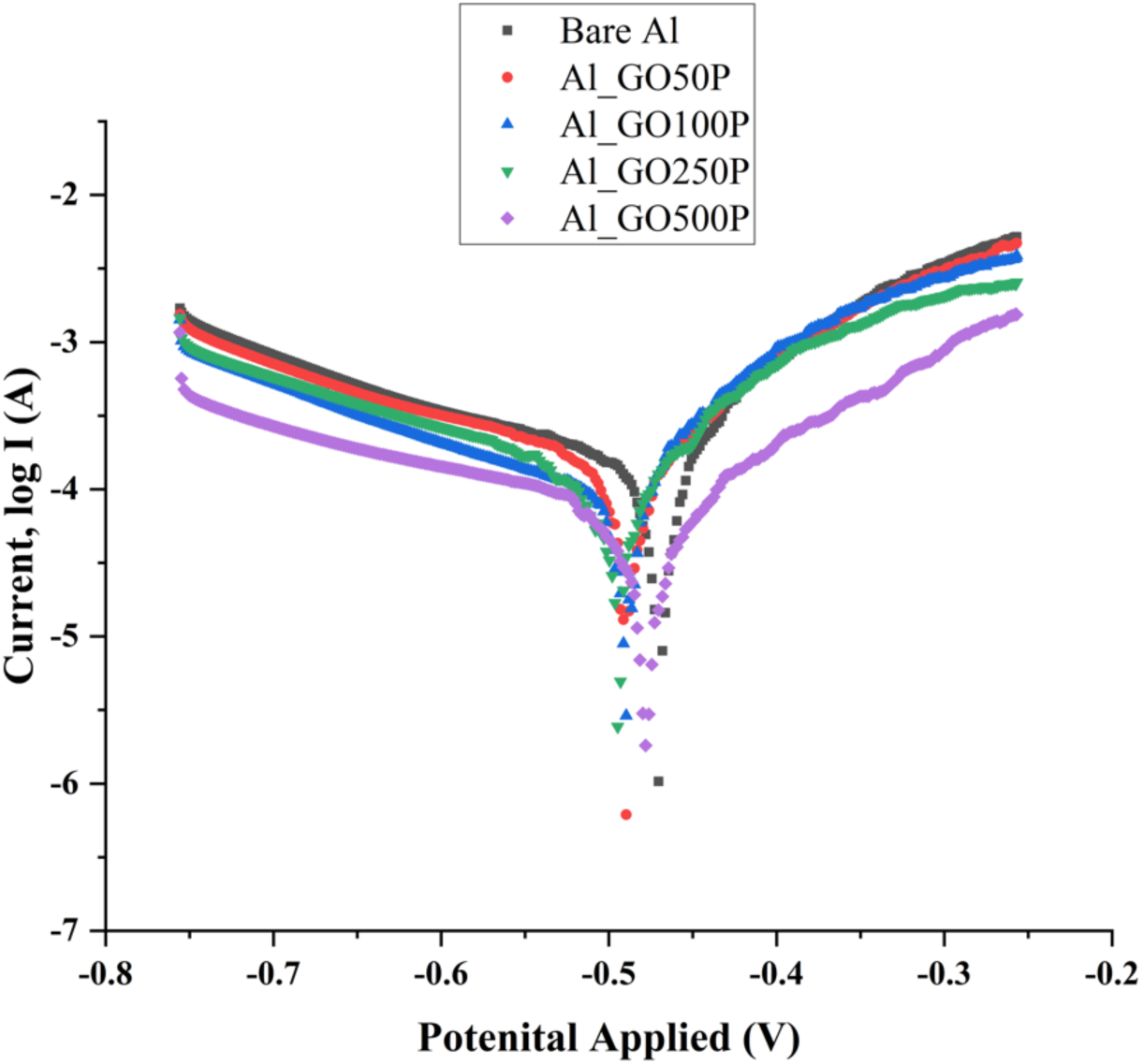
Tafel plot analysis. Linear polarization response comparison between bare Al vs Al_GO/P substrates.

The details for the porosity measurements for GO coatings is described in the Experimental Procedures and the results are provided in Table 3. The Al_GO/P substrates exhibited porosities of 89.1% (Al_GO50P), 82.5% (Al_GO100P), 65.4% (Al_GO250P), and 32% (Al_GO500P), demonstrating a gradual reduction in porosity with increased GO concentration(s) on the Al surface(s).

**Table 3.** Electrochemical characteristics of bare Al and Al_GO/P substrates and porosity (%) calculation.

| Sample | Polarization Resistance, $R_{PS}$ | Polarization Resistance, $R_p$ | Corrosion Potential, $E_{corr,Al}$ (V) | Corrosion Potential, $E_{corr,Coatings}$ (V) | $\Delta E_{corr}$ | Anodic Potential, $\beta_{a,Al}$ (mV) | Porosity (%) |
| --- | --- | --- | --- | --- | --- | --- | --- |
| Al_GO50P | 145 | 163 | -0.47 | -0.49 | -0.02 | 32.40 | 89.10 |
| Al_GO100P | 145 | 176 | -0.47 | -0.49 | -0.02 | 32.40 | 82.50 |
| Al_GO250P | 145 | 222 | -0.47 | -0.49 | -0.02 | 32.40 | 65.40 |
| Al_GO500P | 145 | 454 | -0.47 | -0.48 | -0.01 | 32.40 | 32.00 |

As per prior literature porosity measurements have also been correlated to the corrosion resistance properties. Coating porosity critically governs corrosion resistance at the microscopic level by controlling electrolyte penetration pathways to the underlying metal^117^. This influence becomes evident when contrasting the effects of large, open pores with small, isolated ones^118^. In systems with large pores, electrolyte and oxygen can circulate freely, facilitating stable passivation and improved resistance to corrosion. In contrast, small pores tend to trap electrolytes and deplete oxygen, promoting aggressive micro-environments for crevice corrosion. Moreover, coatings with low porosity percentage has been shown to maximize barrier performance by restricting ion transport pathways^69,119^. Current results suggest a reduction in porosity percentage with increase in GO concentration for the Al_GO/P test surfaces (Table 3). The observed porosity percentage reduction is from 89.1% through 32%, corresponding to increased GO concentration from 50 to 500 µg/ml. Therefore, surfaces with higher GO concentration with lower porosity percentage should improve corrosion resistance as per earlier literature. The assessment of the corrosion properties for the synthesized substrates is mentioned later in this manuscript.

### 3.2. Assessment of antimicrobial potential of Al_GO/P substrates

#### 3.2.1. Al_GO/P inhibited growth of microbial cells

In a preliminary study, growth inhibition assay was conducted to evaluate the antimicrobial activity of the fabricated Al_GO/P substrates. This is a conventional technique where microbial cells are exposed to substrates for a specific time, followed by growth on nutrient rich medium overnight^63^. The antibacterial and antifungal activities of Al_GO/P substrates (Al_GO50P, Al_GO100P, Al_GO250P, and Al_GO500P) were tested against Gram-positive *S. aureus*, Gram-negative *E. coli*, and fungus *C. albicans*. Bare Al (uncoated Al sheet) and glass slides were used as control surfaces.

Figures 5A, 5B and 5C show the bacterial (Gram-positive *S. aureus* and Gram-negative *E. coli*) and fungal (*C. albicans*) colonies on LB and YPD agar plates, respectively. Control surfaces glass and bare Al displayed rich growth of microbial colonies for both bacterial and fungal species. While significantly low colony growth was observed on Al_GO50P substrate, no bacterial and fungal cell growth were noted on rest of the Al_GO/P substrates. Thus, the growth inhibitory activity of these substrates was conserved across all the microbial species with Al_GO100P, Al_GO250P, and Al_GO500P exhibiting a higher inhibitory potential compared to controls (glass and bare Al), whereas Al_GO50P displayed a moderate inhibitory effect. However, control Al_GO substrates without PEDOT:PSS failed to display significant growth inhibitory activity relative to Al_GO/P test substrates (Figure S5, Supporting Information)

**Figure 5.**
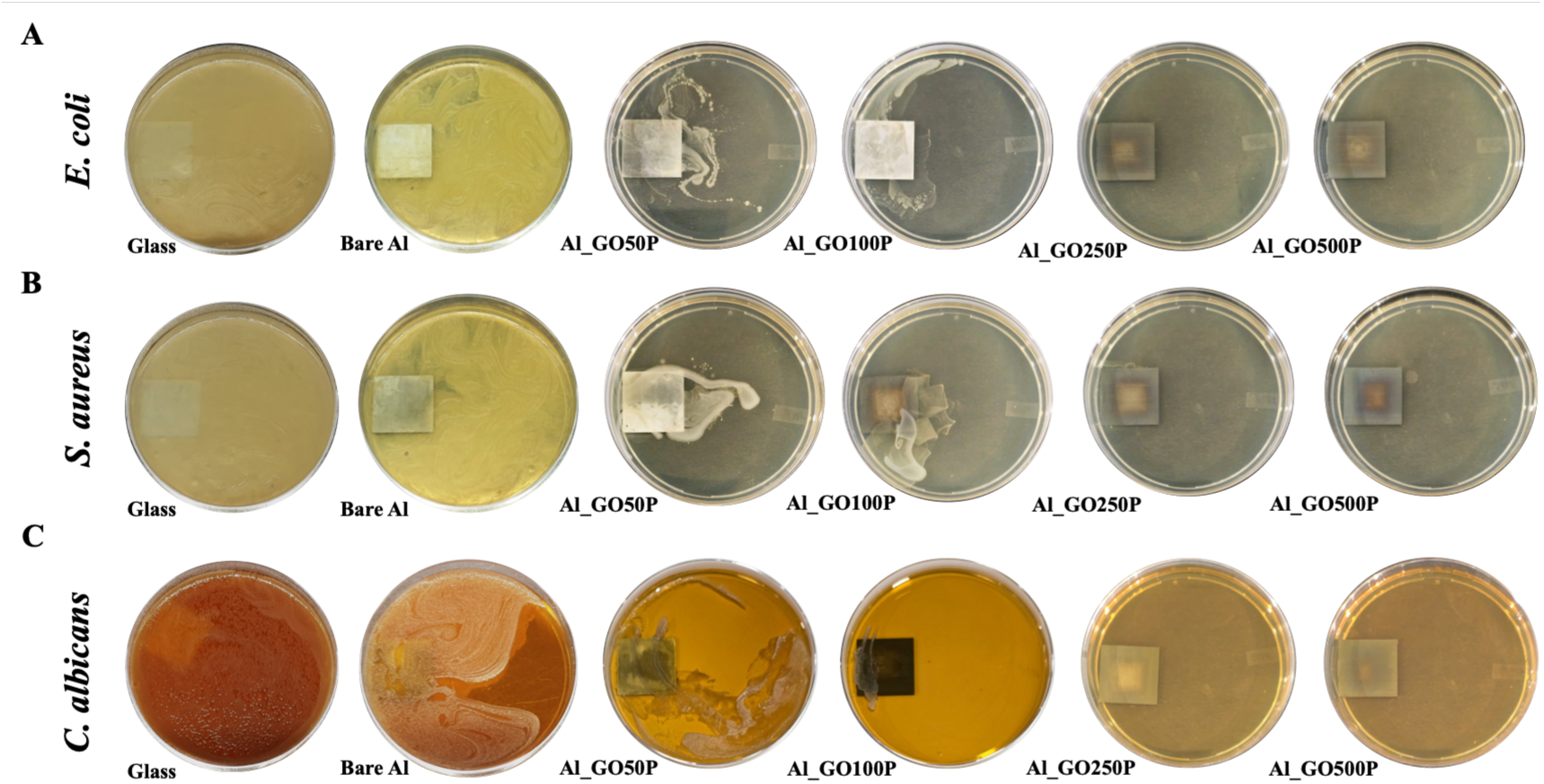
Growth Inhibition Assay of Al_GO/P substrates. After 10 hour of incubation on different control (glass and bare Al) and Al_GO/P substrates, (**A, B**) bacterial (*E. coli, S. aureus*) and (**C**) fungal (*C. albicans*) cells were cultured overnight on LB and YPD agar plates, respectively. The inhibitory roles of the coated substrates were assessed by examining colony growth the following day. Cells cultivated on glass and bare Al surfaces exhibited abundant bacterial and fungal colonies, indicating a lack of antimicrobial potential. In contrast, cells cultured on Al_GO50P presented minimal colony growth while rest of the Al_GO/P substrates demonstrated no visible colony growth.

For a comparative analysis, microbial growth inhibition was performed in presence of pristine GO as shown in Supporting Information (Figure S6A). Relative to the activity of GO, the fabricated substrates were shown to exhibit superior antimicrobial potential towards both bacterial and fungal cells. Besides, evaluation of PEDOT:PSS activity revealed no intrinsic antimicrobial potential over a wide concentration range (1 µg/ml, 50 µg/ml, 100 µg/ml, 250 µg/ml, 500 µg/ml, 1 mg/ml, and 2 mg/ml) (Figure S6B, Supporting Information). Therefore, the relative efficacy of Al_GO/P substrates could be correlated to varied concentrations of GO on Al_GO/P substrates correlating to earlier literatures^120,121^.

Next, cell viability assay was conducted for a quantitative evaluation of the antimicrobial nature of Al_GO/P substrates. Percentage loss of cell viability was calculated via spread plate method following exposure to different control (Glass, bare Al, Al_GO50, Al_GO100, Al_GO250, and Al_GO500) and test (Al_GO50P, Al_GO100P, Al_GO250P, and Al_GO500P) substrates for a period of 0, 5, and 10 hours as described in the Experimental Procedures. For *E. coli* cells (Figure S7A, Supporting Information), at 5 h, the control surfaces showed minimal loss of cell viability, with glass at 1.9%, bare Al at 4.4%, Al_GO50 at 4.1%, Al_GO100 at 9%, Al_GO250 at 17.1%, and Al_GO500 at 20.9%. The corresponding test substrates showed markedly higher loss in cell viability percentage, with Al_GO100P at 44.1%, Al_GO250P at 94.2%, and Al_GO500P at 95.1%, while Al_GO50P caused only 16.3% loss in cellular viability. At 10 h, the loss of cell viability when exposed to controls remained low, with glass at 4.4%, bare Al at 6.4%, Al_GO50 at 8.7%, Al_GO100 at 14.3%, Al_GO250 at 19.6%, and Al_GO500 at 23.4%. The Al_GO/P substrates gave rise to higher loss in cell viability, with Al_GO100P at 65.7%, Al_GO250P at 95.1%, and Al_GO500P at 97%. Al_GO50P induced only a 20.1% loss in cell viability following exposure to *E. coli* cells.

In case of *S. aureus* (Figure S7B, Supporting Information), 5 h exposure caused minor loss in cell viability on control surfaces, with glass at 1.5%, bare Al at 1.7%, Al_GO50 at 2.6%, Al_GO100 at 3.6%, Al_GO250 at 5.1%, and Al_GO500 at 4.8%. In contrast, the Al_GO/P surfaces showed higher loss in cell viability percentage, with Al_GO250P at 96.1% and Al_GO500P at 96.6%, while Al_GO50P at 7.8% and Al_GO100P at 27.9% displayed drastically low loss in cell viability percentages. After 10 h, negligible loss in cell viability were displayed by control substrates: glass at 2.4%, bare Al at 3.3%, Al_GO50 at 4.7%, Al_GO100 at 6.1%, Al_GO250 at 6.3%, and Al_GO500 at 7.0%. The Al_GO/P substrates yielded higher loss in cell viability, with Al_GO100P at 88.1%, Al_GO250P at 97.1%, and Al_GO500P at 98.2%, while Al_GO50P at 15.1% failed to show antimicrobial activity.

For *C. albicans* cells (Figure S7C, Supporting Information), 5 h exposure on control substrates resulted in glass at 1.2% loss, bare Al at 1.8%, Al_GO50 at 1.8%, Al_GO100 at 3.5%, Al_GO250 at 2.5%, and Al_GO500 at 15.4%. Al_GO/P substrates exhibited higher antifungal activity, with Al_GO100P at 50.2%, Al_GO250P at 97.4%, and Al_GO500P at 98.5%, except Al_GO50P at 3.3%. By 10 h, loss of cell viability in presence of glass was 1.9%, bare Al 3.1%, Al_GO50 2.6%, Al_GO100 5.5%, Al_GO250 3.8%, and Al_GO500 44.8%; whereas the Al_GO/P samples produced viability loss of 73.2% (Al_GO100P), 98.3% (Al_GO250P), and 99.4% (Al_GO500P), except Al_GO50P at 6%. Overall, substrates Al_GO250P and Al_GO500P achieved near complete loss (∼100%) of cell viability, closely followed by Al_GO100P, showing superior broad-spectrum antimicrobial activity. However, Al_GO50P remained the least effective. Additionally, cell viability assessments were further validated using standard antimicrobials as positive controls: chloramphenicol (15 µg/ml) against *E. coli* cells, vancomycin (15 µg/ml) against *S. aureus* cells, and fluconazole (15 µg/ml) against *C. albicans* cells; exhibiting complete inhibition of microbial growth at both 5 h and 10 h (Figure S8, Supporting Information) of exposure time.

#### 3.2.2. Al_GO/P substrates demonstrate potent antibiofilm activity

Biofilms are mixed community of microbial population adherent to various biotic and abiotic surfaces^2^. These organized microbial structures play a crucial role in infection dissemination, antibiotic resistance, and may pose a threat to the host’s immune system^122^. As previously reported, the antimicrobial efficacy of GO has been well documented across literature^53,123–125^. In this study, we have evaluated the activity of GO-PEDOT:PSS coated Al (Al_GO/P) surfaces towards mixed species (*C. albicans*, *S. aureus*, and *E. coli*) biofilm formation by examining the progress of biofilm maturation over a period of 10 hours under biofilm stimulating conditions.

Fluorescent stains were used to visualize Gram-positive *S. aureus* (acridine orange, red), Gram-negative *E. coli* (sodium fluorescein, green), and fungus *C. albicans* (calcofluor white, blue). Control surfaces, such as glass (Figure 6A) and bare Al (Figure 6B), demonstrated dense biofilm formation consisting of bacterial cells within intricate fungal hyphae. This indicates robust microbial colonization on these surfaces that provide a favorable environment for biofilm maturation. On the contrary, significant biofilm inhibition was observed on Al_GO/P substrates. A complete disruption of biofilm structure with only a few adherent microbial cells were noted in presence of Al_GO500P (Figure 6F), closely followed by Al_GO250P (Figure 6E), Al_GO100P (Figure 6D). A moderately inhibited biofilm structure was visualized in presence of Al_GO50P (Figure 6C).

**Figure 6.**
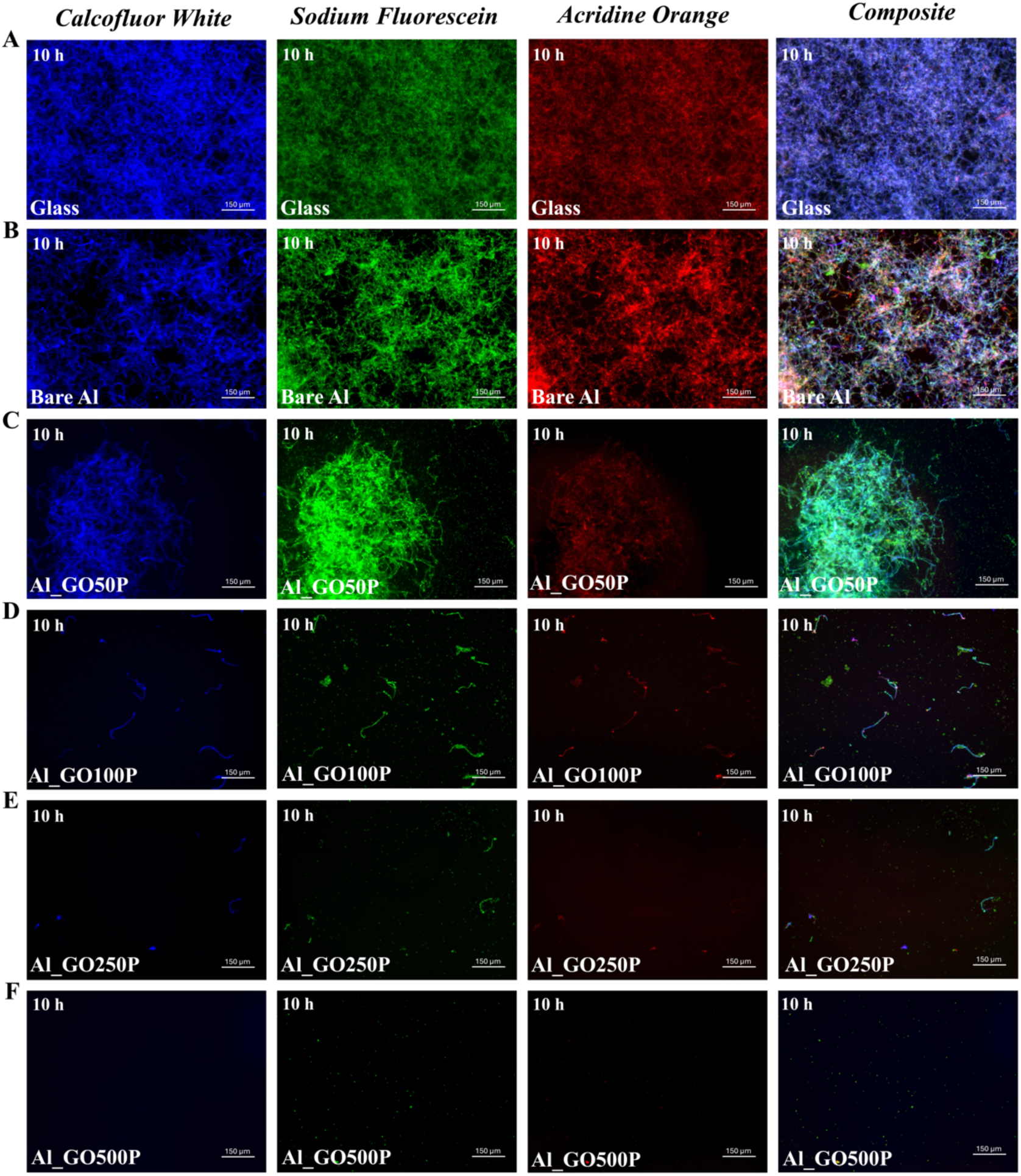
Fluorescence microscopic images of mixed species (*C. albicans*, *S. aureus*, and *E. coli*) biofilm formation (10^7^ CFU). Mixed candida-bacterial suspensions were exposed, (**A, B**)to control surfaces (glass and bare Al), and (**C–F**) to various Al_GO/P substrates. Post-exposure, mixed species biofilm were allowed to grow for 10 hours under biofilm stimulating conditions after which the samples were stained with calcofluor white (*blue*), acridine orange (*red*), and sodium fluorescein (*green*) to visualize *C. albicans*, *S. aureus*, and *E. coli*, respectively. Biofilms were imaged using an in-house fluorescence microscope at 40× magnification.

Further to show the robust activity of Al_GO/P, antibiofilm potential was examined at a higher microbial load of 10^8^ CFU (Figure S9, Supporting Information). The extent of antibiofilm activity of Al_GO/P substrates were comparable when tested with 10^7^ CFU and 10⁸ CFU of mixed microbial population. These findings further support the broad-spectrum antimicrobial nature of the Al_GO/P substrates as shown by growth inhibition assay, while substrates with higher GO concentrations yield a more pronounced biofilm inhibitory effect. For a quantitatively assessment, mean fluorescent intensity (MFI) was calculated from fluorescent images of mixed biofilms both at 10^7^ and 10^8^ CFU using ImageJ software (Figure 6 and Figure S9, Supporting Information). Compared to glass and bare Al controls, Al_GO/P test surfaces exhibited a significant decrease in the MFI for biofilm densities at both the inoculum concentration (10^7^ and 10^8^ CFU), indicating compromised microbial attachment (Figure S10, Supporting Information).

Next, crystal violet (CV) staining technique was performed to quantitate mixed species (10^7^ CFU) biofilm formation over various test substrates. Glass was used as a positive growth control exhibiting 100% biofilm formation. It was observed that biofilm biomass was lowest when exposed to Al_GO500P, decreasing significantly to 14.3% (p < 0.0001), indicating the highest inhibitory activity among the tested Al_GO/P surfaces (Figure S11 and Table S4, Supporting Information). This was followed by Al_GO250P and Al_GO100P, with biofilm biomass reductions to 39.3% and 48.5%, respectively (p < 0.0001 for both). The biofilm biomass was reduced to 91.3% (p > 0.05) in presence of Al_GO50P, indicating the least inhibitory effect (Figure S11 and Table S4, Supporting Information). A similar result was obtained at higher microbial inoculum load at 10^8^ CFU (Figure S12 and Table S4, Supporting Information).

Current results highlights a clear correlation between increasing GO concentration on Al surface and its inhibitory effect on biofilm formation. Al_GO50P appears to have a non-significant (p > 0.05) role in biofilm inhibition, whereas Al_GO/P substrates with higher concentration of GO (100, 250, and 500 µg/ml) could remarkably inhibit biofilm formation. Besides, a non-significant (p > 0.05) inhibition of biofilm formation over bare Al (Figure S11, Figure S12 and Table S4, Supporting information) suggested a superior antibiofilm activity of Al_GO/P surfaces. Additionally in a comparative study, we used pristine GO at corresponding concentrations, which resulted in non-significant (p > 0.05) inhibition of biofilm formation relative to control surface glass (Figure S13 and Table S4, Supporting Information). A pairwise analysis of antibiofilm efficacy between pristine GO and test surfaces indicated significant inhibition (p < 0.0001) induced by Al_GO/P at corresponding concentration (Figure S14, Supporting Information). The quantitative data of the antibiofilm assay for control and test substrates are provided in Table S4, Supporting Information.

Summarizing the results from growth inhibition (Figure 5), cell viability (Figure S7, Supporting Information) and antibiofilm assays; it was observed that Al_GO100P, Al_GO250P, and Al_GO500P substrates exhibit superior antimicrobial activity compared to Al_GO50P and control surfaces. This is possibly due to the suppression of microbial cell growth, colonization, and biofilm maturation in the presence of Al_GO/P substrates^57^. Furthermore, the overall lower porosity percentage of Al_GO/P substrates at increasing GO concentrations from 50-500 µg/ml (Table 3) may have contributed to the enhanced antimicrobial activity by limiting microbial adhesion^116^. Further analysis via SEM is required to visualize the membrane-damaging (biocidal) activity of Al_GO/P, as demonstrated in the following section.

#### 3.2.3. Al_GO/P substrates induced cell membrane damage in mixed species microbial cell population

In a comprehensive study examining the effects of Al_GO/P surfaces on bacterial and fungal cells, SEM was employed to analyze alterations in cellular architecture and membrane integrity. SEM analysis revealed that cells exposed to the control surfaces (glass, bare Al) maintained their normal cellular morphology, appearing healthy and uniform (Figure 7A and 7B). Exposure to Al_GO50P, displayed less severe but notable morphological damage, including fused cells with wrinkled appearance (Figure 7C). In contrast, exposure to the Al_GO100P (Figure 7D), Al_GO250P (Figure 7E), and Al_GO500P (Figure 7F) induced significant damage to cell membrane with complete loss of cellular architecture for all (*C. albicans*, *S. aureus*, and *E. coli*) cell types. The study demonstrated consistent biocidal activity of the test substrates (Al_GO100P, Al_GO250P, Al_GO500P) towards a mixed microbial population consisting of Gram-positive and Gram-negative bacteria, as well as fungi.

**Figure 7.**
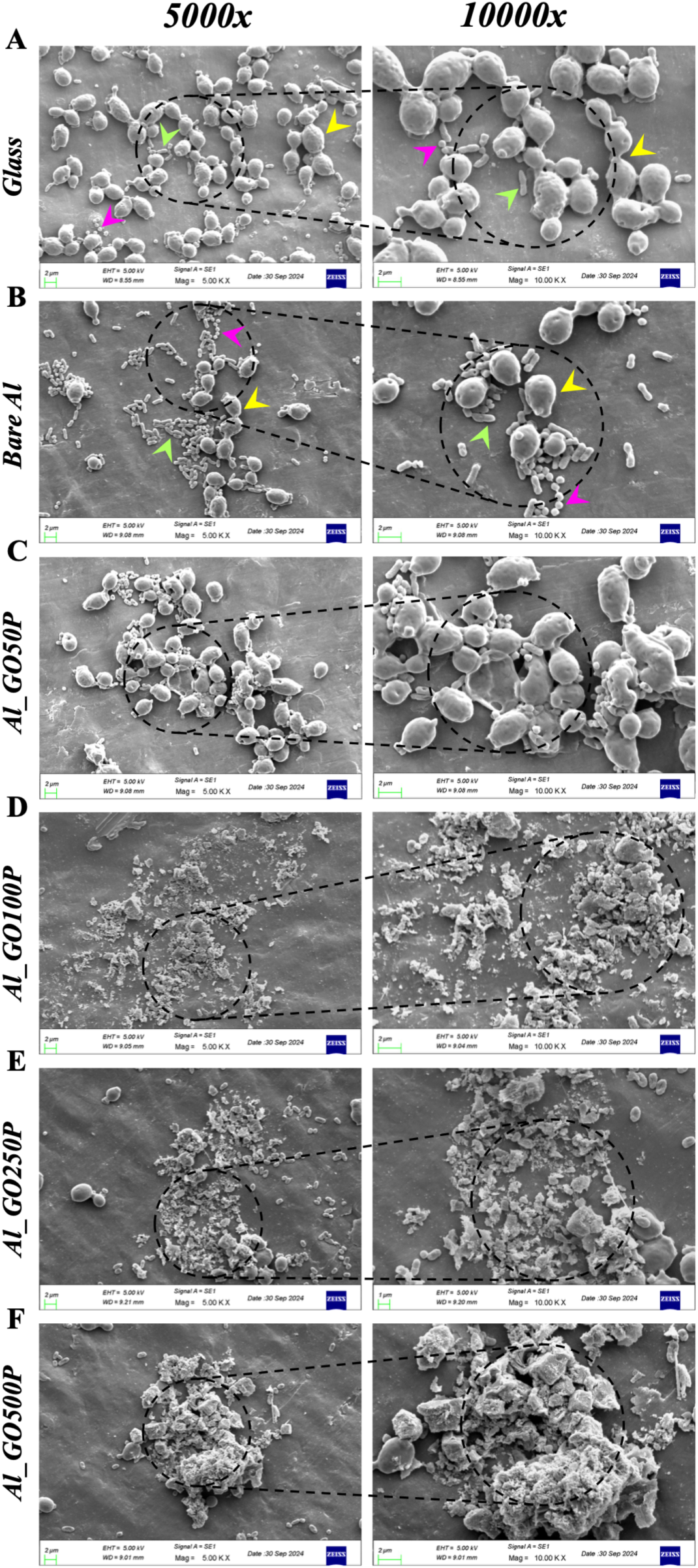
SEM analysis to visualize mixed microbial (*C. albicans*, *S. aureus*, and *E. coli*) cells. Representative SEM micrographs show the morphology of mixed *C. albicans* (*arrowheads, yellow*), *S. aureus* (*arrowheads, pink*), and *E. coli* (*arrowheads, green*) cells after 10 hours of exposure to different control and test substrates: (**A**) Glass, (**B**) Bare Al, (**C**) Al_GO50P, (**D**) Al_GO100P, (**E**) Al_GO250P, and (**F**) Al_GO500P. Images captured at 5000× and 10,000× magnifications (*dashed circles, cyan*) revealed extensive membrane damage, and complete loss of cellular architecture in microbial cells exposed to Al_GO100P, Al_GO250P and Al_GO500P, compared to the intact morphology observed on control surfaces (Glass, Bare Al). Exposure to Al_GO50P resulted in minor morphological alterations, such as fused and wrinkled cell structures.

The enhanced biocidal activity of Al_GO/P may be correlated to superior surface hydrophobicity at different GO concentrations relative to bare Al (Table 2 and Figure S3, Supporting Information). This may have facilitated sustained interactions between the microbial cell wall and the Al_GO/P surfaces, thereby compromising membrane integrity^126^. Besides, we cannot exclude the possibility of electron transfer events occurring in a conductive background of the Al_GO/P substrates, which may lead to the generation of reactive oxygen species (ROS), as shown in previous literature^57,63^. Next, we investigated ROS generation at the substrate interface and intracellularly to further validate the underlying biocidal mechanism of the Al_GO/P substrates.

#### 3.2.4. Al_GO/P substrates induce intracellular and surface ROS production

The catalytic generation of ROS at metal-environment interfaces is a well-characterized process, mediated by physicochemical interactions that promote oxidative stress conditions^127,128^. These interactions involve complex mechanisms, including redox reactions and surface modifications, which could mediate the transfer of electrons and lead to the formation of ROS^63^. The production of ROS can have significant implications for both the metal surfaces and the biological systems they interact with, potentially affecting cellular responses^127^. In this work, ROS production was evaluated using the indicator dye 2’,7’-dichlorodihydrofluorescein diacetate (H_2_DCF-DA) after exposing mixed (bacterial and fungal) microbial cells to various Al_GO/P substrates. This dye is effective in detecting a wide range of ROS, including hydrogen peroxide (H_2_O_2_), hydroxyl radicals (•OH), and other reactive species such as carbonate radicals and nitric oxide^129^. When ROS are present, the initially non-fluorescent H_2_DCF-DA is transformed into the highly fluorescent compound 2’,7’-dichlorofluorescein (DCF+), which emits a green signal observable through fluorescence microscopy.

In addition, dihydroethidium (DHE) is also utilized for ROS detection. DHE displays blue fluorescence in the cell cytoplasm; however, upon reacting with superoxide anions, it is oxidized to form 2-hydroxyethidium, which intercalates into DNA and exhibits red fluorescence (DHE+). This makes DHE particularly useful for detecting cytosolic superoxide (O_2_•⁻)^129^. It can also undergo nonspecific oxidation by other radicals like peroxynitrite (ONOO−) and hydroxyl radicals (•OH). In this work, H_2_DCFDA dye was employed to assess the overall oxidative stress levels in mixed *Candida*-bacterial cell suspensions because it broadly detects multiple ROS species, including H_2_O_2_, •OH, and others, after deacetylation to DCFH, yielding green-fluorescent DCF⁺ suitable for overall oxidative stress assessment across diverse cell types. Additionally, DHE was utilized specifically for mixed bacterial cell suspensions (*S. aureus* and *E. coli*) due to its higher selectivity for superoxide (O_2_•⁻), the primary ROS generated in prokaryotic cells.

We observed that Al_GO/P substrates, especially Al_GO500P, Al_GO250P, Al_GO100P exhibited a significantly higher number of DCF+ signals indicative of elevated ROS relative to control surfaces both intracellularly (Figure 8A) and at the substrate surface (Figure 8C) in mixed pathogen populations consisting of *S. aureus*, *E. coli*, and *C. albicans*. This was closely followed by Al_GO50P displaying relatively lower number of DCF+ fluorescent cells. The control Al surface showed minimal DCF+ signal, while the glass surface displayed no detectable DCF+ (green) signal at all. The elevated ROS levels may directly correlate to an electron transfer event from the microbial cells to the GO functional groups over conductive substrates causing surface ROS production. Elevated intracellular ROS production is indicative of cell death and the activation of related cell signaling pathways^130^.

**Figure 8.**
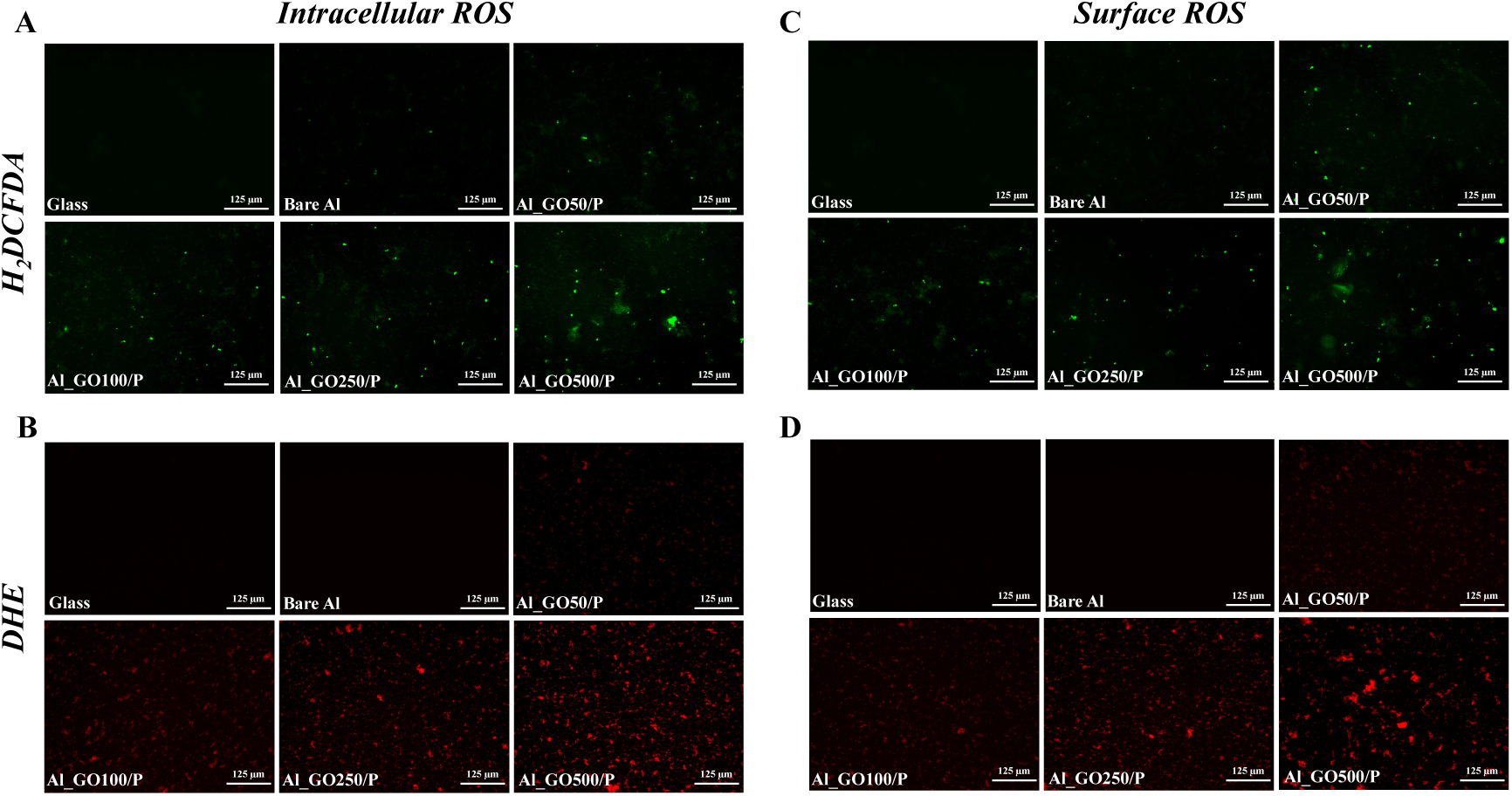
Evaluation of (A, B) intracellular, and (C, D) surface ROS production on Al_GO/P substrates using H_2_DCFDA and DHE dye. Following exposure to various test and control surfaces **(A)** Mixed microbial cell population (*C. albicans*, *S. aureus*, and *E. coli*) were stained with H_2_DCFDA dye (green cells), and **(B)** mixed bacterial cells consisting of *S. aureus* and *E. coli* were stained with DHE dye (red cells), for detection of intracellular ROS production. No ROS production was observed on glass, and only minimal signal was detected on Al relative to the test substrates. Abundant fluorescent signals correlating to intracellular ROS generation were observed for the bacterial and fungal cells exposed to Al_GO500P, Al_GO250P, and Al_GO100P surfaces using H_2_DCFDA (green) and DHE (red) fluorescent dyes. Al_GO50P induced relatively lesser number of fluorescent cells; Detection of ROS production on the surface of Al_GO/P substrates after a 10-hour of exposure to **(C)** mixed population of cells (*C. albicans*, *S. aureus*, and *E. coli*) using H2DCFDA dye (green fluorescence), and **(D)** mixed bacterial cells (*S. aureus* and *E. coli*) with DHE dye (red fluorescence). ROS indicator dyes H_2_DCFDA and DHE were applied on different test and control surfaces. No ROS production was noticed on the surface of Glass and Al relative to test substrates. Varied levels of abundant fluorescent signals correlating to ROS generation were observed on the surface of Al_GO500P, Al_GO250P, and Al_GO100P as revealed by H_2_DCFDA (green) and DHE (red) stains. Relatively mild signal was detected on the surface of Al_GO50P. Cells were imaged using in-house fluorescence microscope at 40× magnification.

We further examined the presence of cytosolic superoxide in a mixed bacterial population both intracellularly (Figure 8B) and at the substrate interface (Figure 8D). Similar to our observation in Figure 8A and 8C, control surfaces such as glass and bare aluminium, did not exhibit any DHE+ (red) cells, indicating a lack of ROS production overall. Conversely, varying numbers of fluorescent DHE+ cells were observed on different Al_GO/P substrates. Notably, Al_GO500P, Al_GO250P, and Al_GO100P induced a higher population of DHE+ cells than Al_GO50P. It is plausible that extending the exposure time on the substrate surface may lead to a greater number of ROS-positive cells. ROS production was quantified as average fluorescence intensity per cell using ImageJ analysis of H_2_DCFDA and DHE stained images. For H_2_DCFDA staining (Figure S15A, Supporting Information), bare Al exhibited baseline intracellular and surface ROS levels. Al_GO/P substrates showed concentration-dependent ROS elevation with increasing GO concentration, demonstrating significantly higher intracellular and surface ROS compared to bare Al. Similarly, DHE staining (Figure S15B,

Supporting Information) followed this trend, with moderate to high intracellular and surface ROS generation across Al_GO/P substrates. Notably, bare Al showed no detectable ROS-positive cells with DHE staining. Previous studies have highlighted the role of ROS in the broad-spectrum antimicrobial activity of graphene and its derivatives^57,131^. In fact, oxidative stress can lead to oxidation of proteins, lipids, and nucleic acids in microbial cells, resulting in irreversible membrane damage and cell death^132^. The pronounced microbiocidal activity observed for Al_GO100P, Al_GO250P, and Al_GO500P substrates may be attributed to the conductivity of these substrates (Table 2), possibly facilitating electron transfer from the microbial cell onto the GO surface, enhancing ROS generation. Besides higher surface hydrophobicity values for Al_GO/P substrates (Table 2), may have promoted a stable microbial interaction causing contact mediated membrane damage and ROS induced stress. Although Al_GO50P exhibited higher conductivity, its antimicrobial performance was less, likely due to its lower surface hydrophobicity, causing an attenuated microbial cell interaction with its surface.

### 3.3. Assessment of anticorrosion potential of Al_GO/P substrates

#### 3.3.1. Al_GO/P substrates exhibit biocorrosion resistance

The corrosion rate of the fabricated Al_GO/P substrates was primarily assessed in two designated immersion systems (abiotic and biotic), for a comparative analysis^133^. The abiotic system consisting of growth media and antibiotics allowed to evaluate the chemical corrosion properties of Al_GO/P surfaces, free from biological contamination. Concurrent assessment of the biotic system in presence of microbial cells, monitored microbiologically influenced corrosion (MIC) potential of Al_GO/P substrates. This dual approach not only provides a comprehensive understanding of the corrosion behaviour of the fabricated Al_GO/P substrates but also highlights their potential effectiveness in real-world applications, where both chemical (abiotic) and biological (biotic) factors influence corrosion rates^134^. Additional control materials such as cold-rolled steel (CR steel), which is recognized as a highly corrosive material ^135^, and bare Al were included for the study. Besides, the non-polymeric Al_GO substrates were included for a comparative analyses. Although both chemical corrosion and MIC involve material degradation, chemical corrosion generally proceeds faster and more aggressively due to its direct chemical interactions with the metal surface, especially in high-temperature or harsh environments^136^. Whereas MIC, also an electrochemical process, relies on the activity of microorganisms and their metabolites to promote corrosion, often under specific conditions^25^. While biofilms can occasionally form protective layers under specific conditions^137,138^, they typically accelerate microbiologically influenced corrosion (MIC) through multiple mechanisms including differential aeration cells, production of corrosive metabolic byproducts, and extracellular polymeric substance (EPS) concentration effects, to name a few^25^.

Figure 9 illustrates the corrosion rates of various substrates after 45 days of immersion in the studied systems, highlighting the effects of both abiotic and biotic conditions. In the abiotic system, the corrosion rates were measured as follows: 1528.3 ± 0.1 µm/year for CR steel (Figure 9Ai), 94.3 ± 0.8 µm/year for aluminium (Figure 9Aii); 64.3 ± 1.8 µm/year for Al_GO50 and 38.8 ± 0.5 µm/year for Al_GO50P (Figures 9Aiii and 9Aiv); 53.0 ± 9.5 µm/year for Al_GO100 and 36.9 ± 0.5 µm/year for Al_GO100P (Figures 9Av and 9Avi); 52.7 ± 3.4 µm/year for Al_GO250 and 35.4 ± 1.0 µm/year for Al_GO250P (Figures 9Avii and 9Aviii); 51.1 ± 7.6 µm/year for Al_GO500 and 15.3 ± 4.1 µm/year for Al_GO500P (Figures 9Aix and 9Ax). In contrast, the corrosion rates in the biotic system were recorded as follows: 2674.2 ± 0.3 µm/year for CR steel (Figure 9Ai), 74.2 ± 0.8 µm/year for aluminium (Figure 9Aii); 75.3 ± 2.9 µm/year for Al_GO50 and 31.6 ± 0.5 µm/year for Al_GO50P (Figures 9Aiii and 9Aiv); 65.4 ± 3.7 µm/year for Al_GO100 and 17.6 ± 0.6 µm/year for Al_GO100P (Figures 9Av and 9Avi); 57.5 ± 0.8 µm/year for Al_GO250 and 1.0 ± 0.8 µm/year for Al_GO250P (Figures 9Avii and 9Aviii); 49.0 ± 3.0 µm/year for Al_GO500 and 0.6 ± 0.3 µm/year for Al_GO500P (Figures 9Aix and 9Ax). It was noted that, in general, the immersion systems lacking microbial presence (abiotic) tended to display higher corrosion rates than the corresponding biotic conditions in agreement with prior literature ^136,139,140^. However, a known corrosive material CR steel did not exhibit this trend, but displayed highest overall corrosion rates. These observations align with the understanding that chemical corrosion often occurs at a faster and higher rate than MIC, which is primarily driven by electrochemical reactions in aggressive environments^136,139,140^.

**Figure 9.**
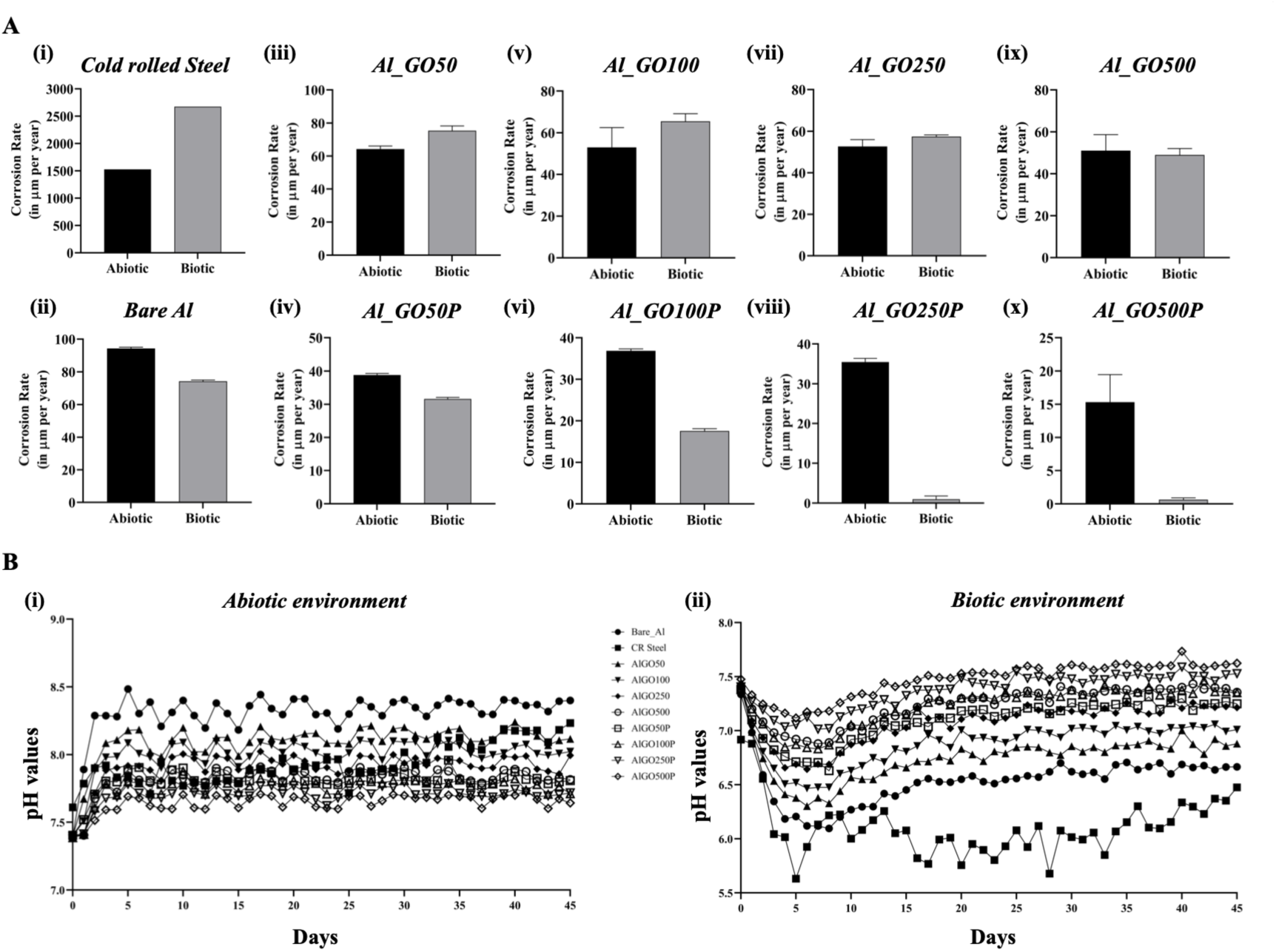
Corrosion rates and pH changes of various control and Al_GO/P substrates over 45 days of exposure to abiotic and biotic environments. **(A)** Corrosion rate(s) of different controls and test substrates after 45 days of exposure to abiotic and biotic immersion environments. Error bars represent ±1 standard deviation (SD) from mean of three independent biological replicates (n=3). **(B)** pH values of **(i)** abiotic and **(ii)** biotic immersion environments in presence of different controls, Al_GO, and Al_GO/P substrates over a period of 45 days.

Corrosion rates for the Al_GO/P substrates immersed in the abiotic system corresponded to a moderate corrosion intensity as defined by the NACE Standard RP-07-75^82^.

The substrates in biotic immersion systems showed lower corrosion rates, reflecting the variability of MIC, which can be influenced by factors such as the type of microorganisms present and their metabolic by products ^24,25^. Further, in the biotic system, Al_GO/P substrates, particularly Al_GO500P and Al_GO250P, followed by Al_GO100P and Al_GO50P, exhibited significantly lower corrosion rates compared to the control surfaces (bare Al and CR steel), indicating enhanced resistance to MIC. This may be explained by a reduction in porosity percentage with increase in GO concentration for the Al_GO/P test surfaces, which leads to reduced microbial adhesion, enhanced barrier characteristics and improved corrosion resistance ^141–144^. Moreover, Al_GO/P displayed lower corrosion rate compared to the corresponding non-polymeric Al_GO substrates (Figures 9Aiii-x) possibly due to higher stability and reduced leaching as evaluated by ICP-MS later.

Additionally, FESEM analysis revealed distinct morphological alterations in control samples (bare Al, CR steel, and Al_GO) compared to the synthesized test Al_GO/P substrates following immersion under abiotic (Figure S16 and S17, Supporting Information) and biotic (Figure S18 and S19, Supporting Information) environments. These observations align with the corrosion rate analyses (Figure 9A), underscoring the superior anticorrosion performance of the GO-PEDOT:PSS functionalized Al system. In contrast, uncoated bare Al, CR steel and Al_GO surfaces exhibited pronounced localized pitting and deep corrosion features, suggesting their susceptibility towards chemical corrosion as well as MIC. For visual comparison, the optical images of the control and test substrates captured before and after immersion in abiotic and biotic environments are presented in Figure S20 (Supporting Information).

Additionally, the pH variation in both abiotic and biotic immersion systems was monitored over a period of 45 days. The pH of each immersed sample was recorded daily, and the average value was determined from two measurements to minimize experimental error. Under abiotic conditions (Figure 9Bi), CR steel displayed a pH rise to alkaline values (∼7.4 to ∼8.5) over the 45 day immersion time, possibly due to release to Fe^2+^. Bare Al demonstrated a pH rise from ∼7.4 to 8.5 during the early phase (0-7 days), possibly driven by Al dissolution and Al(OH)_3_ hydrolysis generating OH⁻ ions^145^, followed by mid-late phase pH stabilization (7-45 days) possibly due to accumulation of corrosion products and surface passivation^146^. Likewise, Al_GO surfaces displayed pH rise to an alkaline value, may be due to compromised barrier activity of GO coating in the absence of PEDOT:PSS^147^. Al_GO/P substrates maintained nearly constant pH (7.4) under abiotic immersion condition, likely due to the superior barrier properties of GO-PEDOT:PSS and improved adhesion on Al surface.

In biotic conditions (Figure 9Bii), CR steel demonstrated an sharp pH drop to an acidic range (6–5.5) over the 45 day immersion period. Similarly, bare Al, Al_GO50, Al_GO100, Al_GO250, and Al_GO500 gave rise to overall acidic pH likely due to microbial colonization and release of microbial products during the immersion period^148,149^. In contrast, Al_GO/P surfaces induced a neutral pH (7.0∼7.5) condition in the immersion medium throughout, possibly due to inhibition of microbial colonization and strong antibiofilm effect (Figure 6).

Previous studies have demonstrated that antimicrobials can serve as green corrosion inhibitors. For instance, Obot and coworkers in 2008 reported fluconazole (antifungal) as an excellent corrosion inhibitor for aluminium in acidic medium (0.1M HCl)^150^. Similarly, Ayoola *et al.* reported the corrosion protection of chloramphenicol (antibiotic) on A315 mild steel in 0.1 M HCl^151^, while Alamry *et al.* showed the inhibitory capacity of ampicillin (antibiotic) on mild steel in 5% HCl^152^. However, the anticorrosion performance noted for Al_GO/P substrates in the abiotic immersion system cannot be attributed to such inhibitory roles of antimicrobials, since pH analysis (Figure 9Bi) confirmed neutral conditions. Further the pH environment for the control surfaces were alkaline over 45 day immersion period, ruling out corrosion inhibitory potential for the antimicrobials used for the study. Therefore, in the current work, the antimicrobials used in the abiotic system, did not influence the observed corrosion rates for the test substrates.

Next, ICP–MS analysis was conducted to evaluate the long-term stability of the substrates after 45 days of immersion under abiotic and biotic conditions (Table S5, Supporting Information). Under abiotic conditions, Fe^2+^ ions were detected at a concentration of 3 ppm for the known corrosive control, CR steel. The release of Al^3+^ ions was measured as 0.82 ppm, 0.02 ppm, 0.26 ppm, 0.88 ppm, and 1.30 ppm for bare Al, Al_GO50, Al_GO100, Al_GO250, and Al_GO500, respectively. In comparison, the Al_GO/P substrates demonstrated markedly lower Al^3+^ ion release, with concentrations of 0.03 ppm (Al_GO50P), 0.09 ppm (Al_GO100P), 0.33 ppm (Al_GO250P), and 0.43 ppm (Al_GO500P), indicating superior corrosion protection.

Under biotic conditions, Fe^2+^ ion concentration was comparatively lower (2 ppm) for CR steel, indicating moderated corrosion dynamics under microbial environment. The corresponding Al^3+^ ion release values were 0.75 ppm, 0.05 ppm, 0.12 ppm, 1.04 ppm, and 1.56 ppm for bare Al, Al_GO50, Al_GO100, Al_GO250, and Al_GO500, respectively. Similar to abiotic environment, the Al_GO/P substrates showed reduced ionic leaching, with Al^3+^ ion concentrations of 0.01 ppm (Al_GO50P), 0.04 ppm (Al_GO100P), 0.45 ppm (Al_GO250P), and 0.54 ppm (Al_GO500P).

Thus, the observed overall suppression of ion leaching in the Al_GO/P coatings under both abiotic and biotic conditions may be attributed to improved surface integrity arising from molecular-level interactions between GO and PEDOT:PSS^153^. The oxygenated functional groups on GO (hydroxyl, epoxy, carbonyl, and carboxyl) provide negatively charged sites that electrostatically associate with the positively charged PEDOT backbone, while simultaneously engaging in hydrogen bonding and dipolar interactions with the sulfonate-rich PSS domains. In parallel, the sulfonate oxygens of PSS establish additional electrostatic and hydrogen-bonding contacts with these GO functionalities, creating a dense interfacial network that possibly “locks” the GO sheets into the PEDOT:PSS matrix and suppresses GO leaching^154^. Taken together, these noncovalent interactions may stabilize the GO carbon lattice within the GO–PEDOT:PSS hybrid coating, such that PEDOT:PSS primarily serves as an adhesive interlayer that enhances GO adhesion at the molecular level rather than acting as an independent functional component.

Collectively, these results underscore the long-term stability of GO-PEDOT:PSS hybrid coated Al system in improving corrosion resistance under chemical and bio-corrosive environments. Improved GO adhesion due to PEDOT:PSS and higher surface hydrophobicity of Al_GO/P substrates may further enhance a stable interaction with GO, inducing contact-killing mechanisms (GO-edge effect, ROS production) more efficiently. These findings are further supported by results from the antibiofilm assays (Figure 6, Figure S11 and Table S4, Supporting Information), SEM analysis (Figure 7), ROS generation (Figure 8), and corrosion analysis (Figure 9 and Table S5, Supporting Information). The reduced surface leaching and superior corrosion resistance properties of Al_GO/P surfaces, led us to evaluate the biocompatibility of the test substrates.

### 3.4. Cytotoxicity assessment: Al_GO50P and Al_GO100P demonstrate superior biocompatibility

MTT assay is a widely utilized method for assessing cytotoxicity, while relying on the metabolic activity of living cells^86,155^. This method involves the reduction of the yellow tetrazolium salt, MTT (3-(4,5-dimethylthiazol-2-yl)-2,5-diphenyltetrazolium bromide), to an insoluble purple formazan product by mitochondrial enzymes present in metabolically active cells. The amount of formazan produced is directly proportional to the number of viable cells, allowing for quantification through absorbance measurements at 570 nm.

For biocompatibility assessment, Al_GO1000P, featuring a GO concentration of 1 mg/ml, was included for comparative analysis with the test Al_GO/P substrates^57^. We conducted MTT assay with material extracts as well as direct exposure to the substrates to gain a comprehensive understanding of their effects on HEK293T kidney cell lines^156,157^. Given that the release of nanomaterials from antimicrobial surfaces poses a significant concern, potentially leading to systemic toxicity affecting various organs and functional systems, it was crucial to investigate this further^27^. An extract test was performed on controls (glass, bare Al, and Al_GO) and test Al_GO/P substrates to determine leaching-related cytotoxicity of HEK293T cells ^87^. The quantitative values used to plot the graphs are tabulated in Table S6A and S6B (Supporting Information) for indirect (extract) and direct contact assay, respectively.

The assessment of HEK293T cell viability after exposure to the Al_GO/P material extracts (indirect contact assay) (Al_GO50P, Al_GO100P, Al_GO250P, Al_GO500P, Al_GO1000P) revealed non-significant (p > 0.05) impact on cell growth or viability cell population when compared to the control HEK293T cells in growth media alone (Figure 10A and Table S6A, Supporting Information). No changes in HEK293T cell viability were observed upon indirect contact with the glass surface, as anticipated. In contrast, extracts from bare aluminum resulted in a notable decrease in HEK293T cell viability, dropping to 71.9% (p ≤ 0.0001) and 72.6% (p ≤ 0.0001) at 24 and 48 hours, respectively; compared to unexposed control cells, which exhibited a viability of 100% at similar time points. As shown by previous literatures, systemic toxicity concern arise from the ability of bare aluminum to release ions (Al^3+^) or particles that can disrupt cellular function, leading to reduced viability and potential long-term health effects^158,159^. Current results suggested that fabricated Al_GO/P substrates possess a stable GO-PEDOT:PSS coating, effectively minimizing the risk of harmful leaching and subsequent toxicity.

**Figure 10.**
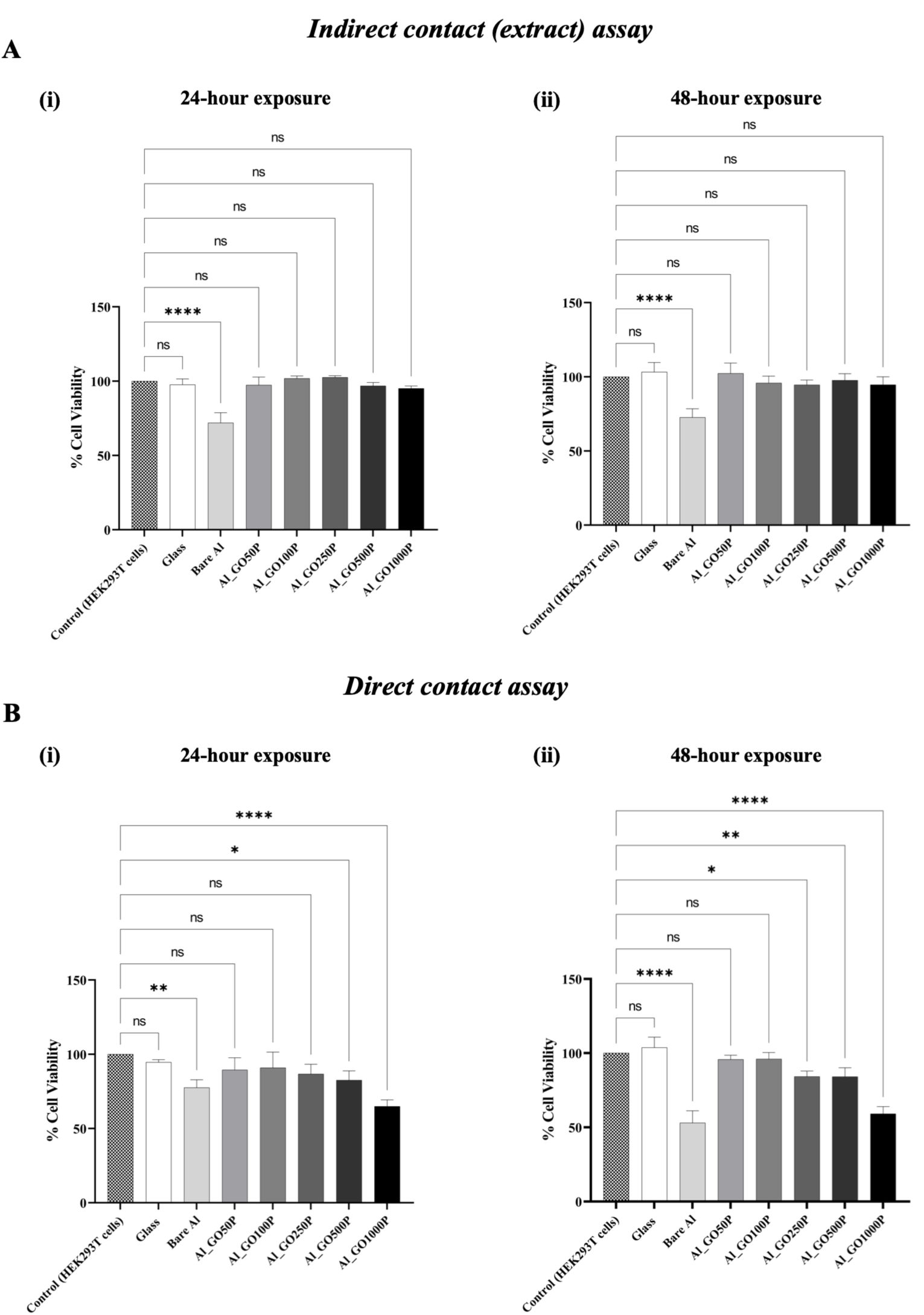
Cytotoxicity evaluation of control and Al_GO/P substrates. Comparison of cell viability, assessed by MTT method in the DMEM culture medium, after exposure to (**A**) Al_GO/P sample extracts by indirect contact assay and (**B**) after direct contact with the Al_GO/P samples; at (**i**) 24-hour and (**ii**) 48-hour. Data are expressed as % values of result obtained in comparison to control cells (HEK293T) not exposed to materials or material extracts. Results from indirect contact assay showed Al_GO/P substrates maintained optimal biocompatibility compared to bare Al over 48 hours. While direct contact assay indicated high cell viability for Al_GO50P and Al_GO100P, with Al_GO250P showing minor viability changes at 48 hours, while significant decreases were observed for Al_GO500P and Al_GO1000P over 48 hours. Error bars represent ±1 standard deviation (SD) from mean for three independent biological replicates (n=3). Statistical significance was assessed using one-way ANOVA; specific p-values of <0.05, <0.01, <0.001, and <0.0001 were indicated as *,**,***, and ****, respectively.

Additionally, direct contact assay was conducted to validate uses of Al_GO/P substrates for potential *in vivo* biomedical applications. HEK293T cell viability data (Figure 10B and Table S6B, Supporting Information) indicate that exposure to Al_GO50P (89.5% at 24 h and 95.8% at 48 h; p > 0.05) and Al_GO100P (90.9% at 24 h and 96.1% at 48 h; p > 0.05) resulted in no significant change in HEK293T cell viability compared to the control. Al_GO250P (86.7% at 24 h, p > 0.05; 84.3% at 48 h, p = 0.0116) caused a non-significant inhibition of cell viability percentage over 24 h, with a mild inhibition of viable cells at 48 h. In contrast, Al_GO500P (82.5% at 24 h, p = 0.0234; 84.1% at 48 h, p = 0.0108) showed a moderate reduction in cell viability over 24-48 hour. Notably, Al_GO1000P (64.9% at 24 h, p < 0.0001; 59.1% at 48 h, p < 0.0001) caused a significant decrease in HEK293T cell viability. HEK293T cells exposed to bare Al demonstrated similar loss in cell viability percentage, 77.5% at 24 hours (p = 0.0038) and 53% at 48 hours (p < 0.0001). As expected there was no alteration in the viability of HEK293T cells in direct contact with the glass surface. These results collectively highlight the superior biocompatibility of Al_GO50P, Al_GO100P, and Al_GO250P underscoring their potential for possible biomedical applications.

As a comparative study, we additionally synthesized Al_GO control substrates without PEDOT:PSS and evaluated their cytotoxicity via extract (indirect) and direct contact assays. Results revealed that Al_GO substrates exhibited significantly low cell viability percentages relative to Al_GO/P substrates (Figure S21, Supporting Information), indicating that the incorporation of PEDOT:PSS in Al_GO/P system displayed improved biocompatibility. We further examined the cell viability of HEK293T cells exposed to pristine GO and PEDOT:PSS at corresponding test concentrations. The cell viability percentage(s) decreased with increasing concentrations of pristine GO (50 µg/ml, 100 µg/ml, 250 µg/ml, 500 µg/ml, and 1000 µg/ml) and PEDOT:PSS (1 µg/ml, 50 µg/ml, 100 µg/ml, 250 µg/ml, 500 µg/ml, 1 mg/ml, and 2 mg/ml) (Figures S22 and S23, Supporting Information) when exposed to HEK293T cells for a period of 24 and 48 hours. Notably, PEDOT:PSS at a concentration used to prepare the hybrid coating in this study (1 µg/ml) led to nonsignificant alteration to HEK293T cell viability.

Current findings confirm that Al_GO/P surfaces do not exhibit elevated leaching, as evidenced by the extract (indirect) assay (Figure 10A), corrosion rate measurements (Figure 9A) and ICP-MS analysis (Table S5, Supporting Information). HEK293T cells exposed to extracts (indirect contact assay) from all Al_GO/P test surfaces maintained optimal viability comparable to the untreated controls, whereas extracts from bare aluminium caused a significant (p < 0.0001) reduction in cell viability. The corrosion resistance properties of the Al_GO/P substrates demonstrated in this study likely attenuated the cytotoxic effects associated with ion leaching, as reported in prior literature^160,161^. Additionally, direct contact assays displayed healthy, viable HEK293T cells in presence of Al_GO50P and Al_GO100P surfaces, with no statistically significant differences from the controls. In contrast, direct exposure to bare Al (p < 0.0001), Al_GO500P (p = 0.0108), and Al_GO1000P (p < 0.0001) surfaces resulted in significant inhibition of HEK293T cell viability over a period of 48 hours. Al_GO250P did not influence cell viability over 24 hour of exposure time, however induced minor alteration in viability (p < 0.05) after 48 hour of direct exposure. This indicates the possibly use of Al_GO250P for short-term biomedical applications. While Al_GO500P exhibited remarkable antibiofilm properties (Figure 6F), the significant loss of HEK293T cell viability suggest potential cytotoxicity concerns in direct-contact mediated biomedical applications. Therefore, Al_GO500P may be more suitable towards *in vitro* or exterior uses where robust antimicrobial activity is essential. Although, Al_GO50P demonstrated excellent biocompatibility, it lacked significant antimicrobial, and antibiofilm efficacy, indicating a need for further optimization to enhance its performance. Overall, substrates Al_GO100P and Al_GO250P achieved a desirable balance, combining high biocompatibility and effective antibiofilm activity, and thus stand out as promising candidates for *in vivo* biomedical device applications where cellular compatibility and biofouling resistance are paramount.

In summary, the current results signify that GO-PEDOT:PSS hybrid coated Al demonstrates enhanced antibiofouling, biocorrosion resistance, and biocompatibility properties. Effective GO-PEDOT:PSS hybrid coating on Al surfaces might provide necessary surface integrity enhancing growth inhibition, loss of microbial cell viability, and biocidal activity, upon exposure to substrate surface. The broad-spectrum antimicrobial activity was supported by significant mitigation of mixed-species biofilm formation when exposed to Al_GO/P surfaces. Moreover, the surface hydrophobicity observed on the test Al_GO/P surface might play a distinct role by stabilizing microbial interaction increasing efficient exposure. Further, conductive Al_GO/P surfaces may facilitate electron transfer from microbial cells to the GO functional groups that leads to ROS production. Additional investigations highlight the superior corrosion resistance properties of Al_GO/P substrates through lower corrosion rates, maintenance of neutral pH, and reduced Al^3+^ ion leaching (ICP-MS). Biocompatibility tests highlighted the potential of Al_GO/P to be used as *in vivo* biomedical implant material. Notably, comparative studies with Al_GO substrates lacking PEDOT:PSS polymer exhibited markedly reduced antimicrobial activity, biocorrosion resistance, and biocompatibility compared to Al_GO/P surfaces. Although, PEDOT:PSS alone demonstrated no intrinsic antimicrobial activity in our experiments (Figure S6B, Supporting Information), ICP-MS analysis confirmed the critical contribution to reduced ionic leaching and long-term stability of the Al_GO/P substrates relative to Al_GO surfaces (Table S5, Supporting Information).

## 4. Conclusion

Biofilms are implicated in 60–70% of nosocomial infections and significantly increase the risk of medical device-associated complications, leading to higher infection rates, morbidity, and mortality in clinical settings^162^. The resilience of biofilms to conventional antibiotics underscores the urgency for innovative surface technologies that can effectively prevent microbial colonization and reduce HAIs. Considering these challenges, there is a critical need to design and systematically assess novel materials possessing broad-spectrum antifouling capabilities, while minimizing cytotoxicity and biocorrosion.

Addressing the phenomenon of MIC and understanding the basis of material degradation becomes essential for safe application in human environment. This represents an intersection where biotic and abiotic factors converge to accelerate material degradation through complex bio-electrochemical processes^24,136^. Apart from electrochemical corrosion process, microbial metabolites containing organic acids, carboxyl groups, lower the pH of the metallic environment, accelerating corrosion^149^. Prior research demonstrates that biofilm formation on metal surfaces can alter the microenvironment and cause an increased corrosion rate^163,164^. Extensive scientific literature demonstrates that metal corrosion results from synergistic interactions of multiple mechanisms rather than a single pathway, with biocorrosion operating in tandem with chemical corrosion processes to accelerate material degradation^25,163–165^. Moreover, aluminum alloys are particularly susceptible to MIC due to their inherent tendency to form passive oxide layers, which can be compromised by microbial metabolites leading to localized pitting and crevice corrosion^148^. Although PEDOT:PSS lacks intrinsic antimicrobial activity, it was incorporated to aid GO adhesion and barrier properties. PEDOT:PSS integration is known to enhance mechanical integrity and conductivity of Al-GO/P surfaces while potentially adding barrier protection against corrosive species^166^.

Building upon this understanding, the antibiofouling, biocorrosion resistance and biocompatible nature of the synthesized Al_GO/P substrates were investigated. In order to evaluate the significance of PEDOT:PSS as an additive, Al_GO control surfaces were prepared without PEDOT:PSS. Broad-spectrum antimicrobial activity was assessed qualitatively and quantitatively via growth inhibition and cell viability assay, respectively. Results from antimicrobial assays demonstrated the superior antimicrobial properties of Al_GO100P, Al_GO250P, and Al_GO500P compared to Al_GO50P, Al_GO, and bare Al (Figure 5, Figures S5 and S7, Supporting Information).

Likewise, Al_GO/P substrates; Al_GO100P, Al_GO250P, and Al_GO500P displayed disrupted biofilm structures (Figures 6D, 6E, and 6F) and significantly lower biofilm biomass (Table S4, Supporting Information) relative to bare Al (Figure 6B and Table S4, Supporting Information). These findings were further substantiated through advanced microscopic (SEM) analysis confirming the biocidal nature of the Al_GO/P substrates except Al_GO50P (Figure 7C), which failed to show any contact-mediated membrane damage.

Next, the real-time performance of Al_GO/P were examined by evaluating biocorrosion resistance properties under abiotic and biotic immersion environments. Analyses from corrosion rates (Figure 9A), pH kinetics (Figure 9B), ICP-MS (Table S5, Supporting Information) and FESEM imaging (Figures S11 and S12, Supporting Information) revealed that Al_GO/P substrates exhibited superior biocorrosion resistance in the order; Al_GO500P, Al_GO250P, Al_GO100P, and Al_GO50P compared to controls (CR steel, bare Al, and Al_GO). Abiotic immersion environment in presence of the control substrates (CR steel, bare Al, and Al_GO) exhibited alkaline pH related to ionic leaching (Figure 9Bi), while Al_GO/P substrates maintained neutral pH (Figure 9Bi). Under biotic immersion conditions, Al_GO/P substrates also displayed neutral pH environment (Figure 9Bii), suggesting attenuated microbial colonization, and lack of production of microbial products, correlating to earlier research^148^. The aforementioned observations were further supported by ICP-MS analysis, which showed reduced Al^3+^ release during the 45 day immersion period for Al_GO/P substrates, relative to controls (CR steel, bare Al, and Al_GO). Rhyming with these results FESEM images also highlighted the superior corrosion resistance properties of fabricated Al_GO/P surfaces (Figures S17 and S19, Supporting Information).

Therefore, in the current study, PEDOT:PSS incorporation likely enhanced surface stability preventing ion leaching, significantly improving corrosion resistance of Al_GO/P surfaces. The improved corrosion resistance of Al_GO/P can be further correlated with reduced porosity percentages (Table 3), as shown by earlier literatures^57,116^. Current results suggest a reduction in porosity percentage with increase in GO concentration from 50 µg/ml to 500 µg/ml for the Al_GO/P test surfaces. Coatings with low porosity percentage has been shown to maximize barrier performance by restricting ion transport pathways^69,119^. There are also reports correlating porosity percentage to antimicrobial action; smooth surfaces with fewer porous domains resist microbial attachment and biofilm formation^116^. Therefore, Al_GO/P surfaces with lower porosity percentage should improve corrosion resistance, by decreasing microbial contamination and lower MIC (Figure 8, Figures S17 and S19, Supporting Information).

Next, it was crucial to conduct biocompatibility testing to ensure the safety and suitability of Al_GO/P as implant materials. Al_GO100P was identified as a promising candidate for *in vivo* implant material, owing to its excellent biocompatibility (Figure 10) along with remarkable antibiofilm (Figure 6) and anticorrosion properties (Figure 8, Figures S17 and S19, Supporting Information). Al_GO250P demonstrated favorable biocompatibility profile at 24 hours and minimal cytotoxicity at 48 hours, suggesting potential suitability for short-term biomedical applications. In contrast, the reduced HEK293T cell viability observed with Al_GO500P and Al_GO1000P suggests a potential for cytotoxicity, making them better suited for *ex vivo* applications. Controls; bare Al and Al_GO surfaces displayed remarkable cytotoxic activity (Figure 10, Figure S21, Supporting Information). The toxic effects of aluminum (Al^3+^) result from its ability to disrupt cellular homeostasis through multiple mechanisms, including oxidative stress induction, enzymatic inhibition, and direct cellular membrane damage^158^. The results from both indirect and direct contact assays indicated that GO-PEDOT:PSS hybrid coating including GO at 100 µg/ml and 250 µg/ml concentration, may effectively reduce Al-related cytotoxicity while preserving biocompatibility, essential for biomedical applications. Earlier works have employed GO coatings at higher concentrations (0.5-10 mg/ml) to achieve enhanced antimicrobial activity^49–53^. Further GO at a concentration of 1 mg/ml on AA1050 has shown protection against corrosive agents (water, oxygen, ions), while displaying systemic toxicity due to inflammatory responses. Current work has found that Al_GO/P substrates including lower concentration of GO (100 and 250 µg/ml) have excellent antimicrobial efficacy and biocompatibility for use as implant materials.

As reported earlier, the membrane-damaging activity of GO involves multiple mechanisms, such as GO sharp-edge effects, membrane pore formation, phospholipid extraction, and electron extraction by GO side groups, and ROS production which induce oxidative stress^44,63^. To elucidate the underlying mechanism of Al_GO/P substrates and correlate cell membrane damage with potential electron extraction events, intracellular and surface ROS production assays were performed using H_2_DCFDA and DHE fluorescent probes (Figure 8). The presence of intracellular ROS indicated microbial cell death, whereas the detection of ROS on the Al_GO/P surface pointed towards an interface phenomenon that suggested electron transfer directed ROS generation over the conductive Al_GO/P substrates.

Summarizing the experimental data, a mechanistic model underlying the robust antimicrobial (Figure 5) and antibiofilm activity (Figure 6) of the fabricated Al_GO/P substrates (Al_GO100P, Al_GO250P, and Al_GO500P) is discussed. First, surface hydrophobicity (Table 2) of the Al_GO/P substrates promote stable interactions with microbial cells, facilitating contact-mediated membrane damage on GO-rich surface. Moreover, the conductive nature (Table 2) of the Al_GO/P substrates may have induced electron transfer events that generate ROS (Figure 8), further contributing to microbial cell damage (Figure 7). Further, the Al_GO/P surfaces were evaluated to possess low porosity percentages (Table 3), subsequently limiting microbial colonization and superior antibiofilm activity^116^. Prior work showed low porosity levels correlate with improve corrosion resistance, by decreasing microbial contamination. Together, these physical and electrochemical interactions may have enhanced the antibiofouling efficacy of fabricated Al_GO/P surfaces. The superior antibiofouling activities observed for test substrates; Al_GO500P, Al_GO250P, Al_GO100P relative to Al_GO50P can be linked to higher surface hydrophobicity, low porosity percentage, and higher ROS production (Figure 8, Table 2, Table 3) relative to bare Al.

Beyond their antibiofouling, biocorrosion resistant, and biocompatible nature; the ecological implications of the synthesized Al_GO/P substrates warrant careful consideration to ensure safe deployment in biomedical or environmental settings. Although fabricated Al_GO/P surfaces exhibited improved GO adhesion in presence of PEDOT:PSS, their long-term environmental fate and potential release of micro-debris remain unaddressed. This underscores the need for future lifecycle and ecotoxicity studies on GO and PEDOT:PSS leaching from metal surfaces^167,168^. However, the coated Al surfaces can be further recycled following eco-friendly removal and disposal of the GO-PEDOT:PSS coating.

In toto, the findings from this study may establish a strong foundation for practical implementation of GO-PEDOT:PSS hybrid coated Al substrates. The synthesized substrates may provide a promising approach for improving the durability and performance of Al; particularly where corrosion resistance, antibiofouling nature, and biocompatibility are crucial, such as in the health-care, water treatment, and varied industrial sectors, where MIC plays detrimental roles. Further research and optimization of the Al_GO/P composite system could facilitate the development of advanced Al-based materials, particularly in the design of next-generation biomedical implants with improved material lifespan.

## Author Contributions

**S.S.N.**, Conceptualization; Methodology; Validation; Formal analysis; Investigation; Resources; Data curation; Writing original draft; Writing review and editing; Visualization. **R.E.A.**, Validation; Formal analysis; Writing original draft; Writing review and editing. **A.K.R.**, Validation; Investigation; Writing review and editing; **T.N.N.**, Validation; Formal analysis; Resources; Writing original draft; Writing review and editing. **S.N.**, Conceptualization; Methodology; Validation; Formal analysis; Investigation; Resources; Writing original draft; Writing review and editing; Supervision; Project administration; Funding acquisition.

## Acknowledgements

Aluminum substrates were provided by NALCO India, Ltd. The authors gratefully acknowledge the School of Basic Sciences at IIT Bhubaneswar for granting access to their facilities for substrate synthesis and porosity evaluation. Sincere thanks are extended to the central facilities at TIFR Hyderabad for their valuable support in the characterization of Al_GO/P substrates. The authors also appreciate the Central Research Facility at KIIT-DU, Bhubaneswar, for providing access to the scanning electron microscope (SEM). The infrastructure grant from the Department of Biotechnology, Government of India, under the Boost to University Interdisciplinary Life Science Departments for Education and Research (BUILDER) program at KIIT School of Biotechnology is also duly acknowledged. S.S.N. was supported by DD Innovation Incorporated, U.S.A. (Ref No: DD/KIIT/001/2023).

## Data Availability Statement

All data sources are cited within this article and its Supporting Information.

