## Supplementary Files for "Enhancing the biocorrosion resistance and biocompatibility of Aluminium substrates using Graphene Oxide-PEDOT:PSS Hybrid Coating"

### *Table of Contents*

*1. Determination of $I_D/I_G$ ratios from Raman Spectroscopy Data* ........ *S5*

*Table S1. Raman shift, peak intensities, FWHM, and $I_D/I_G$ ratio of D and G bands for pristine GO, Al_GO, and Al_GO/P substrates.* ........ *S5*

*2. Raman Spectra of pristine PEDOT:PSS* ........ *S5*

*Figure S1. The Raman spectrum of PEDOT:PSS (Sigma-Aldrich, Product No. 655201).* ........ *S6*

*3. Chemical state, functional group retention after deposition confirmed by X-ray photoelectron spectroscopy (XPS)* ........ *S6*

*Figure S2. XPS analysis of Al_GO and Al_GO/P substrates.* ........ *S6*

*Table S2. Atomic weight (Atomic wt%) percentages of pristine GO, Al_GO, and Al_GO/P substrates.* *S7*

*4. Water contact angle measurements* ........ *S7*

*Figure S3. Static water contact angle measurement.* ........ *S7*

*5. Porosity evaluation* ........ *S8*

*Figure S4. Electrochemical impedance spectroscopy (EIS) analysis of bare Al vs Al_GO/P substrates in 3.5% NaCl solution, presented as Nyquist plots.* ........ *S10*

*6. Growth inhibition assay: synthesized Al_GO substrates, pristine GO, & PEDOT:PSS* ........ *S10*

*Figure S5. Growth Inhibition Assay of Al_GO substrates.* ........ *S10*

*Growth non-inhibitory activity of pristine GO and PEDOT:PSS:* ........ *S11*

*Figure S6. (A) Growth Inhibition Assay of E. coli, S. aureus, and C. albicans in presence of GO in solution.* ........ *S11*

*7. Cell viability assay of E. coli, S. aureus, and C. albicans in presence of control and test substrates for a period of 10 hours* ........ *S12*

*Figure S7. Cell viability assay of (A) E. coli, (B) S. aureus, and (C) C. albicans was performed to assess the survival of cells on various control and test surfaces.* ........ *S12*

*8. Cell viability assay using corresponding antibiotics against relevant pathogens* ........ *S13*

*Figure S8. Cell viability assay of E. coli, S. aureus, and C. albicans against different antibiotics* ........ *S13*

*9. Fluorescence microscopic images of biofilm formation assay using mixed Candida-bacterial species at $10^8$ CFU* ........ *S14*

*Figure S9. Fluorescence microscopic images of mixed species (C. albicans, S. aureus, and E. coli) biofilm ($10^8$ CFU). (A, B) Mixed Candida-bacterial suspensions were exposed to control surfaces (Glass and Bare Al), and (C–F) to various Al_GO/P substrates.* ........ *S14*

*10. Fluorescence quantification of mixed biofilm images using ImageJ software* ........ *S15*

*Figure S10. Graphical representation of the mean fluorescence intensity ratios* ........ *S15*

*11. Biofilm biomass quantification of mixed Candida-bacterial species ($10^7$ CFU) after exposure to control and Al_GO/P surfaces* ........ *S16*

*Figure S11. Quantification of mixed species ($10^7$ CFU) biofilm biomass exposed to control (glass and bare Al) and test substrates (Al_GO/P) after crystal violet (CV) staining.* ........ *S16*

*12. Biofilm biomass quantification of mixed Candida-bacterial species ($10^8$ CFU) after exposure to control and Al_GO/P surfaces* ........ *S17*

*Figure S12. Quantification of mixed species ($10^8$ CFU) biofilm biomass exposed to control (glass, bare Al) and test substrates (Al_GO/P) after crystal violet (CV) staining.* ........ *S17*

*13. Biofilm biomass quantification of mixed Candida-bacterial species ($10^7$ CFU) in presence of pristine GO solution* ........ *S18*

*Figure S13. Quantification of mixed species ($10^7$ CFU) biofilm biomass in presence of pristine GO after crystal violet (CV) staining* ........ *S18*

*14. Pairwise comparison of antibiofilm efficiency between pristine GO solution vs Al_GO/P surfaces at $10^7$ CFU inoculum load* ........ *S19*

*Figure S14. Pairwise comparison of mixed species ($10^7$ CFU) biofilm biomass exposed to pristine GO solution vs. Al_GO/P substrates at corresponding concentrations after crystal violet (CV) staining.* . *S19*

*15. Quantitative percentages of biofilm formation* ........ *S20*

*Table S4. Quantitative values of percentage biofilm formation of mixed Candida-bacterial species at two distinct inoculum ($10^7$ and $10^8$ CFU) concentrations exposed to Al_GO/P substrates and pristine GO solution.* ........ *S20*

*16. Quantitative estimation of mean fluorescence intensity per cell corresponding to fluorescent images of ROS production using ImageJ software* ........ *S21*

*Figure S15. Graphical representation of the average fluorescence intensity per cell* ........ *S21*

*17. ICP-MS analysis* ........ *S22*

*Table S5. ICP-MS analysis to determine metal ion leaching in abiotic and biotic immersion environment.* ........ *S22*

*18. FESEM images of Al_GO and Al_GO/P substrates before and after corrosion under abiotic environment* ........ *S23*

*Figure S16. FESEM analysis of control (bare Al, CR Steel, Al_GO50, Al_GO100, Al_GO250, and Al_GO500) substrates under abiotic immersion conditions.* ........ *S23*

*Figure S17. FESEM analysis of control (bare Al and CR Steel) and test (Al_GO/P) substrates under abiotic immersion conditions.* ........ *S24*

*19. FESEM images of Al_GO and Al_GO/P substrates before and after corrosion under biotic environment* ........ *S25*

*Figure S18. FESEM analysis of control (bare Al, CR Steel, Al_GO50, Al_GO100, Al_GO250, and Al_GO500) substrates under biotic immersion conditions.* ........ *S25*

*Figure S19. FESEM analysis of control (bare Al and CR Steel) and test (Al_GO/P) substrates under biotic immersion conditions.......... S26*

*20. Optical images of substrates before and after corrosion under abiotic and biotic environments .. S27*

*Figure S20. Optical images of controls (bare Al, CR Steel, Al_GO substrates) and test Al_GO/P substrates .......... S27*

*21. Cytotoxicity Assay: quantitative assessment of cell viability percentages .......... S28*

*Table S6A. HEK293T cells were exposed to 100% crude extracts of different Al_GO/P substrates for a period of 48 hours and MTT reduction was evaluated. Results are expressed as % of the control cells (HEK293T, 100%) not exposed to crude extracts. Values are the average of three independent biological replicates, expressed as mean ± SD.......... S28*

*Table S6B. HEK293T cells were exposed directly to the synthesized Al_GO/P substrates for a period of 48 hours and MTT reduction was evaluated. Results are expressed as % of the control cells (HEK293T, 100%) not exposed to Al_GO/P substrates. Values are the average of three independent biological replicates, expressed as mean ± SD.......... S29*

*22. Cytotoxicity Assay: quantitative assessment of cell viability percentages of HEK293T cells in presence of Al_GO substrates .......... S30*

*23. Cytotoxicity Assay: quantitative assessment of cell viability percentages of HEK293T cells in presence of pristine GO.......... S31*

*Figure S22. Effect of pristine GO on cell viability of HEK293T cells.......... S31*

*24. Cytotoxicity Assay: quantitative assessment of cell viability percentages of HEK293T cells in presence of PEDOT:PSS.......... S32*

*Figure S23. Effect of PEDOT:PSS on cell viability of HEK293T cells.......... S32*

*Reference.......... S33*

***1. Determination of $I_D/I_G$ ratios from Raman Spectroscopy Data***

***Table S1. Raman shift, peak intensities, FWHM, and $I_D/I_G$ ratio of D and G bands for pristine GO, Al_GO, and Al_GO/P substrates.***

| *Sample* | *Raman shift ($cm^{-1}$)* | | *Intensities* | | *FWHM* | | *$I_D/I_G$* |
|---|---|---|---|---|---|---|---|
| | *D band* | *G band* | *D band* | *G band* | *D band* | *G band* | |
| ***Pristine GO*** | 1345.58 | 1588.47 | 0.91 | 1.00 | 137.14 | 95.3 | 0.91 |
| ***Al_GO50*** | 1342.68 | 1602.07 | 0.93 | 1.00 | 127.07 | 82 | 0.93 |
| ***Al_GO100*** | 1342.68 | 1600.98 | 0.97 | 1.00 | 132.71 | 87.49 | 0.97 |
| ***Al_GO250*** | 1341.54 | 1603.16 | 1.00 | 0.93 | 119.2 | 84.21 | 1.07 |
| ***Al_GO500*** | 1339.27 | 1603.16 | 0.94 | 1.00 | 116.91 | 75.42 | 0.94 |
| ***Al_GO50P*** | 1343.31 | 1601.58 | 0.91 | 1.00 | 116.9 | 76.52 | 0.91 |
| ***Al_GO100P*** | 1342.17 | 1603.77 | 0.88 | 1.00 | 114.65 | 71.01 | 0.88 |
| ***Al_GO250P*** | 1333.08 | 1596.12 | 0.86 | 1.00 | 115.91 | 68.86 | 0.86 |
| ***Al_GO500P*** | 1339.90 | 1601.58 | 0.86 | 1.00 | 106.78 | 66.65 | 0.86 |

***2. Raman Spectra of pristine PEDOT:PSS***

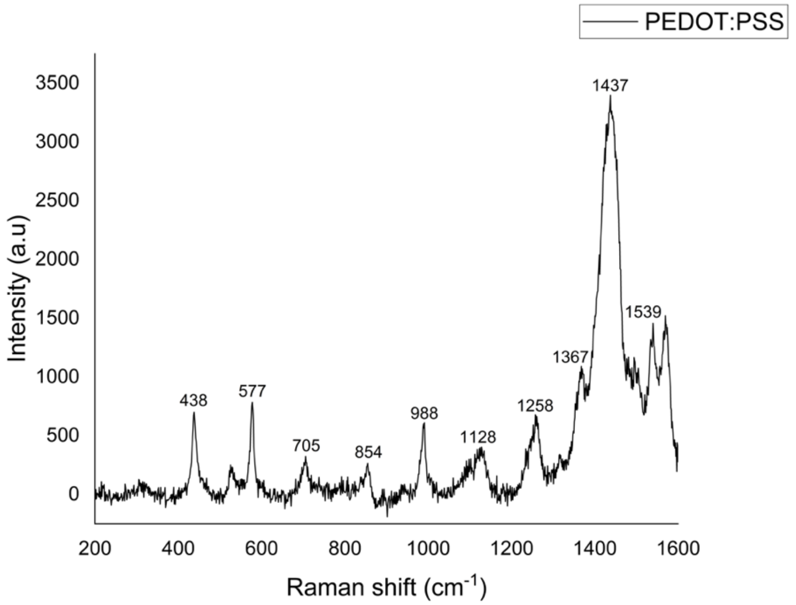

***Figure S1. The Raman spectrum of PEDOT:PSS (Sigma-Aldrich, Product No. 655201).*** *Raman spectroscopy experimentally confirms PEDOT:PSS identity, aligning with structural characteristics reported in prior studies[1].*

#### *3. Chemical state, functional group retention after deposition confirmed by X-ray photoelectron spectroscopy (XPS)*

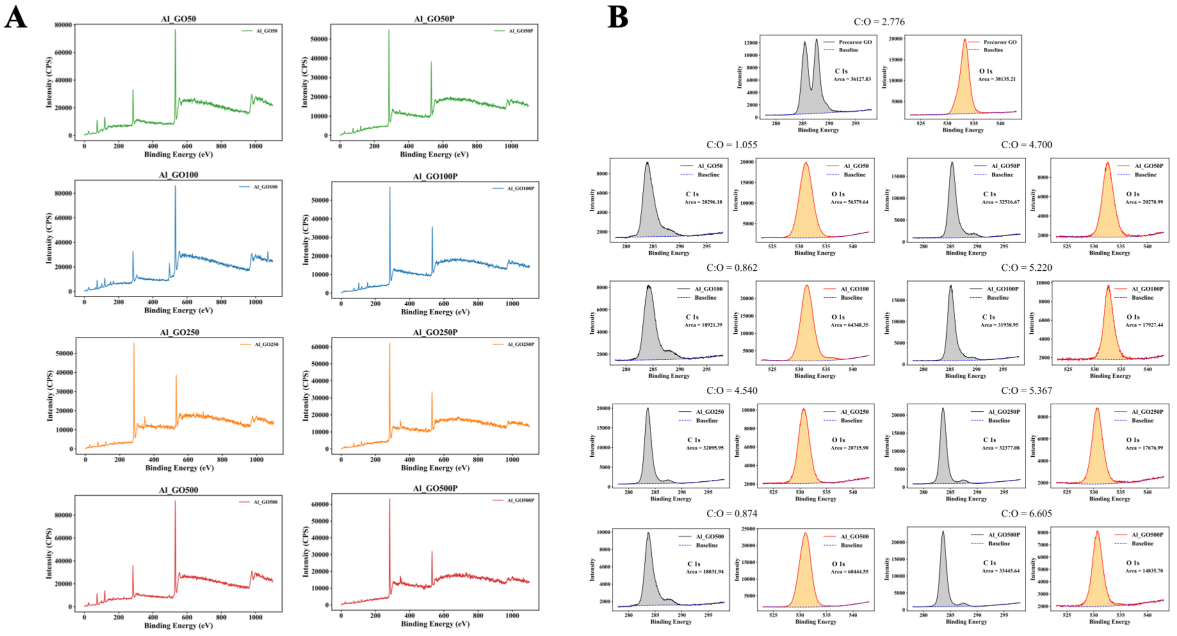

***Figure S2. XPS analysis of Al_GO and Al_GO/P substrates.*** *(A) Survey spectra showing the overall elemental composition of the coated substrates confirming the presence of characteristic $C_{1s}$ and $O_{1s}$ signals associated with GO deposition and PEDOT:PSS modification. (B) High-resolution $C_{1s}$ and $O_{1s}$ spectra for all samples, used to determine carbon and oxygen peak areas and calculate C:O ratios, revealing systematic changes in chemistry with increasing GO concentration and incorporation of PEDOT:PSS.*

***Table S2. Atomic weight (Atomic wt%) percentages of pristine GO, Al_GO, and Al_GO/P substrates.*** *The area under curves is used to calculate the weight percentage of the atomic compositions of pristine GO, GO-only, and GO and PEDOT:PSS coatings on various substrates.*

| ***Elements*** | ***Atomic weight (%)*** | | | | | | | | |
|---|---|---|---|---|---|---|---|---|---|
| | ***GO*** | ***Al_GO50*** | ***Al_GO100*** | ***Al_GO250*** | ***Al_GO500*** | ***Al_GO50P*** | ***Al_GO100P*** | ***Al_GO250P*** | ***Al_GO500P*** |
| ***Carbon*** | 73.5 | 51.3 | 46.3 | 81.9 | 46.6 | 82.5 | 83.9 | 84.3 | 86.9 |
| ***Oxygen*** | 26.5 | 48.7 | 53.7 | 18.1 | 53.4 | 17.5 | 16.1 | 15.7 | 13.1 |
| ***Carbon:Oxygen*** | 2.8 | 1.1 | 0.9 | 4.5 | 0.9 | 4.7 | 5.2 | 5.4 | 6.6 |

### *4. Water contact angle measurements*

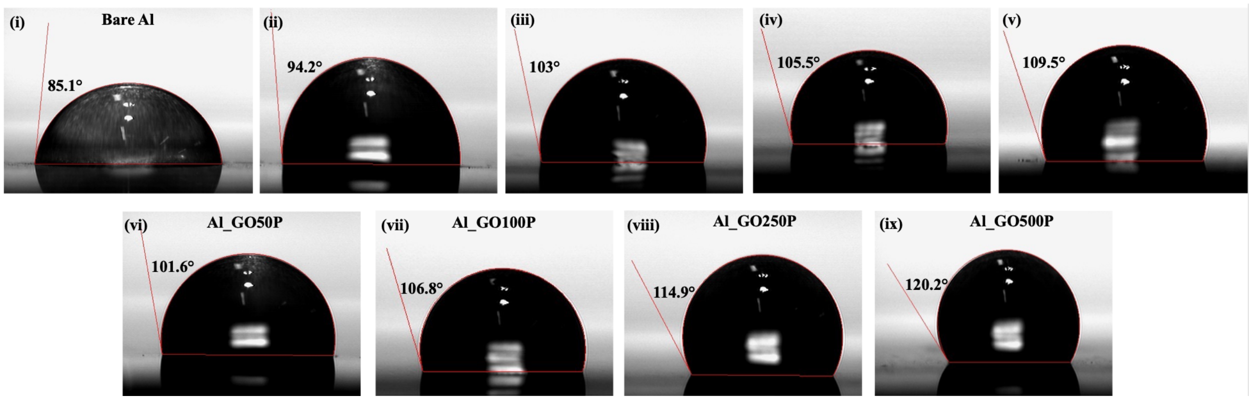

***Figure S3. Static water contact angle measurement.*** *The static water contact angle (WCA) measurements of (i) Bare Al, Al_GO substrates; (ii) Al_GO50, (iii) Al_GO100, (iv) Al_GO250, and (v) Al_GO500, and Al_GO/P substrates; (vi) Al_GO50P, (vii) Al_GO100P, (viii) Al_GO250P, and (ix) Al_GO500P via sessile drop method.*

### *5. Porosity evaluation*

***Table S3. Tafel fit and $R_p$ fit parameters.***

| *Substrates* | *$R_p$ Fit* | *Tafel fit* |
|---|---|---|
| ***Bare Al*** | parameters:<br>beta c = 120.0 mV<br>beta a = 120.0 mV<br>range = +/- 25.0 mV<br>results:<br>Rp = 145 Ohm<br>Ecorr = -471.38 mV vs. Ref<br>correlation = 0.991<br>Icorr = 179.531 µA | results:<br>(x) Ecorr = -470.180 mV<br>(x) Icorr = 37.842 uA<br>(x) beta c = 37.6 mV<br>(x) beta a = 32.4 mV<br>$Chi^2$ = 0.718 931<br>Chi / sqrt(N) = 0.149 889<br>equivalent weight = 9.000 g/eq.<br>density = 2.700 g/$cm^3$<br>surface area = 1.000 $cm^2$<br>corrosion rate = 0.412 73 mmpy |
| ***Al_GO50P*** | parameters:<br>beta c = 120.0 mV<br>beta a = 120.0 mV<br>range = +/- 25.0 mV<br>results:<br>Rp = 163 Ohm<br>Ecorr = -485.025 mV vs. Ref<br>correlation = 0.997 5<br>Icorr = 160.259 µA | results:<br>(x) Ecorr = -485.346 mV<br>(x) Icorr = 116.871 uA<br>(x) beta c = 87.7 mV<br>(x) beta a = 95.9 mV<br>$Chi^2$ = 0.529 288<br>Chi / sqrt(N) = 0.128 609<br>equivalent weight = 9.000 g/eq.<br>density = 2.700 g/$cm^3$<br>surface area = 1.000 $cm^2$<br>corrosion rate = 1.274 67 mmpy |
| ***Al_GO100P*** | parameters:<br>beta c = 120.0 mV<br>beta a = 120.0 mV<br>range = +/- 25.0 mV | results:<br>(x) Ecorr = -491.119 mV<br>(x) Icorr = 233.778 uA<br>(x) beta c = 731 970 304.0 mV |

| | | |
|---|---|---|
| | results:<br>Rp = 176 Ohm<br>Ecorr = -493.717 mV vs. Ref<br>correlation = 0.982 9<br>Icorr = 148.523 µA | (x) beta a = 104.1 mV<br>Chi² = 0.767 443<br>Chi / sqrt(N) = 0.152 499<br>equivalent weight = 9.000 g/eq.<br>density = 2.700 g/cm³<br>surface area = 1.000 cm²<br>corrosion rate = 2.549 74 mmpy |
| ***Al_GO250P*** | parameters:<br>beta c = 120.0 mV<br>beta a = 120.0 mV<br>range = +/- 25.0 mV<br>results:<br>Rp = 222 Ohm<br>Ecorr = -489.79 mV vs. Ref<br>correlation = 0.993 7<br>Icorr = 117.552 µA | results:<br>(x) Ecorr = -488.030 mV<br>(x) Icorr = 364.680 uA<br>(x) beta c = 2 041 958 528.0 mV<br>(x) beta a = 199.3 mV<br>Chi² = 1.016 03<br>Chi / sqrt(N) = 0.097 445 5<br>equivalent weight = 9.000 g/eq.<br>density = 2.700 g/cm³<br>surface area = 1.000 cm²<br>corrosion rate = 3.977 44 mmpy |
| ***Al_GO500P*** | parameters:<br>beta c = 120.0 mV<br>beta a = 120.0 mV<br>range = +/- 25.0 mV<br>results:<br>Rp = 454 Ohm<br>Ecorr = -477.145 mV vs. Ref<br>correlation = 0.997 4<br>Icorr = 57.448 3 µA | results:<br>(x) Ecorr = -477.071 mV<br>(x) Icorr = 23.883 uA<br>(x) beta c = 57.0 mV<br>(x) beta a = 55.8 mV<br>Chi² = 0.931 415<br>Chi / sqrt(N) = 0.170 607<br>equivalent weight = 9.000 g/eq.<br>density = 2.700 g/cm³<br>surface area = 1.000 cm²<br>corrosion rate = 0.260 484 mmpy |

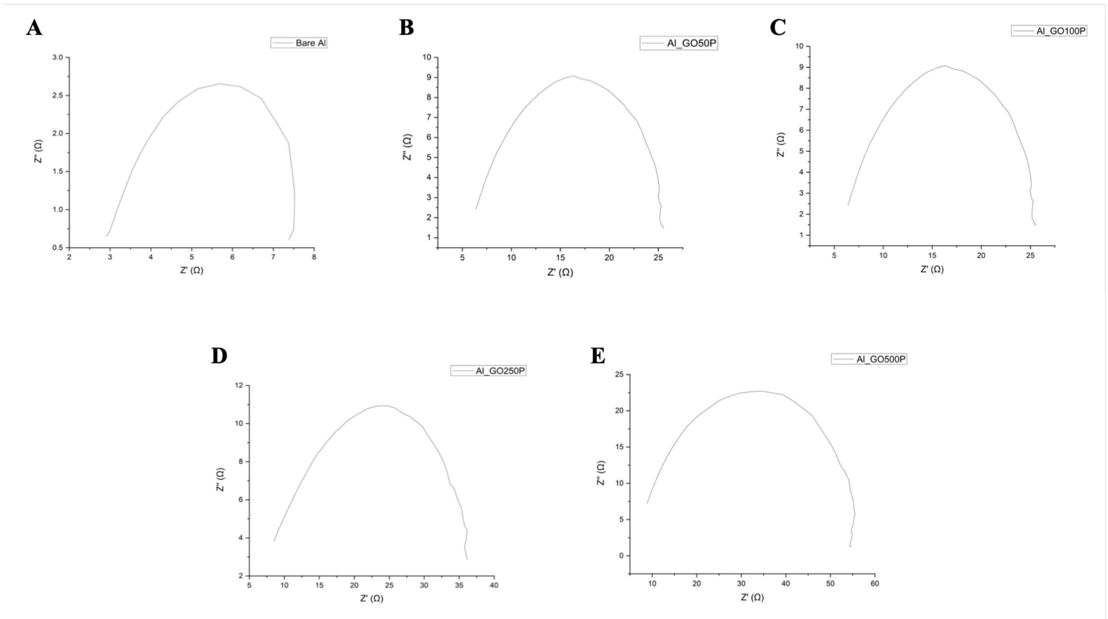

***Figure S4. Electrochemical impedance spectroscopy (EIS) analysis of bare Al vs Al_GO/P substrates in 3.5% NaCl solution, presented as Nyquist plots.***

### *6. Growth inhibition assay: synthesized Al_GO substrates, pristine GO, & PEDOT:PSS*

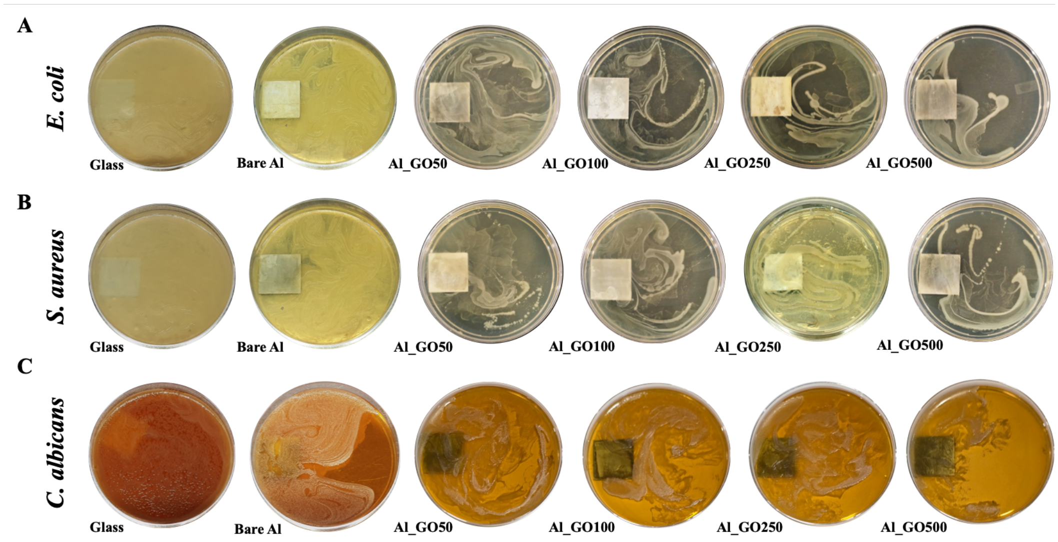

***Figure S5. Growth Inhibition Assay of Al_GO substrates.*** *After 10 hour of incubation on different control (glass and bare Al) and Al_GO substrates, (**A, B**) bacterial (E. coli, S. aureus) and (**C**) fungal (C. albicans) cells were cultured on LB and YPD agar plates, respectively. The inhibitory*

*roles of the coated substrates were assessed by examining colony growth the following day. Cells cultivated on glass and bare Al surfaces showed dense bacterial and fungal colony formation, confirming absence of antimicrobial potential. Likewise, the Al_GO surfaces exhibited no antimicrobial efficacy, as evidenced by the abundant colony growth comparable to the controls.*

Growth non-inhibitory activity of pristine GO and PEDOT:PSS:

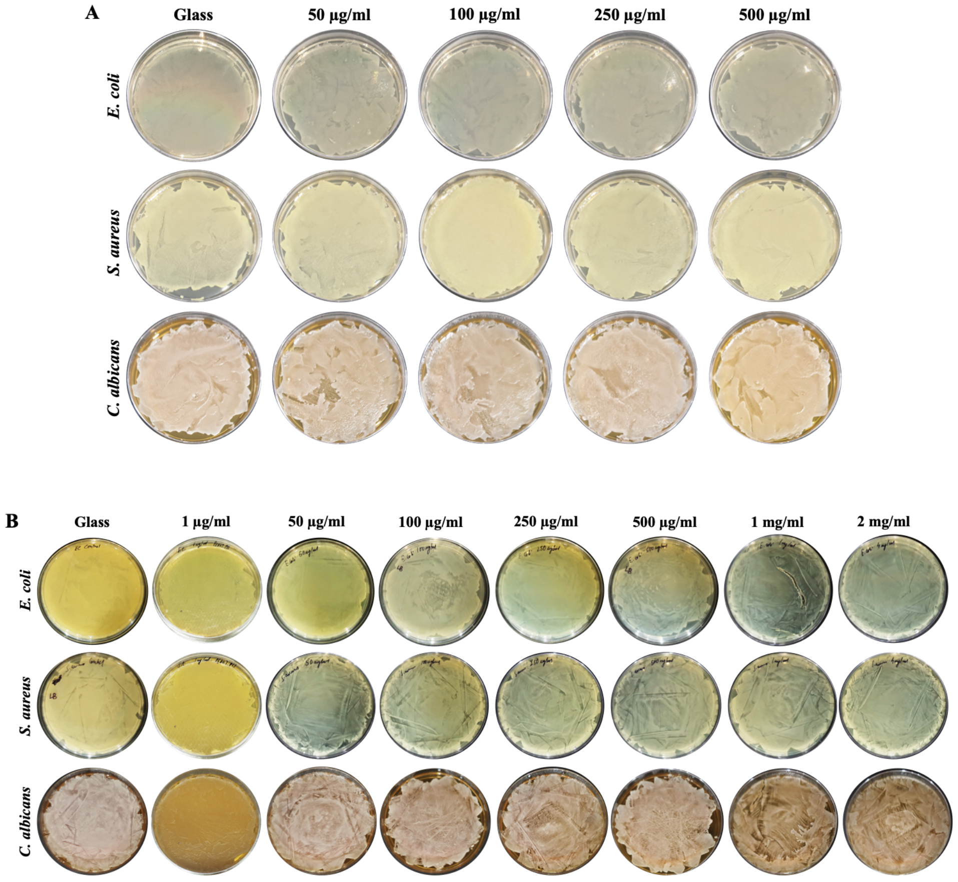

***Figure S6. (A) Growth Inhibition Assay of E. coli, S. aureus, and C. albicans in presence of GO in solution.*** *Cells cultivated on glass surfaces exhibited significant bacterial and fungal colony formation, suggesting minimal antimicrobial properties. Exposure to varying concentrations of GO (50 µg/ml, 100 µg/ml, 250 µg/ml, 500 µg/ml) resulted in abundant colony growth.* ***(B) Growth Inhibition Assay of E. coli, S. aureus, and C. albicans in presence of pristine PEDOT:PSS.*** *Cells cultivated on glass surfaces exhibited significant bacterial and fungal colony formation, suggesting minimal antimicrobial properties. Exposure to varying concentrations of PEDOT:PSS (1 µg/ml, 50 µg/ml, 100 µg/ml, 250 µg/ml, 500 µg/ml, 1 mg/ml, and 2 mg/ml) resulted in abundant colony growth.*

*7. Cell viability assay of E. coli, S. aureus, and C. albicans in presence of control and test substrates for a period of 10 hours*

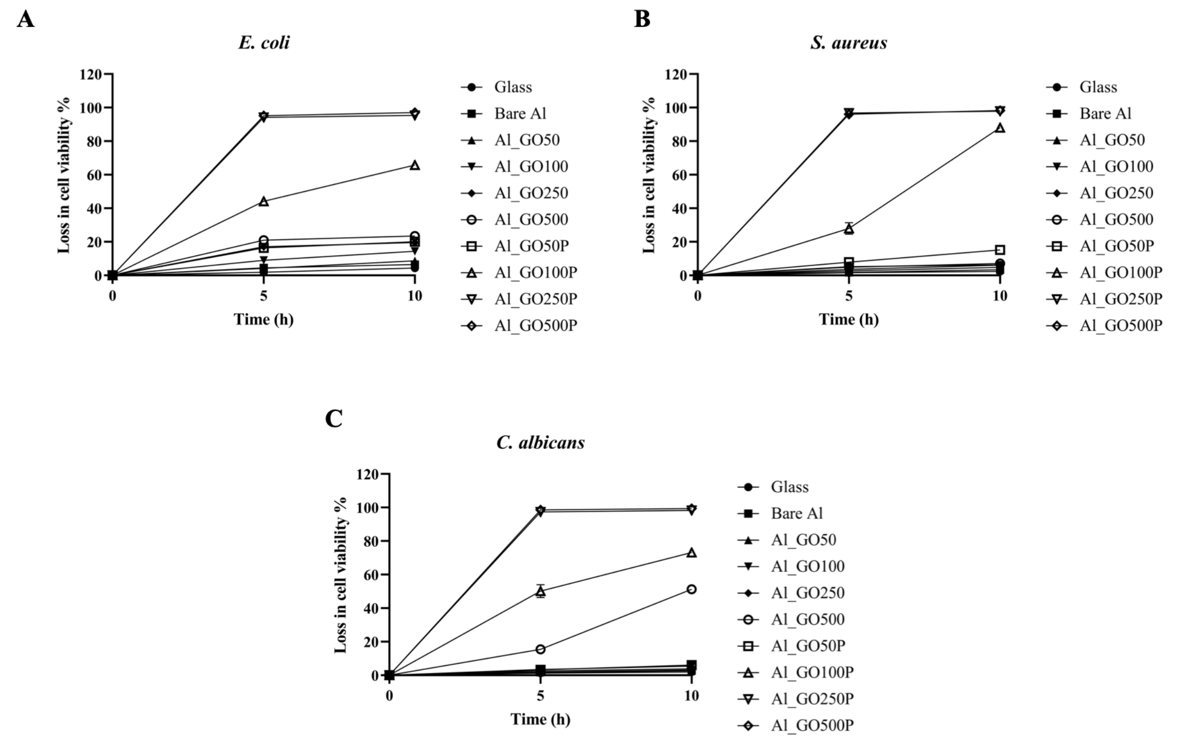

***Figure S7. Cell viability assay of (A) E. coli, (B) S. aureus, and (C) C. albicans was performed to assess the survival of cells on various control and test surfaces.*** *Following incubation for different time intervals (0, 5, and 10 h), samples were collected and recultivated on LB (for bacteria) and YPD (for yeast) agar plates. The reduction in cell viability was quantified by the colony counting method. Error bars indicate ±1 standard deviation (SD) from the mean of three independent experiments (n = 3).*

***8. Cell viability assay using corresponding antibiotics against relevant pathogens***

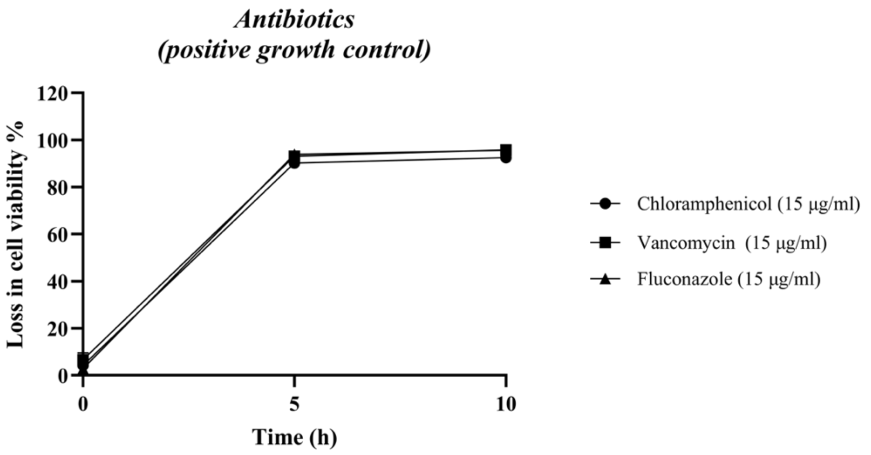

***Figure S8. Cell viability assay of E. coli, S. aureus, and C. albicans against different antibiotics***; Chloramphenicol (15 µg/ml), Vancomycin (15 µg/ml), and Fluconazole (15 µg/ml), respectively at 0, 5, and 10 h. Post-incubation at different time-points, $OD_{600}$ values were measured using UV-Vis spectrophotometer. Loss in cell viability was calculated using standard formula. Error bars represent ±1 standard deviation (SD) from mean for three independent biological replicates (n=3).

***Cell viability assay-*** Antibacterial; chloramphenicol (15 µg/ml) and vancomycin (15 µg/ml) was used against *E. coli* and *S. aureus*, respectively. Antifungal; fluconazole (15 µg/ml) was used against *C. albicans* cells. The cells were grown in presence of relevant antibiotics and 1 ml of sample was collected at 0, 5, and 10 h for $OD_{600}$ measurement using UV-Vis spectrophotometer (Agilent, Cary 60). The loss of cell viability was calculated using formula:

$$Loss\ of\ cell\ viability\ (\%) = 100 - \left[\frac{OD_{600,test}}{OD_{600,untreated}} x\ 100\right]$$

The results were plotted using GraphPad Prism version 9.0 for Windows, GraphPad Software, San Diego, CA, U.S.A. (www.graphpad.com).

***9. Fluorescence microscopic images of biofilm formation assay using mixed Candida-bacterial species at $10^8$ CFU***

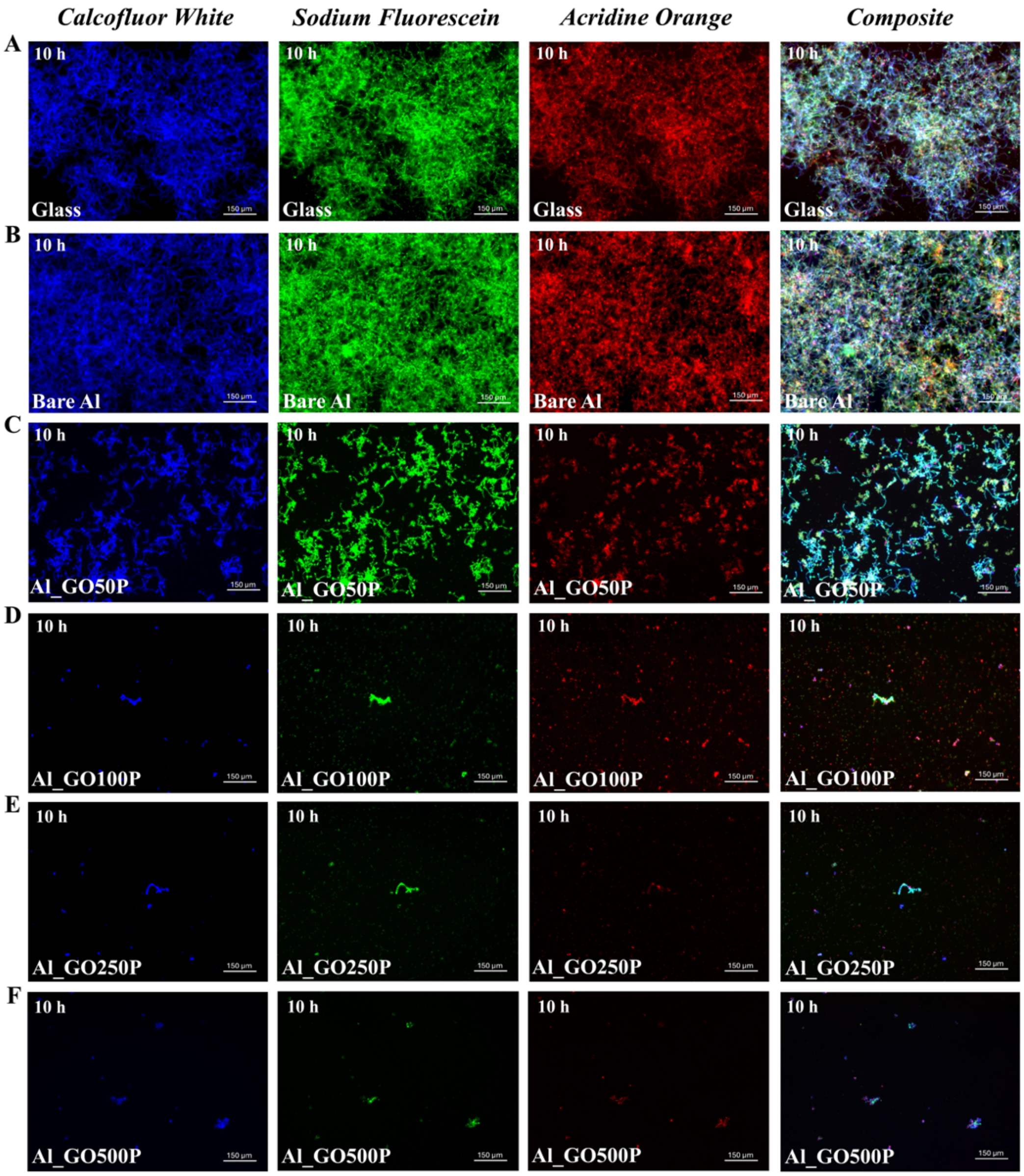

***Figure S9. Fluorescence microscopic images of mixed species (C. albicans, S. aureus, and E. coli) biofilm ($10^8$ CFU). (A, B) Mixed Candida-bacterial suspensions were exposed to control surfaces (Glass and Bare Al), and (C–F) to various Al_GO/P substrates.*** *Mixed biofilms were allowed to grow for 10 hours under biofilm stimulating conditions after which the samples were stained with calcofluor white (blue), acridine orange (red), and sodium fluorescein (green) to visualize C. albicans, S. aureus, and E. coli, respectively. Biofilms were imaged using an in-house fluorescence microscope at 40× magnification.*

#### *10. Fluorescence quantification of mixed biofilm images using ImageJ software*

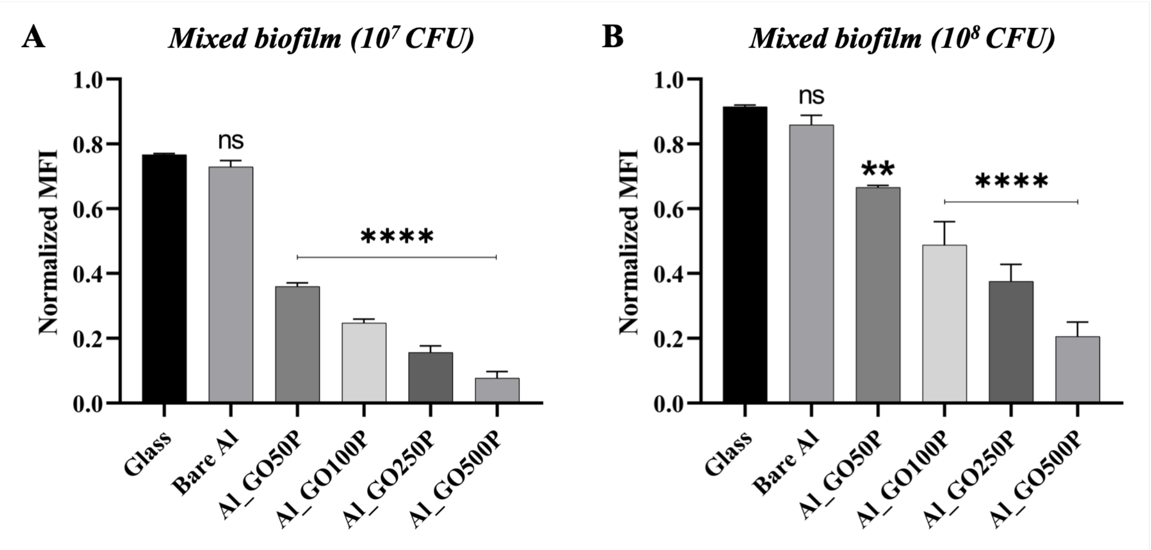

***Figure S10. Graphical representation of the mean fluorescence intensity ratios*** *(red/green) corresponding to fluorescence images of mixed-species biofilm formation at* ***(A)*** *$10^7$ CFU and* ***(B)*** *$10^8$ CFU microbial concentrations. Error bars represent ±1 standard deviation (SD) from the mean of fluorescence quantifications performed using ImageJ software.*

***11. Biofilm biomass quantification of mixed Candida-bacterial species ($10^7$ CFU) after exposure to control and Al_GO/P surfaces***

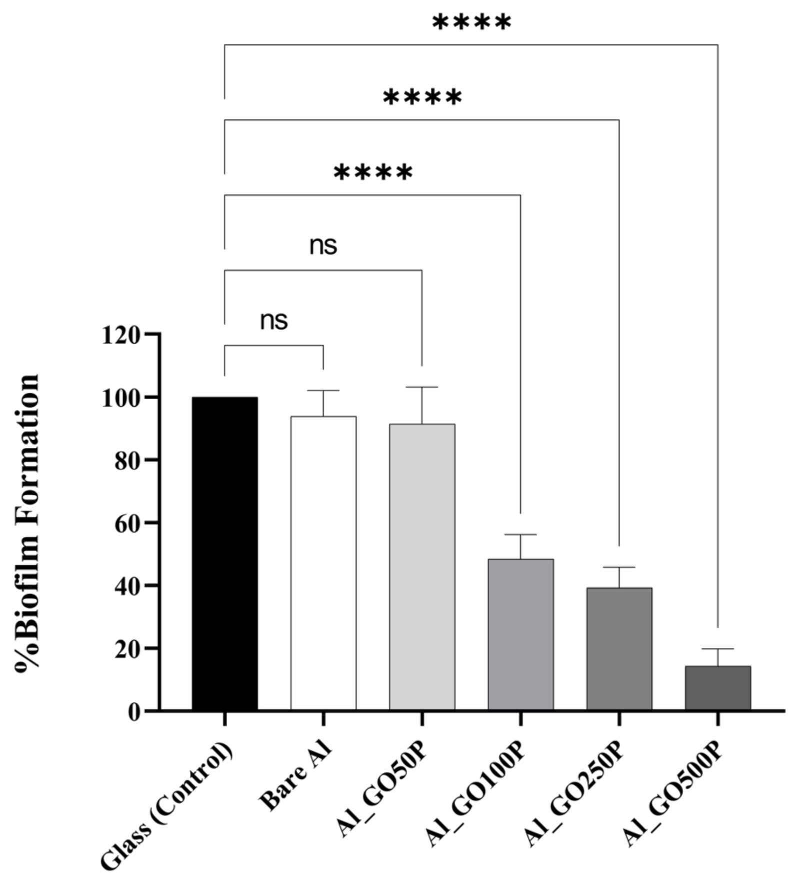

***Figure S11. Quantification of mixed species ($10^7$ CFU) biofilm biomass exposed to control (glass and bare Al) and test substrates (Al_GO/P) after crystal violet (CV) staining.*** *Inhibition of biofilm formation was assessed by incubating the mixed microbial cell suspension on glass, bare Al and Al_GO/P substrates (150 µl/cm$^2$) in sterile petri plates at room temperature under constant humidity for 10 hours. Post exposure, the cells were allowed to grow for 10 hours under biofilm stimulating conditions. Growth medium was discarded, and the adhered biomass was stained using CV. Percentage of biofilm formation is reported relative to glass (defined as 100%) and sterility control wells (defined as 0%). Error bars represent ±1 standard deviation (SD) from mean for three independent biological replicates (n=3). Statistical significance was assessed using one-way ANOVA; specific p-values of <0.05, <0.01, <0.001, and <0.0001 were indicated as *,**,***, and ****, respectively.*

***12. Biofilm biomass quantification of mixed Candida-bacterial species ($10^8$ CFU) after exposure to control and Al_GO/P surfaces***

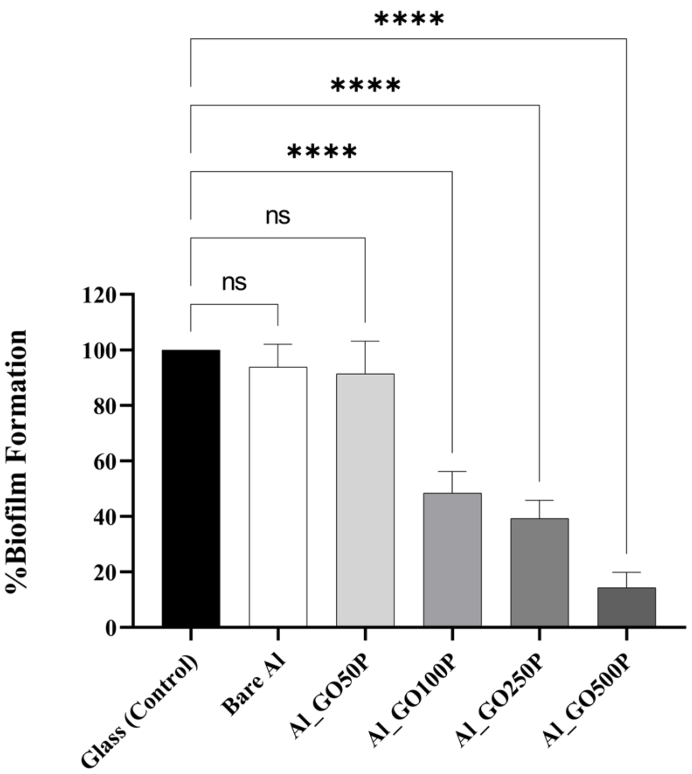

***Figure S12. Quantification of mixed species ($10^8$ CFU) biofilm biomass exposed to control (glass, bare Al) and test substrates (Al_GO/P) after crystal violet (CV) staining.*** *Inhibition of biofilm formation was assessed by incubating the mixed microbial suspension on glass, bare Al and Al_GO/P substrates (150 µl/cm$^2$) in sterile petri plates at room temperature under constant humidity for 10 hours. Post exposure the mixed microbial cells were allowed to grow for 10 hours under biofilm stimulating conditions. Growth medium was discarded, and the adhered biomass was stained using CV. Percentage of biofilm formation is reported relative to glass (defined as 100%) and sterility control wells (defined as 0%). Error bars represent ±1 standard deviation (SD) from mean for three independent biological replicates (n=3). Statistical significance was assessed using one-way ANOVA; specific p-values of <0.05, <0.01, <0.001, and <0.0001 were indicated as *,**,***, and ****, respectively.*

***13. Biofilm biomass quantification of mixed Candida-bacterial species ($10^7$ CFU) in presence of pristine GO solution***

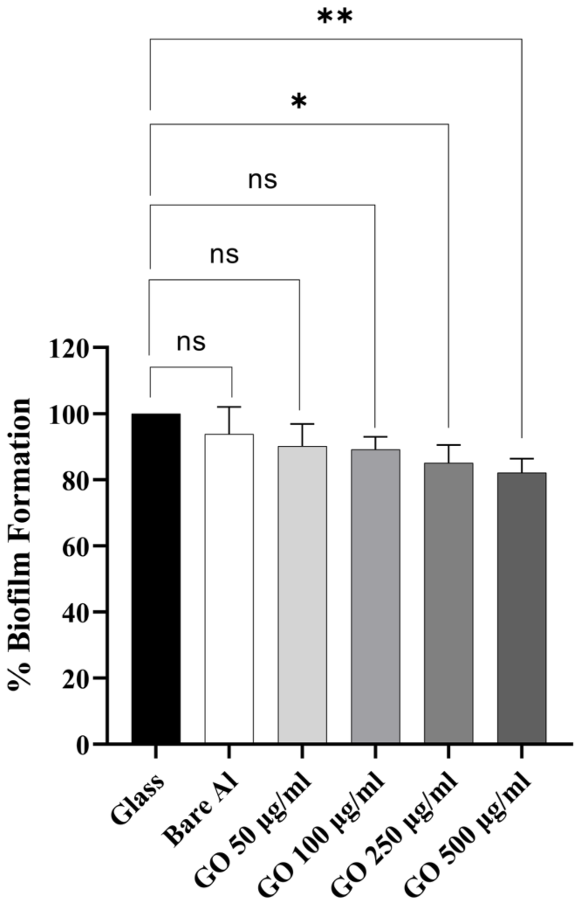

***Figure S13. Quantification of mixed species ($10^7$ CFU) biofilm biomass in presence of pristine GO after crystal violet (CV) staining.*** *Inhibition of biofilm formation was assessed by incubating the mixed microbial suspension in presence of pristine GO (50, 100, 250 , and 500 µg/ml) under optimal conditions for 10 hours. Post exposure the mixed microbial cells were allowed to grow for 10 h under biofilm stimulating conditions. Growth medium was discarded, and the adhered biomass was stained using CV. Percentage of biofilm formation is reported relative to glass (defined as 100%) and sterility control wells (defined as 0%). Error bars represent ±1 standard deviation (SD) from mean for three independent biological replicates (n=3). Statistical significance was assessed using one-way ANOVA; specific p-values of <0.05, <0.01, <0.001, and <0.0001 were indicated as *,**,***, and ****, respectively.*

***14. Pairwise comparison of antibiofilm efficiency between pristine GO solution vs Al_GO/P surfaces at $10^7$ CFU inoculum load***

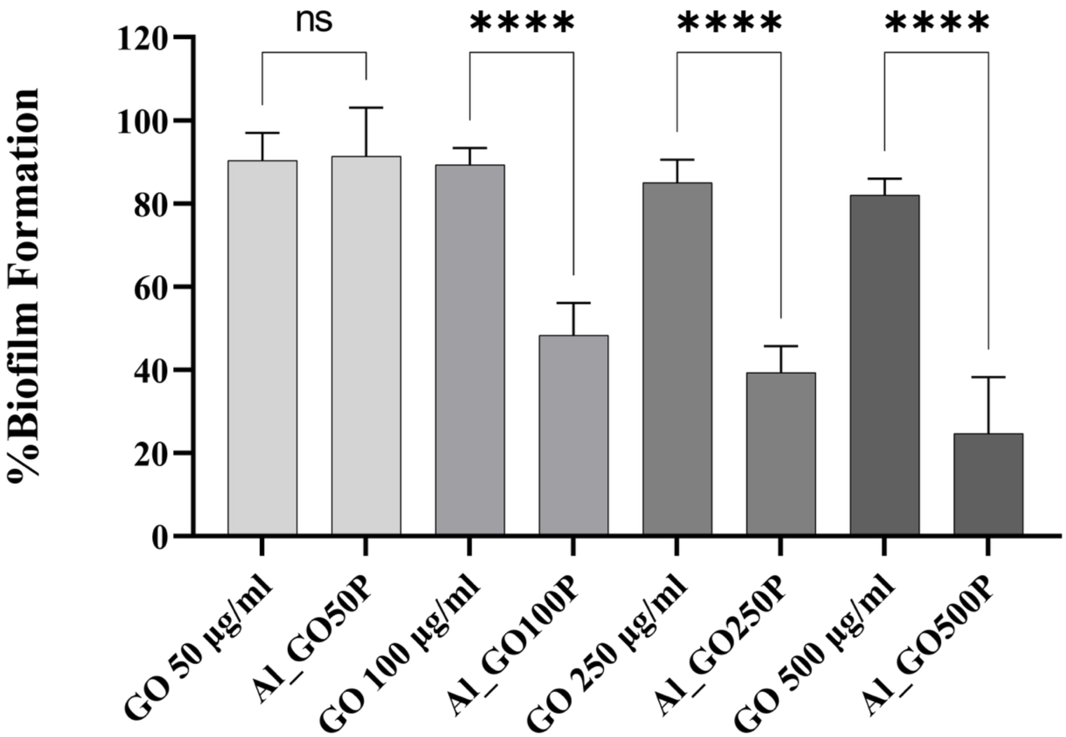

***Figure S14. Pairwise comparison of mixed species ($10^7$ CFU) biofilm biomass exposed to pristine GO solution vs. Al_GO/P substrates at corresponding concentrations after crystal violet (CV) staining.*** *Percentage of biofilm formation is reported relative to glass (defined as 100%) and sterility control wells (defined as 0%). Error bars represent ±1 standard deviation (SD) from mean for three independent biological replicates (n=3). Statistical significance was assessed using one-way ANOVA; specific p-values of <0.05, <0.01, <0.001, and <0.0001 were indicated as *,**,***, and ****, respectively.*

***15. Quantitative percentages of biofilm formation***

***Table S4. Quantitative values of percentage biofilm formation of mixed Candida-bacterial species at two distinct inoculum ($10^7$ and $10^8$ CFU) concentrations exposed to Al_GO/P substrates and pristine GO solution.***

| **Inoculum load** | **Surfaces** | **% Biofilm Formation** |
|---|---|---|
| **$10^7$ CFU** | *Glass* | *100.0 ± 0.0* |
| | *Bare Al* | *93.8 ± 8.2* |
| | *Al_GO50P* | *91.3 ± 11.8* |
| | *Al_GO100P* | *48.5 ± 7.7* |
| | *Al_GO250P* | *39.3 ± 6.5* |
| | *Al_GO500P* | *14.3 ± 5.4* |
| **$10^8$ CFU** | *Glass* | *100.0 ± 0.0* |
| | *Bare Al* | *95.7 ± 3.5* |
| | *Al_GO50P* | *94.5 ± 4.6* |
| | *Al_GO100P* | *48.7 ± 3.3* |
| | *Al_GO250P* | *40.3 ± 9.9* |
| | *Al_GO500P* | *19.0 ± 2.2* |
| **Inoculum load** | **Pristine GO** | **% Biofilm Formation** |
| **$10^7$ CFU** | *GO 50 µg/ml* | *90.2 ± 6.7* |
| | *GO100 µg/ml* | *89.2 ± 3.8* |
| | *GO 250 µg/ml* | *85.1 ± 5.4* |
| | *GO 500 µg/ml* | *82.1 ± 4.2* |

***16. Quantitative estimation of mean fluorescence intensity per cell corresponding to fluorescent images of ROS production using ImageJ software***

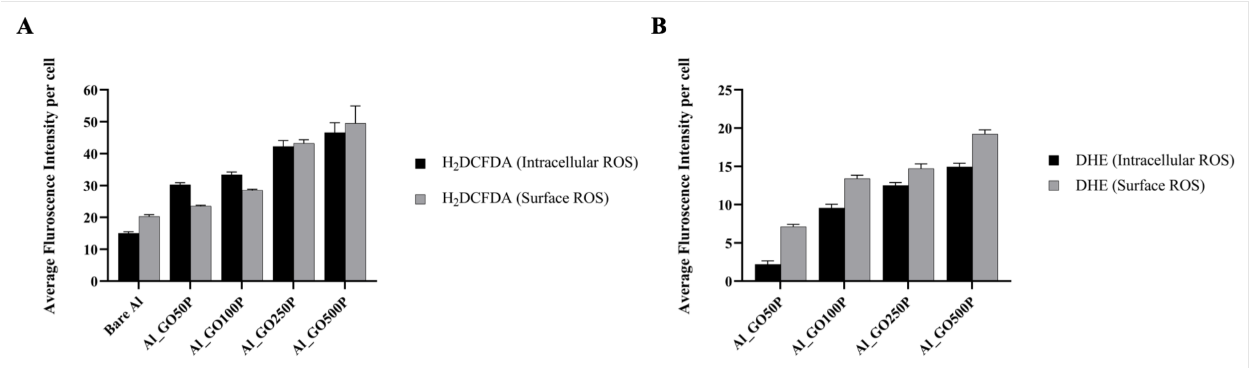

***Figure S15. Graphical representation of the average fluorescence intensity per cell*** *corresponding to fluorescence images of ROS production using* ***(A)*** *$H_2$DCFDA and* ***(B)*** *DHE fluorescent indicators. Error bars represent ±1 standard deviation (SD) from the mean of fluorescence quantifications performed using ImageJ software.*

### *17. ICP-MS analysis*

***Table S5. ICP-MS analysis to determine metal ion leaching in abiotic and biotic immersion environment.***

| ***Samples*** | ***Metal Ions of Interest*** | ***Concentration of metal ions detected by ICP-MS (Abiotic)*** | ***Concentration of metal ions detected by ICP-MS (Biotic)*** |
|---|---|---|---|
| ***Bare Al*** | $Al^{3+}$ | 0.82 ppm | 0.75 ppm |
| ***CR Steel*** | $Fe^{2+}$ | 3.00 ppm | 2.00 ppm |
| ***Al_GO50*** | $Al^{3+}$ | 0.02 ppm | 0.05 ppm |
| ***Al_GO100*** | $Al^{3+}$ | 0.26 ppm | 0.12 ppm |
| ***Al_GO250*** | $Al^{3+}$ | 0.88 ppm | 1.04 ppm |
| ***Al_GO500*** | $Al^{3+}$ | 1.30 ppm | 1.56 ppm |
| ***Al_GO50P*** | $Al^{3+}$ | 0.03 ppm | 0.01 ppm |
| ***Al_GO100P*** | $Al^{3+}$ | 0.09 ppm | 0.04 ppm |
| ***Al_GO250P*** | $Al^{3+}$ | 0.33 ppm | 0.45 ppm |
| ***Al_GO500P*** | $Al^{3+}$ | 0.43 ppm | 0.54 ppm |

#### *18. FESEM images of Al_GO and Al_GO/P substrates before and after corrosion under abiotic environment*

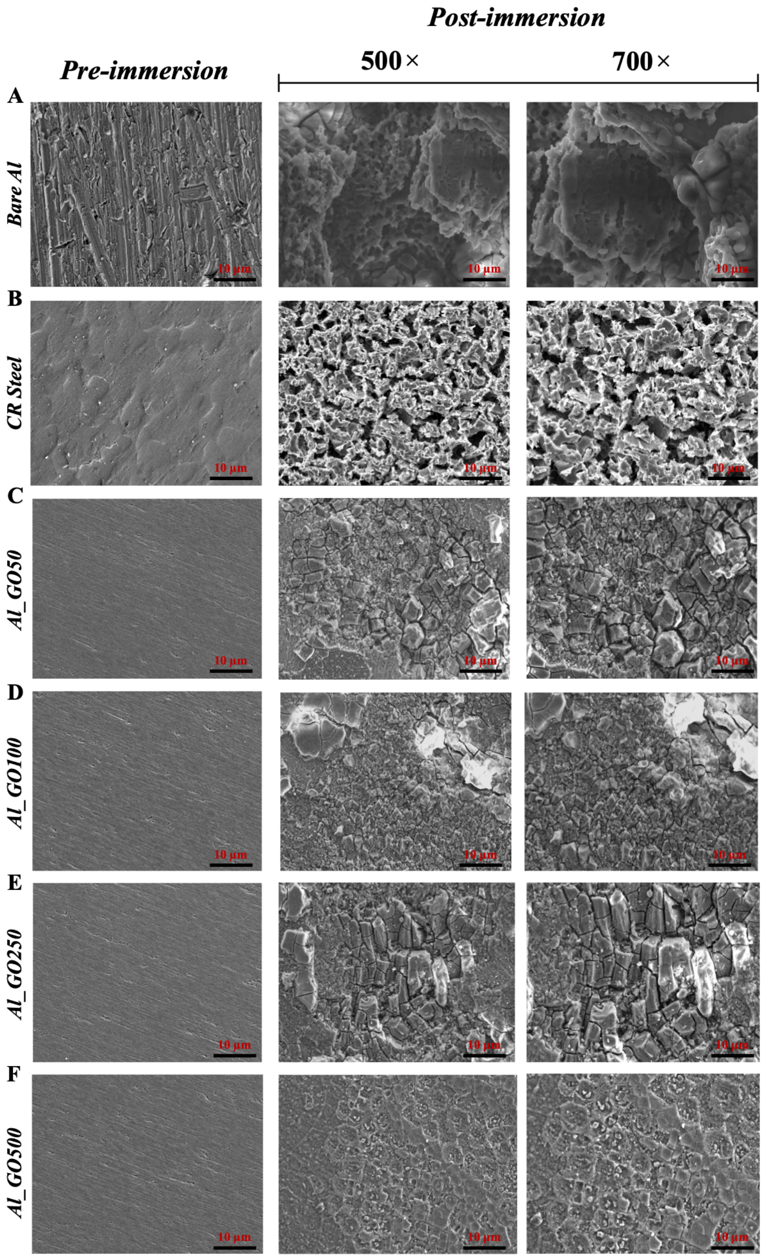

***Figure S16. FESEM analysis of control (bare Al, CR Steel, Al_GO50, Al_GO100, Al_GO250, and Al_GO500) substrates under abiotic immersion conditions.*** *FESEM micrographs of (**A**) bare Al, (**B**) CR Steel, and (**C–F**) synthesized Al_GO substrates following 45 days of exposure to abiotic environment, illustrating rough surface morphology at magnifications of 500× and 700×. Al_GO coatings exhibited corrosion features similar to that of Bare Al and CR Steel.*

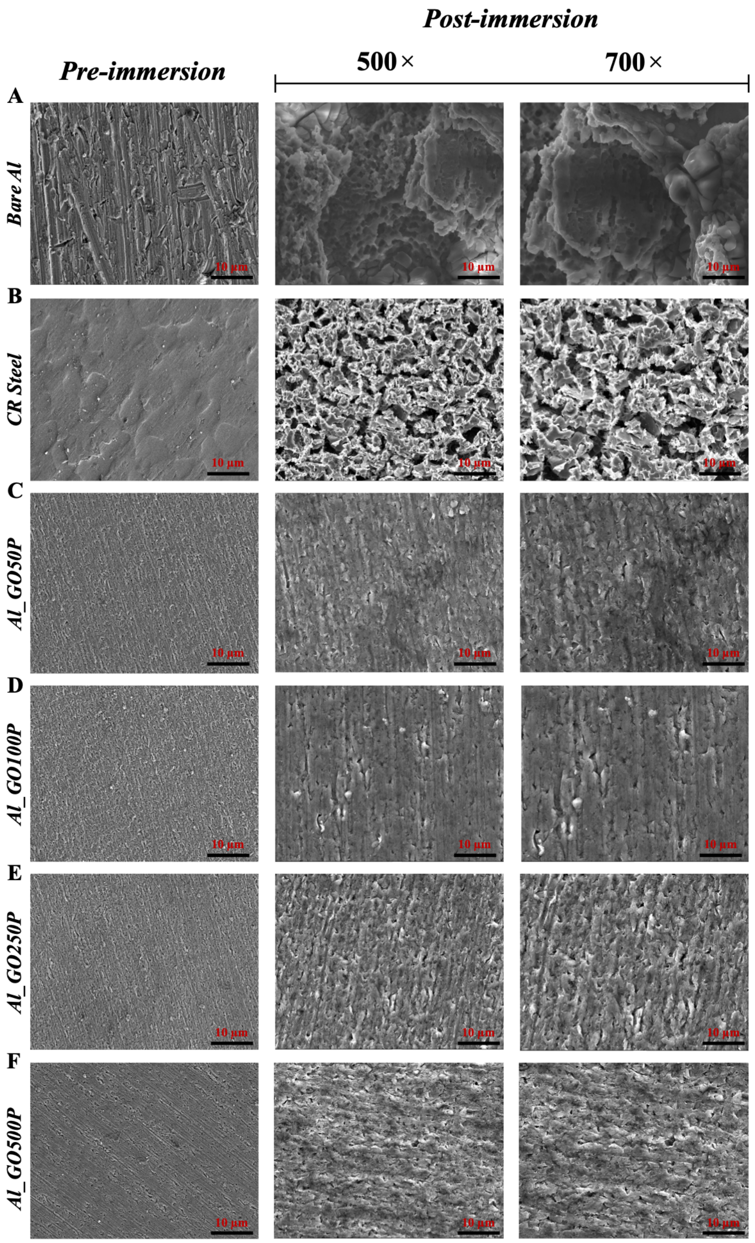

***Figure S17. FESEM analysis of control (bare Al and CR Steel) and test (Al_GO/P) substrates under abiotic immersion conditions.*** *FESEM micrographs of (**A**) bare Al, (**B**) CR Steel, and (**C–F**) synthesized Al_GO/P substrates following 45 days of exposure to abiotic environment, illustrating rough to smooth surface morphology at magnifications of 500× and 700×. Bare Al and CR Steel exhibited pitting and depth corrosion, whereas the Al_GO/P coatings maintained a comparatively uniform and intact surface, devoid of visible corrosion features.*

*19. FESEM images of Al_GO and Al_GO/P substrates before and after corrosion under biotic environment*

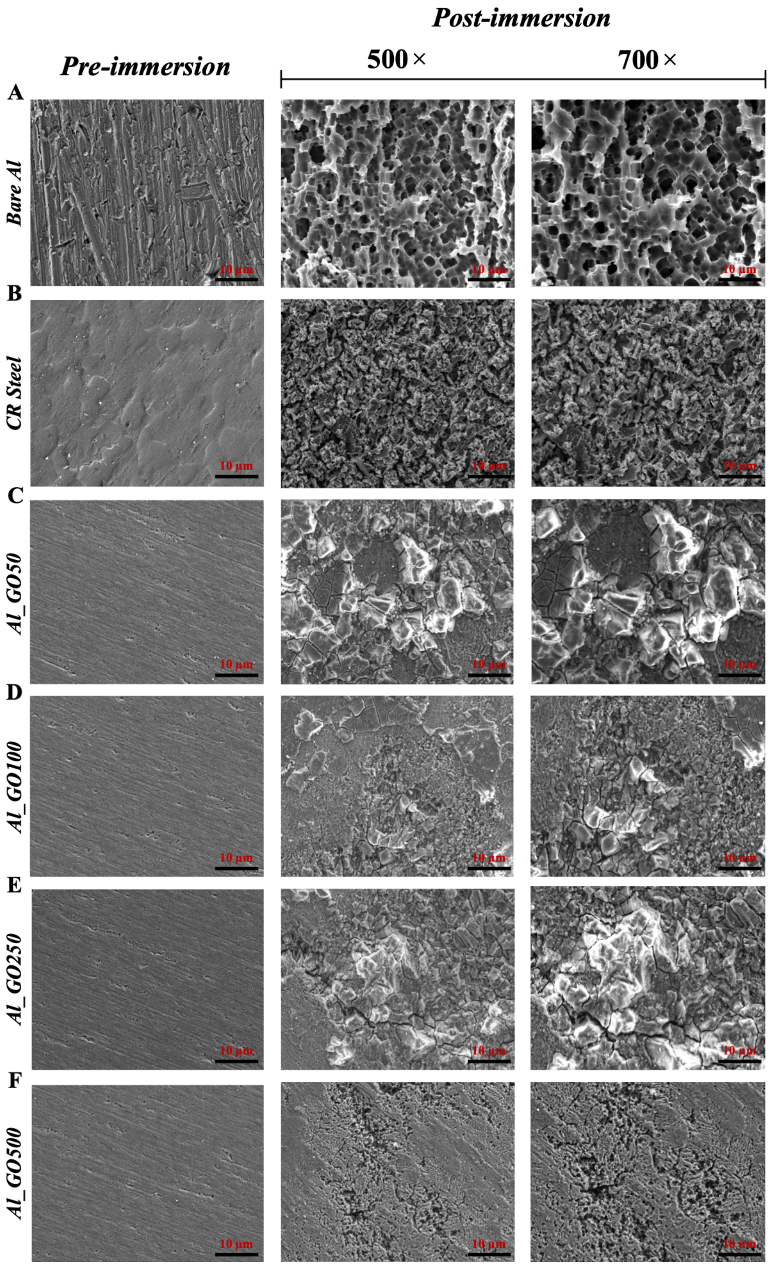

***Figure S18. FESEM analysis of control (bare Al, CR Steel, Al_GO50, Al_GO100, Al_GO250, and Al_GO500) substrates under biotic immersion conditions.*** *FESEM micrographs of (**A**) bare Al, (**B**) CR Steel, and (**C–F**) synthesized Al_GO substrates following 45 days of exposure to abiotic environment, illustrating rough surface morphology at magnifications of 500× and 700×. Al_GO coatings exhibited corrosion features similar to that of Bare Al and CR Steel.*

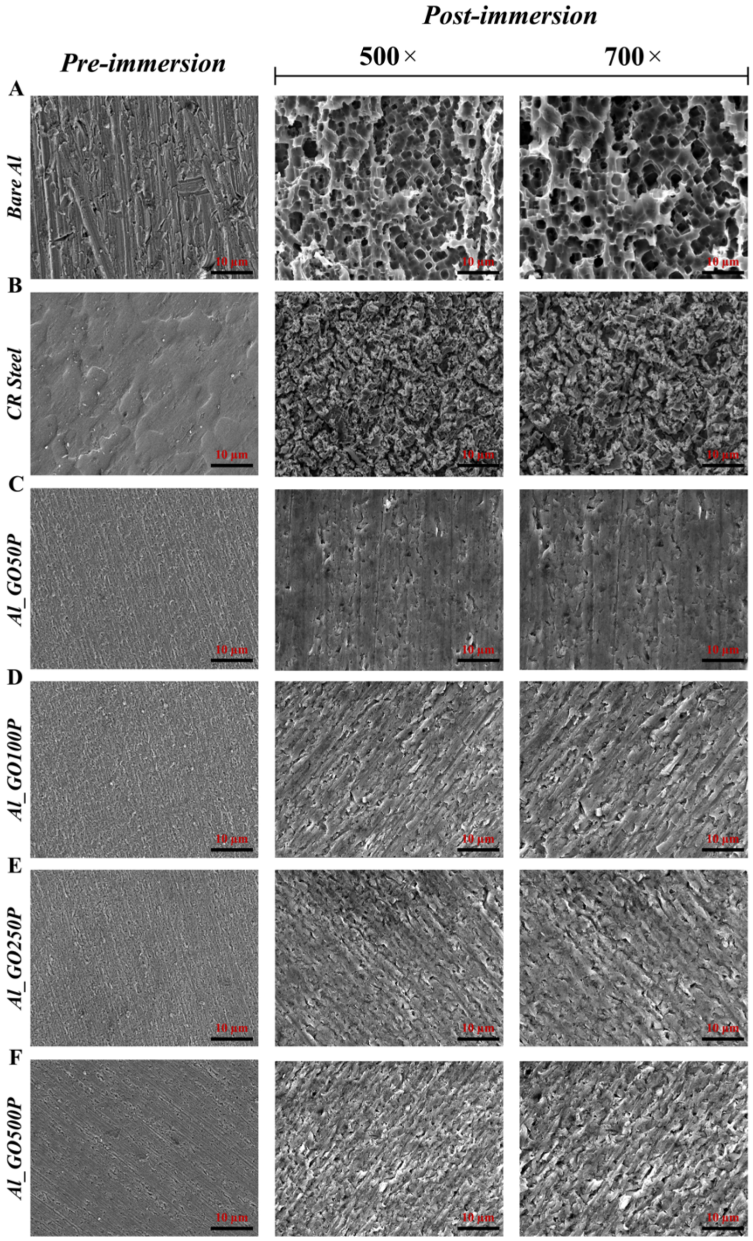

***Figure S19. FESEM analysis of control (bare Al and CR Steel) and test (Al_GO/P) substrates under biotic immersion conditions.*** *FESEM micrographs of (**A**) bare Al, (**B**) CR Steel, and (**C–F**) synthesized Al_GO/P substrates following 45 days of exposure to biotic environment, illustrating rough to smooth surface morphology at magnifications of 500× and 700×. The bare Al and CR Steel surfaces revealed extensive pitting and irregular corrosion patterns, whereas the Al_GO/P substrates preserved a smooth surface morphology with no visible signs of microbial-induced degradation.*

***20. Optical images of substrates before and after corrosion under abiotic and biotic environments***

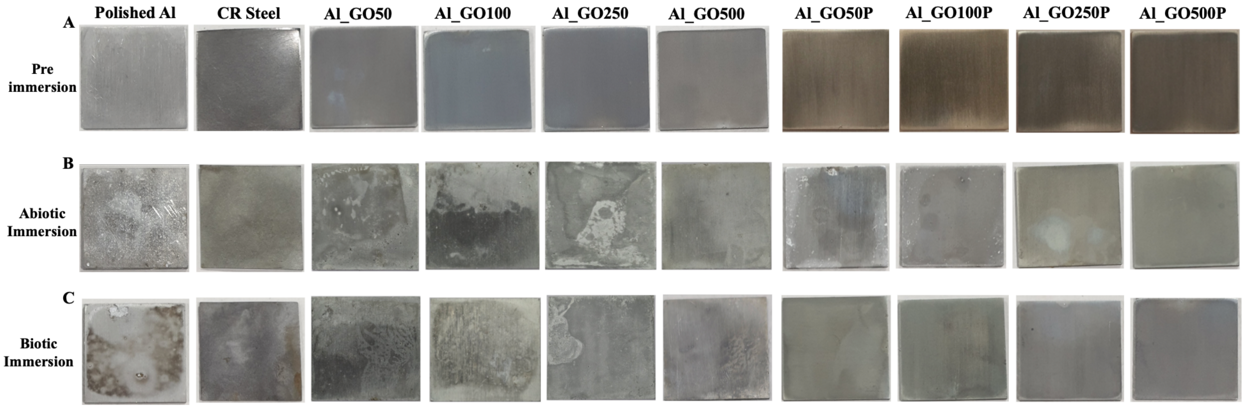

***Figure S20. Optical images of controls (bare Al, CR Steel, Al_GO substrates) and test Al_GO/P substrates.*** *(A) pre-immersion, (B) immersed for 45 days in abiotic conditions, and (C) immersed for 45 days under biotic conditions.*

***21. Cytotoxicity Assay: quantitative assessment of cell viability percentages***

***Table S6A.*** *HEK293T cells were exposed to 100% crude extracts of different Al_GO/P substrates for a period of 48 hours and MTT reduction was evaluated. Results are expressed as % of the control cells (HEK293T, 100%) not exposed to crude extracts. Values are the average of three independent biological replicates, expressed as mean ± SD.*

| ***Indirect (Extract) assay*** | | | | |
|---|---|---|---|---|
| ***Samples*** | ***24 hours*** | | ***48 hours*** | |
| | ***% Cell viability*** | ***P-value (*significance)*** | ***% Cell viability*** | ***P-value (*significance)*** |
| HEK293T cell control | 100 ± 0.0 | - | 100 ± 0.0 | - |
| Glass | 97.6 ± 3.8 | ns | 103.3 ± 6.3 | ns |
| Bare Al | 71.9 ± 6.9 | <0.0001 (****) | 72.6 ± 5.9 | <0.0001 (****) |
| Al_GO50 | 85.4 ± 3.3 | 0.0059 (**) | 72.5 ± 2.4 | <0.0001 (****) |
| Al_GO100 | 81.8 ± 2.7 | 0.0009 (***) | 65.9 ± 0.2 | <0.0001 (****) |
| Al_GO250 | 81.5 ± 3.6 | 0.0008 (***) | 52.7 ± 3.6 | <0.0001 (****) |
| Al_GO500 | 72.8 ± 6.6 | <0.0001 (****) | 40 ± 4.7 | <0.0001 (****) |
| Al_GO50P | 97.4 ± 5.3 | ns | 102.2 ± 7.0 | ns |
| Al_GO100P | 101 ± 1.7 | ns | 95.8 ± 4.6 | ns |
| Al_GO250P | 102 ± 1.1 | ns | 94.6 ± 3.3 | ns |
| Al_GO500P | 96.9 ± 2.2 | ns | 97.6 ± 4.5 | ns |
| Al_GO1000P | 95 ± 1.7 | ns | 94.6 ± 5.4 | ns |

***Table S6B.*** *HEK293T cells were exposed directly to the synthesized Al_GO/P substrates for a period of 48 hours and MTT reduction was evaluated. Results are expressed as % of the control cells (HEK293T, 100%) not exposed to Al_GO/P substrates. Values are the average of three independent biological replicates, expressed as mean ± SD.*

| ***Direct Contact Assay*** | | | | |
|---|---|---|---|---|
| | ***24 hours*** | | ***48 hours*** | |
| ***Samples*** | ***% Cell viability*** | ***P-value (*significance)*** | ***% Cell viability*** | ***P-value (*significance)*** |
| HEK293T cell control | 100 ± 0.0 | - | 100 ± 0.0 | - |
| Glass | 94.7 ± 1.7 | ns | 103.8 ± 6.9 | ns |
| Bare Al | 77.5 ± 5.2 | 0.0025 (**) | 53.0 ± 8.2 | <0.0001 (****) |
| Al_GO50 | 77.2 ± 7.0 | 0.0001 (***) | 65.4 ± 1.4 | <0.0001 (****) |
| Al_GO100 | 75.6 ± 6.8 | <0.0001 (****) | 64.5 ± 1.5 | <0.0001 (****) |
| Al_GO250 | 61.7 ± 1.3 | <0.0001 (****) | 56.2 ± 6.5 | <0.0001 (****) |
| Al_GO500 | 61 ± 3.8 | <0.0001 (****) | 49.9 ± 5.9 | <0.0001 (****) |
| Al_GO50P | 89.5 ± 8.2 | ns | 95.8 ± 2.8 | ns |
| Al_GO100P | 90.9 ± 10.5 | ns | 96.1 ± 4.4 | ns |
| Al_GO250P | 86.7 ± 6.6 | ns | 84.3 ± 3.7 | 0.0104 (*) |
| Al_GO500P | 82.5 ± 6.3 | 0.0183 (*) | 84.1 ± 6.1 | 0.0096 (*) |
| Al_GO1000P | 64.9 ± 4.4 | <0.0001 (****) | 59.1 ± 4.9 | <0.0001 (****) |

### 22. *Cytotoxicity Assay: quantitative assessment of cell viability percentages of HEK293T cells in presence of Al_GO substrates*

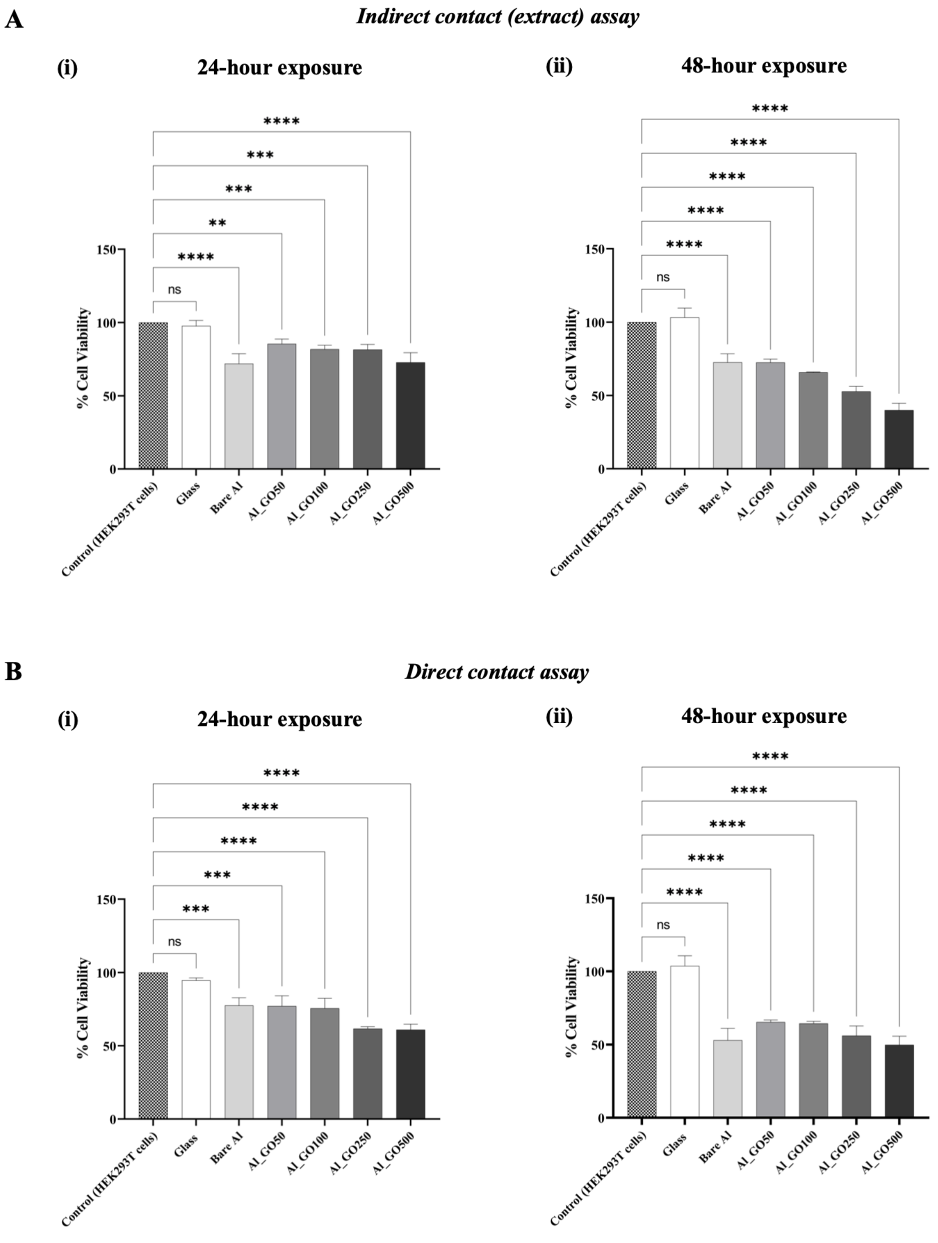

***Figure S21. Cytotoxicity evaluation of different control (Al_GO) substrates.*** *Comparison of cell viability, assessed by MTT method in the DMEM culture medium, after exposure to (**A**) Al_GO sample extracts (100%) and (**B**) after direct contact with the Al_GO samples at (**i**) 24 hours and (**ii**) 48 hours. Data are expressed as % values of result obtained for control cells not exposed to materials or material extracts. In the extract and direct test, all tested Al_GO substrates show*

*consistently lower cell viability relative to control cells over 48 hours. Error bars represent ±1 standard deviation (SD) from mean for three independent biological replicates (n=3). Statistical significance was assessed using one-way ANOVA; specific p-values of <0.05, <0.01, <0.001, and <0.0001 were indicated as *,**,***, and ****, respectively.*

#### *23. Cytotoxicity Assay: quantitative assessment of cell viability percentages of HEK293T cells in presence of pristine GO*

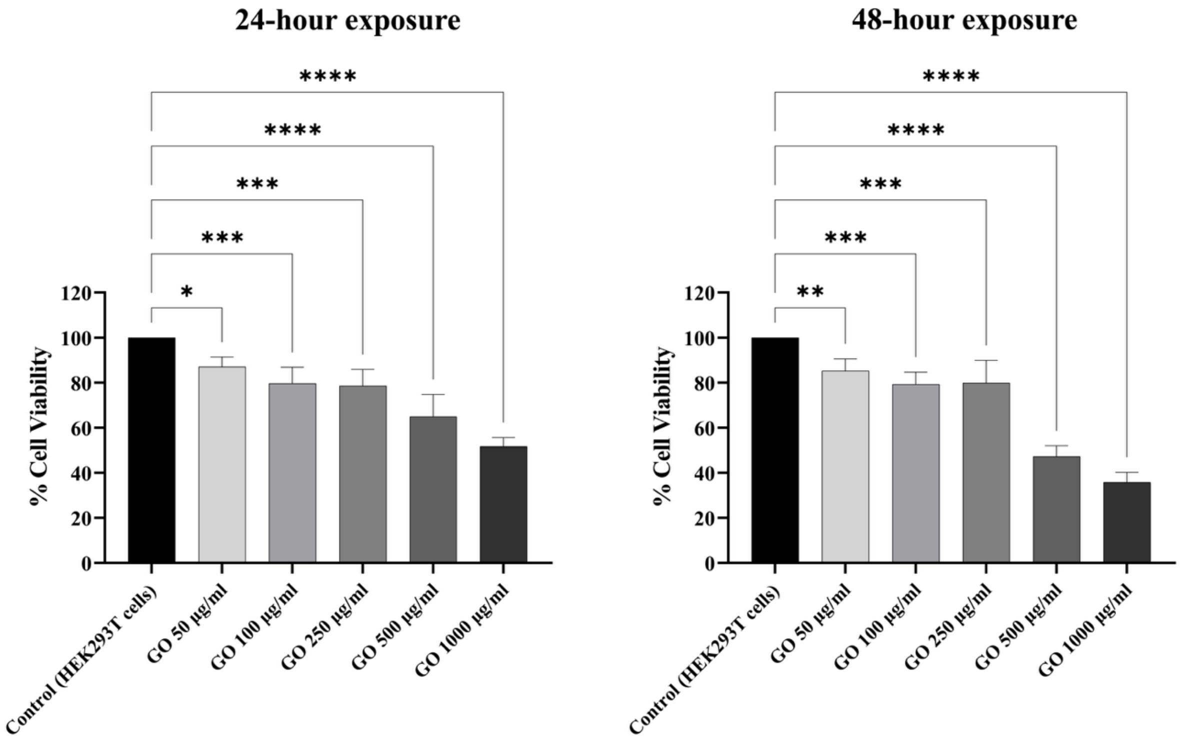

***Figure S22. Effect of pristine GO on cell viability of HEK293T cells.*** *Comparison of cell viability, assessed by MTT method in the DMEM culture medium, in presence of pristine GO at different concentrations (50, 100, 250, 500, and 1000 µg/ml) for 24 hours and 48 hours. Data are expressed as % values of result obtained for control cells (HEK293T) not exposed to pristine GO. Error bars represent ±1 standard deviation (SD) from mean for three independent biological replicates (n=3). Statistical significance was assessed using one-way ANOVA; specific p-values of <0.05, <0.01, <0.001, and <0.0001 were indicated as *,**,***, and ****, respectively.*

***24. Cytotoxicity Assay: quantitative assessment of cell viability percentages of HEK293T cells in presence of PEDOT:PSS***

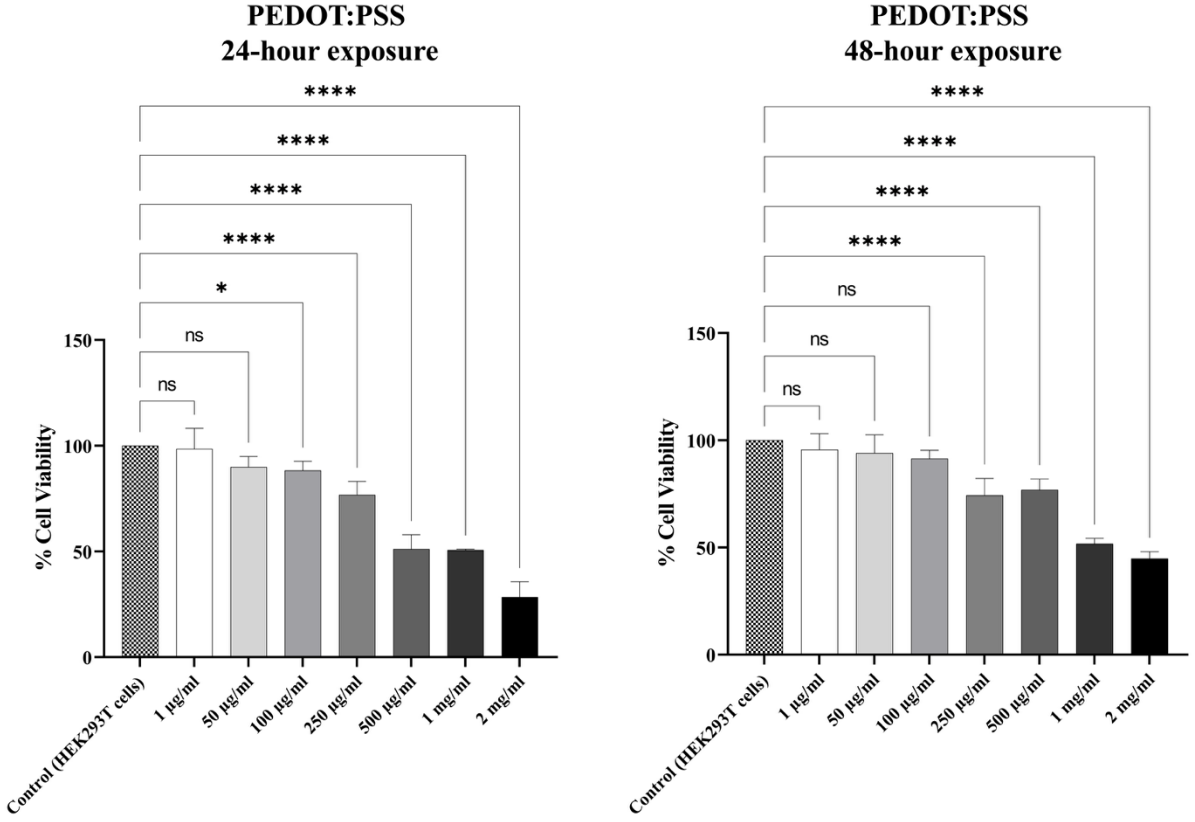

***Figure S23. Effect of PEDOT:PSS on cell viability of HEK293T cells.*** *Comparison of cell viability, assessed by MTT method in the DMEM culture medium, in presence of PEDOT:PSS at different concentrations (1 µg/ml, 50 µg/ml, 100 µg/ml, 250 µg/ml, 500 µg/ml, 1 mg/ml, and 2 mg/ml) for 24 hours and 48 hours. Data are expressed as % values of result obtained for control cells (HEK293T) not exposed to PEDOT:PSS. Error bars represent ±1 standard deviation (SD) from mean for three independent biological replicates (n=3). Statistical significance was assessed using one-way ANOVA; specific p-values of <0.05, <0.01, <0.001, and <0.0001 were indicated as *,**,***, and ****, respectively.*

***Reference***

*(1) Kong, M.; Garriga, M.; Reparaz, J. S.; Alonso, M. I. Advanced Optical Characterization of PEDOT:PSS by Combining Spectroscopic Ellipsometry and Raman Scattering. ACS Omega 2022, 7 (43), 39429–39436. https://doi.org/10.1021/ACSOMEGA.2C05945.*
